# Comparing the transmission blocking efficacy of Primaquine and Tafenoquine with *in vivo* pre-clinical models

**DOI:** 10.64898/2026.03.03.709295

**Authors:** Maëlle Duffey, Sara E. Zakutansky, Christin Gumpp, Michael J. Delves, Katarzyna A. Sala, Ellie Sherrard-Smith, Jake Baum, Didier Leroy, Matthias Rottmann, Andrew M. Blagborough

**Affiliations:** MMV Medicines for Malaria Venture, Route de Pré-Bois 20, PO Box 1826, 1215 Geneva 15, Switzerland; Department of Life Sciences, Sir Alexander Fleming Building, Imperial College London, Imperial College Road, South Kensington, London, SW7 2AZ, UK; University of Basel, 4001 Basel, Switzerland; London School of Hygiene and Tropical Medicine, Keppel Street, London, WC1E 7HT, UK; MRC Centre for Outbreak Analysis and Modelling, Department of Infectious Disease Epidemiology, Imperial College London, Norfolk Place, London, W2 1PG, UK; School of Biomedical Sciences, University of New South Wales (UNSW) Sydney, Kensington NSW 2052, Australia; Division of Microbiology and Parasitology, Department of Pathology, University of Cambridge, Tennis Court Road, Cambridge, CB2 1QP, UK

**Author notes:** equal contribution (Maëlle Duffey, Sara E. Zakutansky). current address: Global Antibiotic Research & Development Partnership (GARDP), 15 Chemin Camille-Vidart, 1202 Geneva, Switzerland (Maëlle Duffey).

## Abstract

The 8-aminoquinoline family are a well-established class of antimalarial drugs containing two clinically relevant analogues, primaquine and tafenoquine. These compounds have two therapeutically significant activities across the plasmodial lifecycle; elimination of *Plasmodium vivax* hypnozoites as part of a radical cure (relapse prevention), and as prophylactic transmission blocking compounds. Primaquine is currently recommended as a single low (0.25 mg/kg) dose administered with artemisinin-based combination therapy to reduce malarial transmission in areas at high risk of artemisinin partial resistance. Tafenoquine was approved in 2018 for radical cure when co-administered with chloroquine however, its transmission blocking efficacy in humans has not yet been fully evaluated, and direct head-to-head comparisons of transmission-blocking efficacy of primaquine and tafenoquine have not been performed.

Given that primaquine and tafenoquine are presumed to have similar mechanisms of action, tafenoquine may have potential use as a transmission blocking intervention. Furthermore, tafenoquine has a substantially longer half-life than primaquine, which could provide a pharmacokinetic advantage if such efficacy is confirmed. However, assessment of 8-aminoquinoline efficacy against the sexual stages of *Plasmodium* is complicated by the requirement for metabolic activation to generate parasite-reactive species, limiting the utility of *in vitro* assays.

Here, we directly compare the transmission blocking effects of primaquine and tafenoquine using two *in vivo* preclinical models. We titrate the transmission blocking efficacy of each compound, and evaluate the pharmacokinetics of each compound over time, linking drug exposure to efficacy. The evidence presented here suggests that beyond 24 hours, a single dose of tafenoquine is likely to have more clinically desirable pharmacokinetics, resulting in higher transmission blocking efficacy than primaquine. These findings are observed across multiple models, both in the absence or presence of a partner schizonticide and thus demonstrate a potential advantage of the utilisation of tafenoquine when compared to primaquine.

## Introduction

Malaria remains a major global health burden, with an estimated 282 million cases and 610,000 deaths annually (1). Although the introduction of effective antimalarial combination therapies has resulted in an approximately two-fold reduction in mortality over the past decade, eradication efforts remain fragile and incomplete. Sustained progress will require effective strategies to inhibit or interrupt parasitic transmission (2). Assessment of transmission-blocking interventions in the field is technically and conceptually challenging, encompassing potential variations in gametocytemia, screening methods, sample sizes, and mosquito-related metrics such as entomological inoculation rate (EIR) and sporozoite positivity. Previous studies have highlighted difficulties in identifying appropriate efficacy endpoints and target populations. For example, a community-based intervention in Sapone, Burkina Faso demonstrated that screening asymptomatic gametocyte carriers using rapid diagnostic tests and microscopy failed to identify a substantial proportion of still infectious individuals, who subsequently sustained community-level transmission in the following season (*3*). These findings underscore the need for more sensitive approaches to assess transmission-blocking efficacy, either through direct measurement of viable sexual-stage parasites or indirect monitoring of reductions in blood-stage infection incidence. The complexity and cost of such trials have limited their implementation, creating a major bottleneck in the development and evaluation of transmission-blocking antimalarials.

Historically, transmission blocking activity in anti-malarial drug discovery has been assessed primarily using the standard membrane feeding assay (SMFA), in which mosquitoes are fed human blood containing cultured *Plasmodium falciparum* stage V gametocytes in the presence or absence of test compounds (*4*). Infection prevalence and oocyst intensity are quantified following mosquito dissection 10–12 days later. While the SMFA provides robust measurements of gametocyte-to-oocyst inhibition, it captures only a limited portion of the transmission cycle (from stage V gametocyte to oocyst only) and is entirely *ex vivo*, precluding integration of pharmacokinetic parameters that are critical for translational relevance. Consequently, the development of *in vivo* preclinical models is important for the examination and triage of transmission-blocking compounds, considering the challenging long-term objective of conducting specific clinical trials with an anti-transmission efficacy end point as a goal. A relevant murine model that encompasses the entire parasite lifecycle *in vivo*, and specifically, transmission of malaria from infected mouse population to naïve mouse population was first developed with the aim of testing transmission blocking vaccine candidates, and subsequently was revealed to be a promising tool to test the transmission blocking potential of antimalarials like atovaquone (5) or compounds within the development pipeline having gametocytocidal or sporontocidal potency such as MMV390048 (*6*). More recently, the generation of *P. falciparum* stage V gametocytes and their use in humanized NSG mice engrafted with human erythrocytes has expanded the utility of murine models for studying transmission with human malaria parasites (*7*).

Several antimalarials in clinical development, including cipargamin (*8*), ganaplacide (*9*), and MMV390048 (*6*), have demonstrated potent transmission-blocking activity in the SMFA. In contrast, 8-aminoquinolines require metabolic activation and are not amenable to SMFA, despite their clinical potential as transmission-blocking agents (*10–13*). This drug class originated with the discovery of pamaquine in 1926, followed by the development of primaquine (PQ) and tafenoquine (TQ). PQ was identified as a highly effective compound with activity against *P. vivax* hypnozoites and was widely adopted for clinical use (*14–16*). PQ exhibits activity against *P. vivax* hypnozoites, liver and blood stages, and crucially, as a transmission blocking drug with efficacy against *P. falciparum* gametocytes (*17*). Its efficacy depends on metabolic activation, with strong evidence linking potency to CYP2D6-dependent and independent hydroxylated metabolites (*18–20*). This requirement for metabolism has historically limited the *in vitro* assessment of PQ activity against both hypnozoites and sexual stages.

The World Health Organization recommends the use of a single low dose of primaquine as a gametocytocide to help reduce malaria transmission and support elimination efforts in areas with resistance (*21*). Currently, administration of a single 0.25 mg/kg PQ dose is recommended in all patients with parasitologically-confirmed *P. falciparum* malaria on the first day of treatment, in addition to ACT in areas of partial artemisinin resistance (*22*). Use of single low dose (0.25 mg/kg) primaquine in combination with artemether-lumefantrine (AL) has been shown to significantly reduce gametocyte prevalence and infectivity to mosquitoes in randomized, double blind, placebo-controlled field trials in sub-Saharan Africa (*18*). Negative issues related to PQ usage include a prolonged treatment regime for radical cure (prevention of relapse) (7-14 days), increased trends of failing treatment outcomes (particularly at 15 mg/d regimen) (*23*), short half-life (∼6 hours (*24*)), and safety concerns related to the ability of PQ (and 8-aminoquinoline) to induce hemolysis in glucose-6-phosphate deficient (G6PD) individuals (*25*). Nonetheless, the dual hypnozoitocidal and gametocytocidal activity of PQ highlights the value of antimalarials that target multiple lifecycle stages, particularly in the context of emerging resistance to artemisinin and partner drugs (*26, 27*).

Tafenoquine (TQ) is a 5-phenoxy derivative of PQ and has recently been adopted in Brazil and Thailand as a therapy to prevent relapsing *P. vivax* infection following its development by WRAIR, GlaxoSmithKline and Medicines for Malaria Venture (*28*). Its extended half-life of 14 days (*29*), compared to ∼6 hours with PQ (*24*) facilitates single dose radical cure and prophylactic use when co-administered with chloroquine, resulting in increased tolerability and compliance compared to DHA-piperaquine (*30*). Phase I-III studies have also shown that TQ is a well-tolerated and effective oral chemoprophylactic agent for the treatment of *Plasmodium* (*31–33*). Anti-hypnozoite activity as part of a potential *P. vivax* radical cure (prevention of relapse) was demonstrated in Phase II and III trials when used in combination with other antimalarials such as chloroquine (*34*). Early animal studies have also suggested that TQ has greater efficacy than PQ against *P. cynomolgi* hypnozoites and comparable activity against *P. knowlesi* liver stages in non-human primates (*35*).

The anti-hypnozoite efficacy of TQ has been well demonstrated in multiple animal, pre-clinical, and clinical studies. Comparatively, the assessment of the ability of TQ to block transmission has lagged. Sporontocidal activity has been demonstrated in multiple models, with partial inhibition of *P. vivax* development in *Anopheles dirus* and stronger effects against *P. berghei* in *Anopheles stephensi* (*36, 37*). Direct examination of the gametocytocidal activity of TQ is sparse, with a single study demonstrating significant efficacy against *P. gallinaceum* at high doses (*38*). More recently, a phase II clinical trial showed that TQ administered with dihydroartemisinin–piperaquine accelerated gametocyte clearance and significantly reduced transmission at day 7 across all doses tested (0.42–1.66 mg/kg) (*39*). A retrospective analysis of six clinical trials in Mali further demonstrated that both PQ and TQ enhance the transmission-blocking effects of ACT, although TQ exhibited delayed activity relative to PQ (up to day 14) (*40*). A recent experimental human challenge study has demonstrated that single dose (50mg) TQ can significantly reduce transmission of *P. falciparum* to mosquitoes (*41*). Despite these advances, a direct, parallel head-to-head comparison of PQ and TQ transmission-blocking efficacy has not been performed.

Here, we use two recently developed and validated *in vivo* preclinical models (Fig. 1) to directly compare the transmission blocking efficacy of PQ and TQ. We quantify dose-dependent effects of drugs on circulating gametocytes and mosquito infection using multiple murine models, and integrate pharmacokinetic data to clarify exposure-efficacy relationships. These data provide a basis to evaluate the relative utility of PQ and TQ as transmission-blocking agents and inform their potential utilisation within current antimalarial treatment regimens.

**Figure 1:**
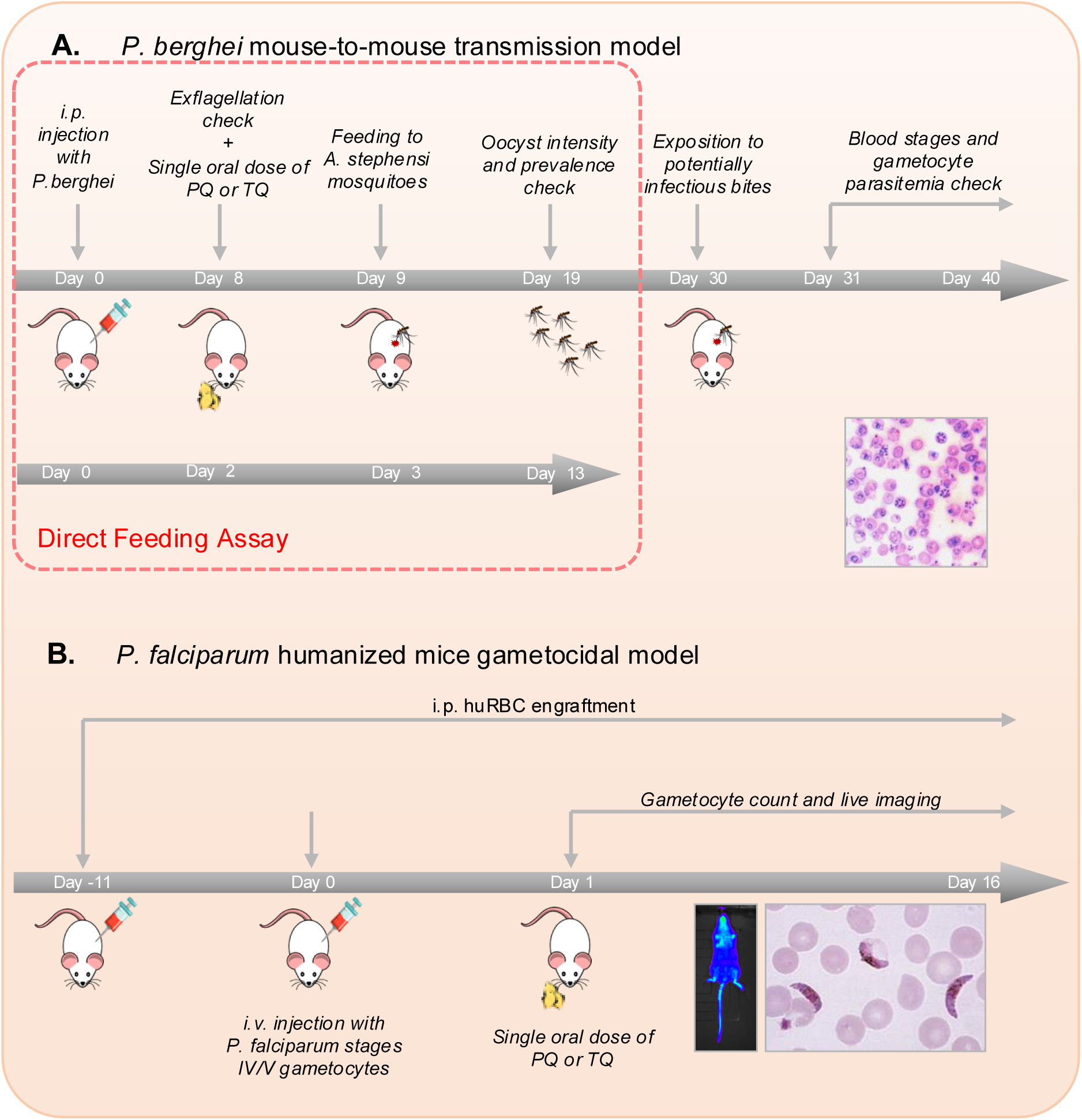
Graphical representation of the two *in vivo* experimental designs used in this study. (A) Mouse-to-mouse *P. berghei* and *A. stephensi* transmission model (adapted from Upton *et al*., 2015), and Direct Feeding Assay. Blood stage *P. berghei ANKA 2.34 i.p*. infected mice (5 mice per group) were treated with a single oral dose of Primaquine (PQ) or Tafenoquine (TQ) 8 days post-infection, after a visual exflagellation check. 500 female *A. stephensi* were allowed to feed on these mice 24h later (day 9 post-infection). Ten days later (day 19 post-infection), oocyst intensity and prevalence were determined. The Direct Feeding Assay (DFA) (performed with phenylhydrazine treated mice), is highlighted within the orange dotted line, and ends with the examination of oocyst intensity and prevalence. For continuation within the mouse-to-mouse model, twenty-one days after blood feeding, when sporozoites are estimated to be most infectious, new groups of 5 naïve mice were bitten by either 2, 5 or 10 mosquitoes. Salivary glands were dissected post bite to determine sporozoite intensity and prevalence. Mice exposed to mosquito bite were monitored for up to 10 days post bite (up to day 40 post-infection) for blood stage parasitemia (asexual stages and gametocytes) and time to patency. (B) *P. falciparum* humanized mice gametocidal model. NGS mice were *i.p.* engrafted with human erythrocytes 11 days prior the beginning of the experiment, and at regular interval until the end of the study. These humanized mice were *i.v.* infected with highly enriched *in vitro*-induced *P. falciparum* NF54/iGP_RE9H gametocyte in human erythrocytes (*7*) and treated with a single oral dose of PQ or TQ on the next day (day 1 post-infection). The gametocytemia was monitored by live imaging (bioluminescence) and microscopy (gametocyte count) up to day 16 post-infection.

## Results

### Titration and dose selection of primaquine for use in *P. berghei* transmission blocking assays

To enable a direct comparison of the transmission blocking efficacy between PQ and TQ, a primaquine dose of 50% reduction in oocyst prevalence (ED_50_) was selected using a direct feeding assay (DFA). This dose was selected to allow any significant differences in efficacy (either positive or negative) between both compounds to be detected. Single oral doses of PQ, ranging from 0.1 mg/kg to 12mg/kg were administered to *P. berghei* ANKA 2.34 infected mice, and the subsequent effect on transmission assessed in a direct feeding assay as described in Figure 1A. Blood feeding was performed 24-hours post administration of drug. Mosquitoes were dissected 10 days post bloodmeal, with mean numbers of oocysts (oocyst intensity) and number of infected mosquitoes (infection prevalence) determined, and transmission blocking efficacy from mice to mosquitoes expressed in comparison with negative control (1% methyl cellulose) treated mice.

Titration of the transmission blocking efficacy of PQ in the direct feeding assay is predictable and robust; inhibition in infection prevalence is described in Figure 2A and shows a sigmoidal dose response shape, resulting in an ED_50_ value of 3.35mg/kg (95% Cls; 3.25-3.46). The corresponding dose of TQ for further use in this study, accounting for molar differences, was calculated at 3.41 mg/kg. Similarly, the inhibition of infection *intensity* is characterized by a sigmoidal dose response curve (Fig. 2B), with a maximal inhibition reached at a dose of 4mg/kg and an ED_50_ value of 1.26 mg/kg. The data used to build both graphs in Fig. 2 are listed in Table S1.

**Figure 2:**
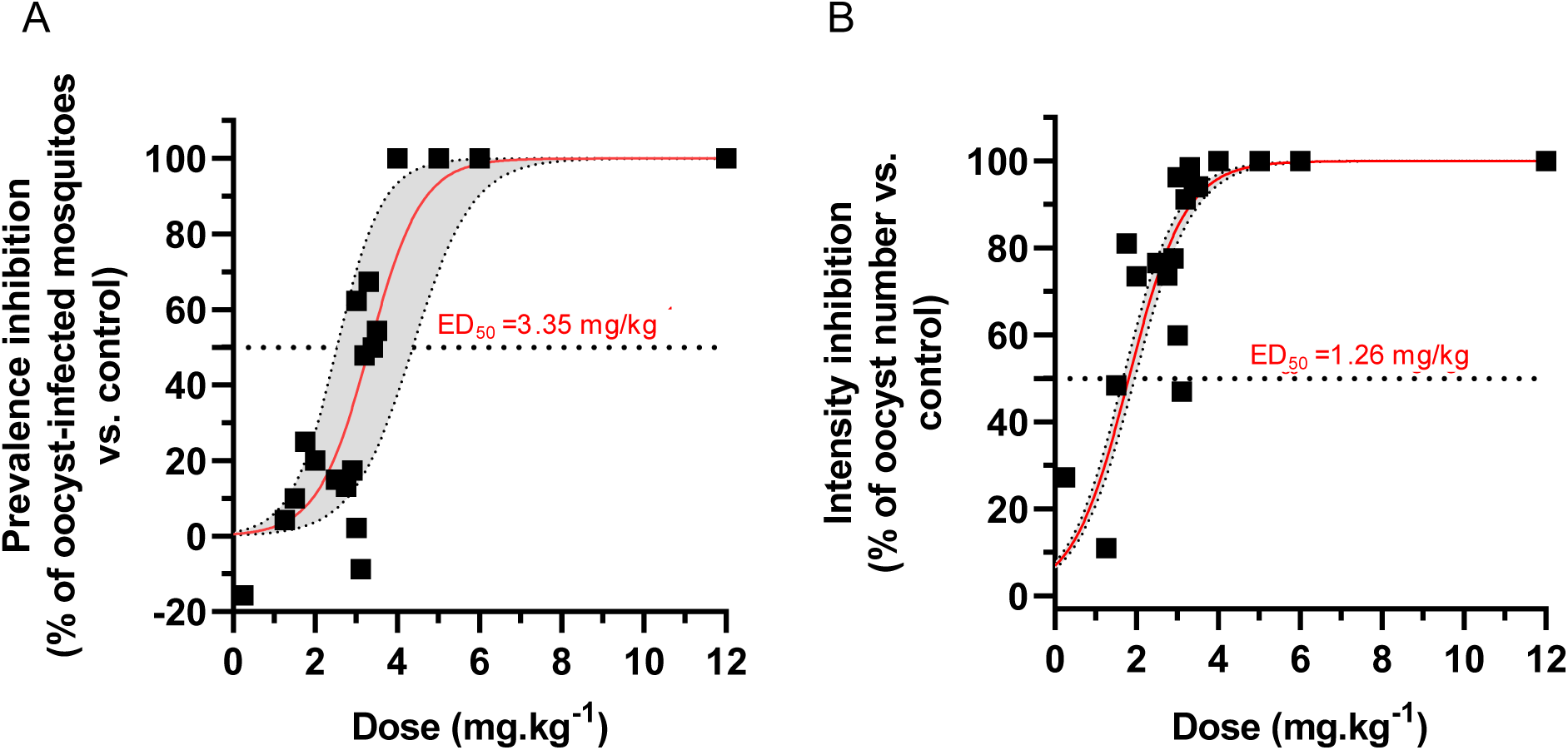
Dose response of transmission-blocking efficacy of primaquine administered orally to *P. berghei*-infected mice. Oocyst prevalence (A) and intensity (B) were determined following repeated direct feeding assays and ED_50_ values calculated. Each data point represents an individual mosquito feed, over 7 independent experiments. Data were fitted with a logistic function (red line and lower and upper limits greyed-out). The dotted line represents 50% of the maximum effect (ED_50_).

### Tafenoquine results in more sustained transmission blocking efficacy than primaquine

The transmission blocking efficacy of PQ and TQ, at identical doses (set at the ED_50_ of PQ against infection prevalence), was directly compared in parallel within a mouse-to-mosquito-to-mouse (M2M2M) model (*42,43*) (Fig. 1) to determine the impact of treatment on both mosquito and vertebrate populations at multiple transmission intensities (2, 5 or 10 mosquito bites per naive mouse). The use of multiple mosquito biting rates within this model allows us to estimate the effect size of this intervention, which is a measure of the ability of the treatment to reduce the basic reproductive number (*R*_o_) (*5,42,43*). The results of head-to-head comparison between primaquine at 3.35 mg/kg and tafenoquine at 3.41 mg/kg within this model are outlined in Table 1, with all raw data in table S4. Briefly, both PQ and TQ exhibited statistically significant transmission blocking efficacy 24 hours post drug treatment in comparison with no drug controls (S4). At the mosquito level, PQ resulted in a superior reduction in intensity and prevalence (42.8% and 28% respectively) when compared to TQ (10.9% reduction in intensity, 17.5% reduction in prevalence). When subsequent transmission to naive vertebrate populations was examined at different transmission intensities (2, 5 or 10 bites), this modest effect on transmission to the mosquito translated into a lack of inhibition in naive mice, and logical, but relatively low effect sizes of 27.1% for PQ and 15.4% for TQ. Under the conditions examined within this assay (single drug dosage, bloodmeal 24 hours post-treatment), TQ was less efficacious than PQ at the equivalent dose.

**Table 1:** Comparative transmission blocking efficacy of primaquine and tafenoquine in the *P. berghei* mouse-to-mouse transmission model at equal molar concentrations. Primaquine was dosed at a concentration of 3.35 mg.kg^-1^ and tafenoquine at 3.41 mg.kg^-1^. Oocyst prevalence and intensity were measured in the infected mosquitoes on day 10 post-bloodmeal. The infection of naïve mice was evaluated over the 10 days following challenge with mosquito bite at three individual biting rates (2, 5 and 10), and the inhibition of blood stage parasitemia at the end of the experiment (day 10 post-bite). The effect size was calculated using a chain binomial model (described in (*6,42,43*)). 95% confidence intervals are displayed. Mean of 2 independent mouse-to-mouse experiments, are presented.

|  |  | Primaquine | Tafenoquine |
| --- | --- | --- | --- |
| *Transmission blocking efficacy to the mosquito host* | Inhibition in oocyst intensity (%) | 42.8 | 10.9 |
|  | Inhibition in prevalence (%) | 28.0 | 17.5 |
| *Transmission blocking efficacy to the mouse host* | Inhibition in infection (%) | -21.4 [-78.5 - 11.1] | -36.84 [-93.8 – 0.00] |
|  | Inhibition of parasitemia (%) | 4.72 [-38.7 - 36.2] | 4.92 [-23.7 - 28.4] |
|  | Effect size (%) | 27.1 [11.3 - 42.3] | 15.4 [0.00 - 32.0] |

To examine this observation further, the transmission blocking efficacy of both PQ and TQ at ED_50_ doses (3.35 mg/kg and 3.41 mg/kg respectively) was measured over time in triplicate direct feeding assays at 24-, 48-, 72– and 96-hours post-dosage (Fig. 3). Impact on oocyst intensity (A), prevalence (B), asexual parasitemia (C) and gametocytemia (D) were measured in parallel. Twenty-four hours post drug-treatment, PQ exhibits statistically significant superior transmission blocking efficacy when compared to tafenoquine (Figs. 3A and B). Specifically, PQ results in an 88.37% reduction in oocyst intensity and 54% reduction in oocyst prevalence, compared to a 29.82% and 13% reduction in intensity and prevalence with TQ, respectively. This trend matches the data observed in the M2M2M assay (Table 1), where PQ outperforms TQ 24 hours post drug treatment. From 42 hours onwards, this phenomenon is reversed, with TQ consistently achieving higher transmission blocking efficacy, with maximum reductions observed at 48-hours post treatment (96.28% and 75% reductions in intensity and prevalence respectively). High rates of inhibition persist for TQ until the last time point examined (96hrs), with statistically supported significance, whereas more modest inhibition is observed following treatment with PQ at later time points. Post-exposure with both compounds, asexual parasitemia marginally decreases 24-48 hours post treatment, and then rises rapidly over the subsequent days (Fig. 3C), Conversely, gametocytemia is reduced post-drug treatment with either compound (Fig. 3D). These reductions in circulating parasitic lifecycle stages are consistent with poor activity and modest clearance of asexual blood stage parasites, and partial transmission blocking efficacy.

**Figure 3:**
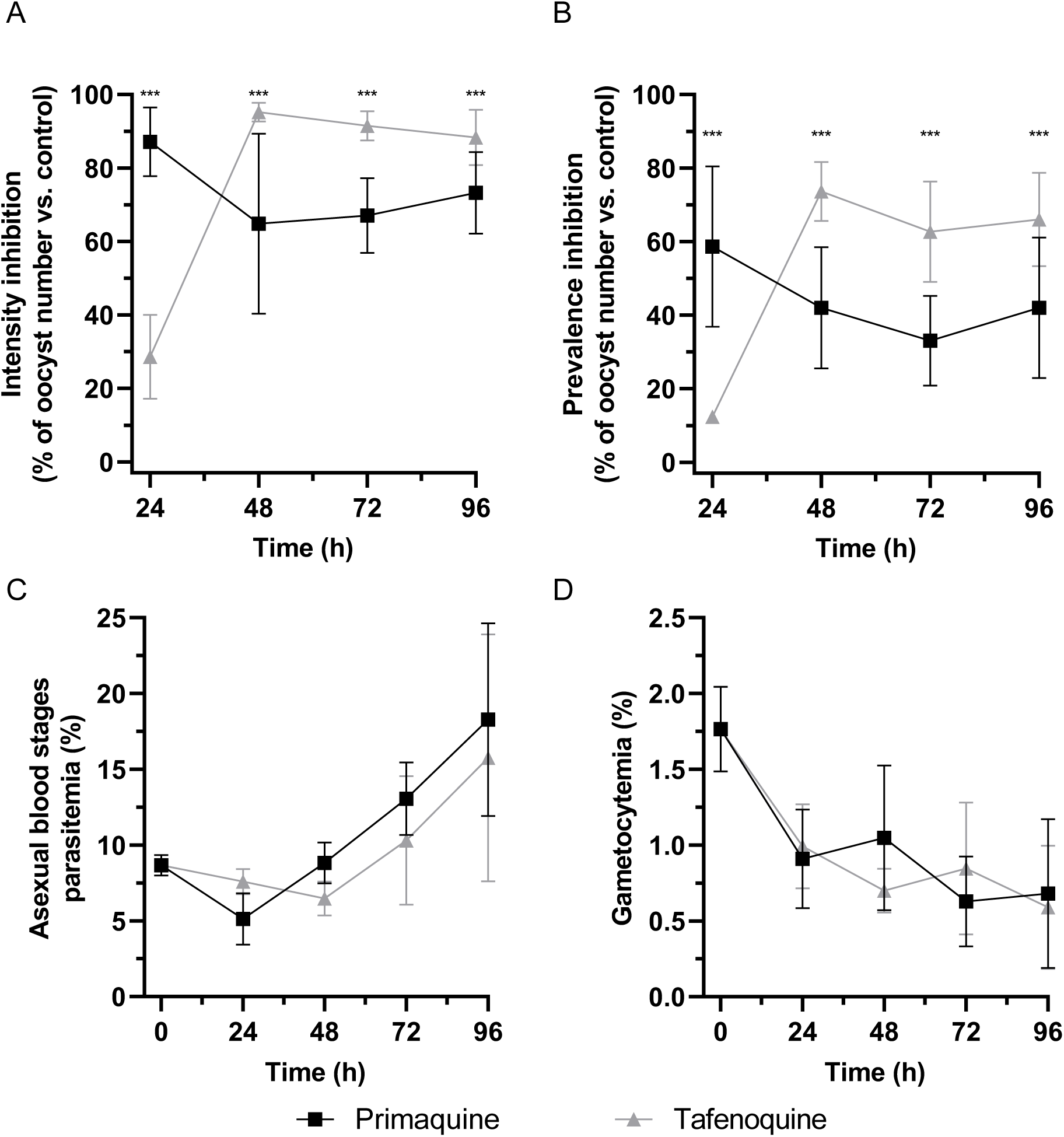
Primaquine and tafenoquine efficacies in the *P. berghei* mouse-to-mouse transmission model at equal molar concentrations. (A and B) Transmission blocking efficacy in a Direct Feeding Assay (DFA) on *P. berghei*-infected mice orally treated with single doses of PQ (dosed at 3.35 mg.kg^-1^) and TQ (dosed at 3.41 mg.kg^-1^) and fed to mosquitoes at 24, 48, 72 and 96 hours post drug administration. Oocyst intensity (A) and prevalence (B) were then determined 10 days after feeding (day 19 post-infection) and compared to carrier control-treated mice. The difference in intensity (Mann-Whitney test) and in prevalence (Fisher’s exact test) between PQ and TQ treatment was significant at each measured time point (p < 0.0001, ***). (C and (D) Efficacy of PQ and TQ at reducing asexual (C) and sexual (D) blood stages. On day 30 post-infection, mice were expose to potentially infectious bites and blood smears were taken for up to 10 days post-bite (day 40 post-infection) to quantify blood stages and gametocytes. No significant differences were observed between PQ and TQ regarding asexual and gametocyte parasitemias (time-points t-test comparisons). Mean +/− SEM of 3 independent experiments.

To explain the respective reductions in parasite levels in further detail, the concentration of PQ and TQ in the blood plasma of each experimentally drug treated mouse was measured by LC-MS at the time of mosquito feeding (Fig. S1, table S3). Within this analysis, PQ was not detected in any samples collected from mice following administration of a single oral dose at 3.35 mg/kg. We conclude that PQ is cleared within 24 hrs post-exposure, with subsequent active metabolites likely in circulation after this point. The anti-gametocytocidal activity of PQ is clearly demonstrated prior to this time point, which was the maximal point of transmission-blockade observed with this compound in the direct feeding assay (Fig. 3A & B). Conversely, measurable concentrations of TQ were detected in all samples collected from mice following administration of a single oral dose at 3.41 mg/kg. Concentrations remained relatively constant over the 96-hour sampling period, with a maximum blood-plasma level of 345.9 ng/ml observed at 96 hours. This corresponds with the comparatively increased transmission-blocking efficacy of TQ in the direct feeding assay post 24-hrs (Fig. 3A & B), where TQ exhibits superior potency compared to PQ.

Collectively, these data concisely illustrate the complexities of assessing the activity of dual (asexual and gametocytocidal) compounds on transmission. Firstly, treatment with either PQ and TQ affects asexual and sexual in a similar (and not statistically different) manner, indicating that any differences observed in transmission-blocking efficacy are not solely due to gametocyte clearance. Secondly, the circulating plasma levels of both PQ and TQ demonstrated differing pharmacokinetics, and correlate with reduction in oocyst intensity and prevalence in direct feeding assays (Figs. 3A & B, & S1), highlighting that the time that efficacy is observed (post-treatment) is key to relevant assessment of activity. Thirdly, the (asexual) sub-curative doses of dual activity compounds affect and complicate the downstream analysis of transmission blocking efficacy, by impacting the subsequent development of newly formed mature gametocytes over time. This has obvious implications for the robust and accurate assessment of transmission blocking efficacy.

### Primaquine and Tafenoquine exhibit similar efficacy at clearing asexual and sexual parasitemia when combined with a schizonticide

To reduce the above confounding effects from ongoing asexual replication and better reflect clinical use in which a transmission-blocking drug would likely be co-administered with a curative schizonticide (*44*), DFAs were repeated with the addition of a curative dose (8.4 mg/kg) of sulphadiazine (SD), a compound with no direct gametocytocidal effect (*42*). SD dosage alone was also examined in this manner to determine the baseline reduction in transmission produced by elimination of asexuals and natural turnover of gametocytes. As previously, effect on intensity and prevalence of infection after a direct feed following drug treatment was performed (Fig. 4A & B), and impact on asexual parasitemia and gametocytemia (Fig. 4C & D) was measured. Twenty-four hours post drug-treatment, the addition of either PQ or TQ to SD results in a large reduction in both oocyst intensity and prevalence (76% and 70% respectively), whereas use of SD alone results in no detectable transmission blocking. Forty-eight hours post-treatment, significant reductions in transmission are observed in both PQ+SD and TQ+SD groups, whereas significant reduction is not observed in SD (alone) treated mice. At this time-point, TQ+SD marginally, but significantly, outperforms PQ+SD in terms of reduction in prevalence (Fig. 4B). By 72 hours post-dosage, 100% transmission-blocking efficacy is observed for all groups, due to the depletion of the viable circulating gametocyte pool by addition of schizonticide. This is supported by the data in Figs. 4C & D, where we observe a linear and predictable decrease in asexual parasitemia in all treated mice, achieving total clearance of parasites detectable by microscopy at 72 hours post-treatment (Fig 4C). Whilst at lower levels, reductions in gametocytemia follow a similar trend (Fig, 4D). No significant differences in asexual parasitemia or gametocytemia were observed between any drug regime.

**Figure 4:**
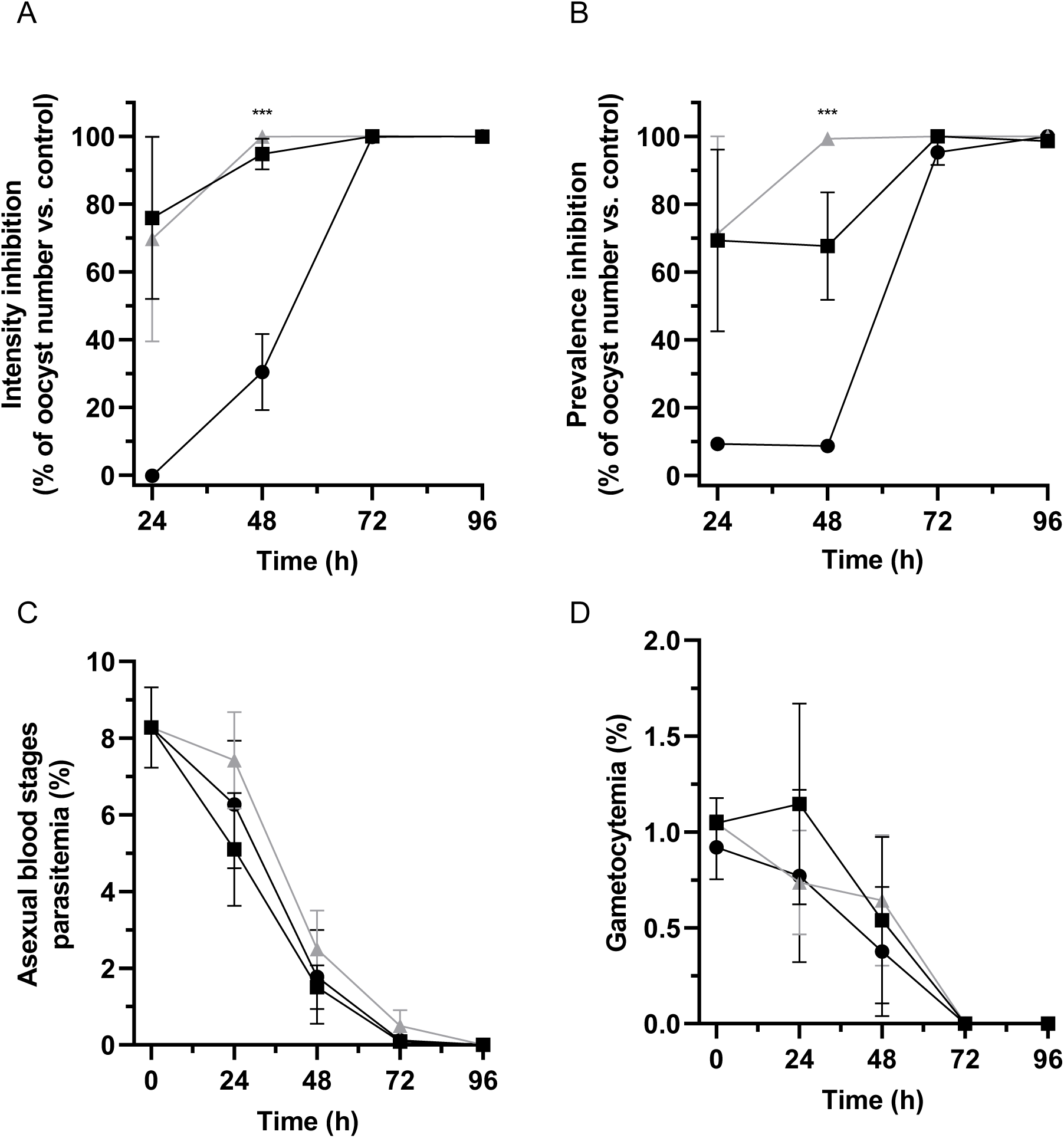
Primaquine and tafenoquine efficacies in the *P. berghei* mouse-to-mouse transmission model at equal molar concentrations, in combination with the schizonticide Sulphadiazine. (A and B) Transmission blocking efficacy in a Direct Feeding Assay (DFA) on *P. berghei*-infected mice orally treated with single doses of PQ (dosed at 3.35 mg.kg^-1^) and TQ (dosed at 3.41 mg.kg^-1^) plus 8.4 mg.kg^-1^ of sulphadiazine, or sulphadiazine alone; and fed to mosquitoes at 24-, 48-, 72- and 96-hours post drug administration. Oocyst intensity (A) and prevalence (B) were then determined 10 days after feeding (day 19 post-infection) and compared to carrier control-treated mice. then determined 10 days after feeding (day 19 post-infection) and compared to carrier control-treated mice. The difference in intensity (Mann-Whitney test) and in prevalence (Fisher’s exact test) between PQ and TQ treatment was significant only at 72 hours post drug administration (p < 0.0001, ***). (C and D) Efficacy of PQ and TQ in combination with sulphadiazine, or sulphadiazine alone at reducing asexual (C) and sexual (D) blood stages. On day 30 post-infection, mice were expose to potentially infectious bites and blood smears were taken for up to 10 days post-bite (day 40 post-infection) to quantify blood stages and gametocytes. No significant differences were observed between PQ+sulphadiazine, TQ+sulphadiazine and sulphadiazine regarding asexual and gametocyte parasitemias (time-points t-test comparisons). Mean +/− SEM of 3 independent experiments.

### Gametocytocidal efficacy of primaquine and tafenoquine in the *P. falciparum* humanized NSG mouse model

Given the potential effects of reduction in asexual parasitemia on transmission blocking efficacy, and the differences in parasite biology between murine and human malaria parasites, the gametocytocidal effects of PQ and TQ were further assessed in a *P. falciparum in vivo* transmission model (Fig. 1B). NGS mice were engrafted with human erythrocytes, as previously described (*7*) and subsequently infected with *P. falciparum* stage V gametocytes expressing a RE9H luciferase under control of the stage V specific ULG8 promotor. Twenty-four hours post infection, mice were treated with a single oral dose of PQ or TQ at a range of concentrations. Circulation and clearance of gametocytes from the blood stream was then continually monitored in mice by bioluminescence analysis and by light microscopy observation of peripheral blood smears, up to 16 days post-infection. Both PQ and TQ induced clearance of gametocytes in a dose-response manner (Fig. 5, Fig. S2). High doses of TQ (100 and 70 mg/kg) and PQ (50 mg/kg) showed comparable effects, with gametocytemia decreasing below the detection limit of both read-outs on day 4 post-infection. Comparatively, lower doses, respectively 20, 10 and 5 mg/kg of PQ and 20 and 5 mg/kg of TQ, showed a delayed clearance of gametocytes (Fig. S3, Table S7 and S8).

**Figure 5:**
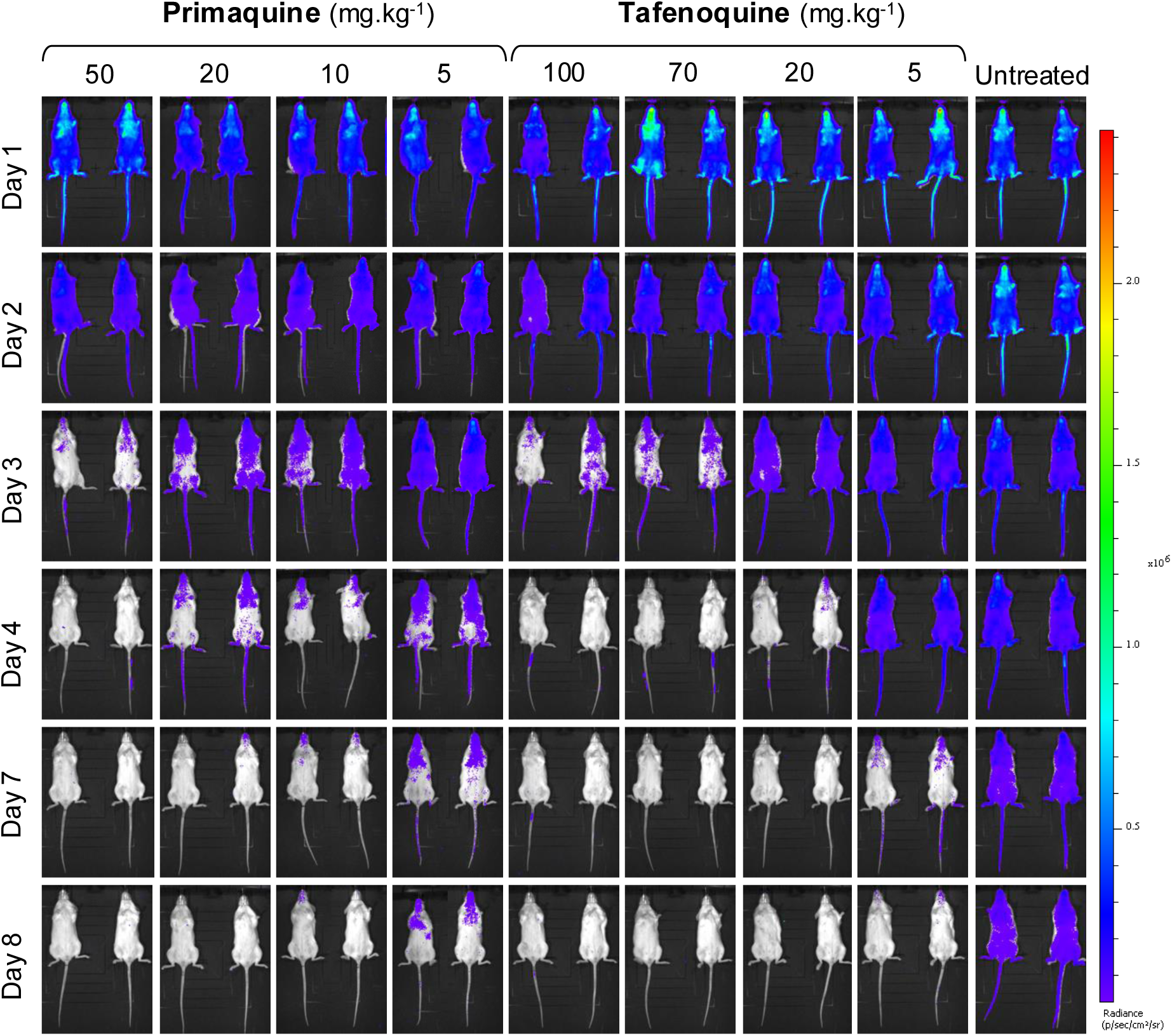
Qualitative therapeutic efficacy against *P. falciparum* NF54/iGP_RE9H gametocytes determined by bioluminescence in NSG humanized mice. Representative ventral images of mice infected with 2×10^8^ stages IV/V gametocytes. Mice were infected on day 0, treated on day 1 with various doses of PQ (left) or TQ (right) and compared to untreated control mice. Pseudo color heat-maps indicate intensity of detected bioluminescence from low (blue) to high (red). Presentative images for all the doses used in this study are displayed in Figure S2.

The calculated ED_50_ (50% reduction in gametocytemia) values using the bioluminescence read-out at day 4 post-infection did not significantly differ between PQ and TQ (respectively 7.5 and 10 mg/kg, extra sum-of-squares F-test, p-value = 0.32), but showed a significant difference at day 7 post-infection (respectively 6.2 and 1 mg/kg, extra sum-of-squares F-test, p-value = 0.039) (Fig. 6). Similar results were obtained using the microscopy read-out at day 4 post-infection, although no significant difference was observed between the two antimalarials at day 7 post-infection (Fig. S5). In contrast to PQ, which showed a rapid decay in exposure, TQ displayed a sustained exposure up to 5 days post-treatment (Fig. S4, Table S9).

**Figure 6:**
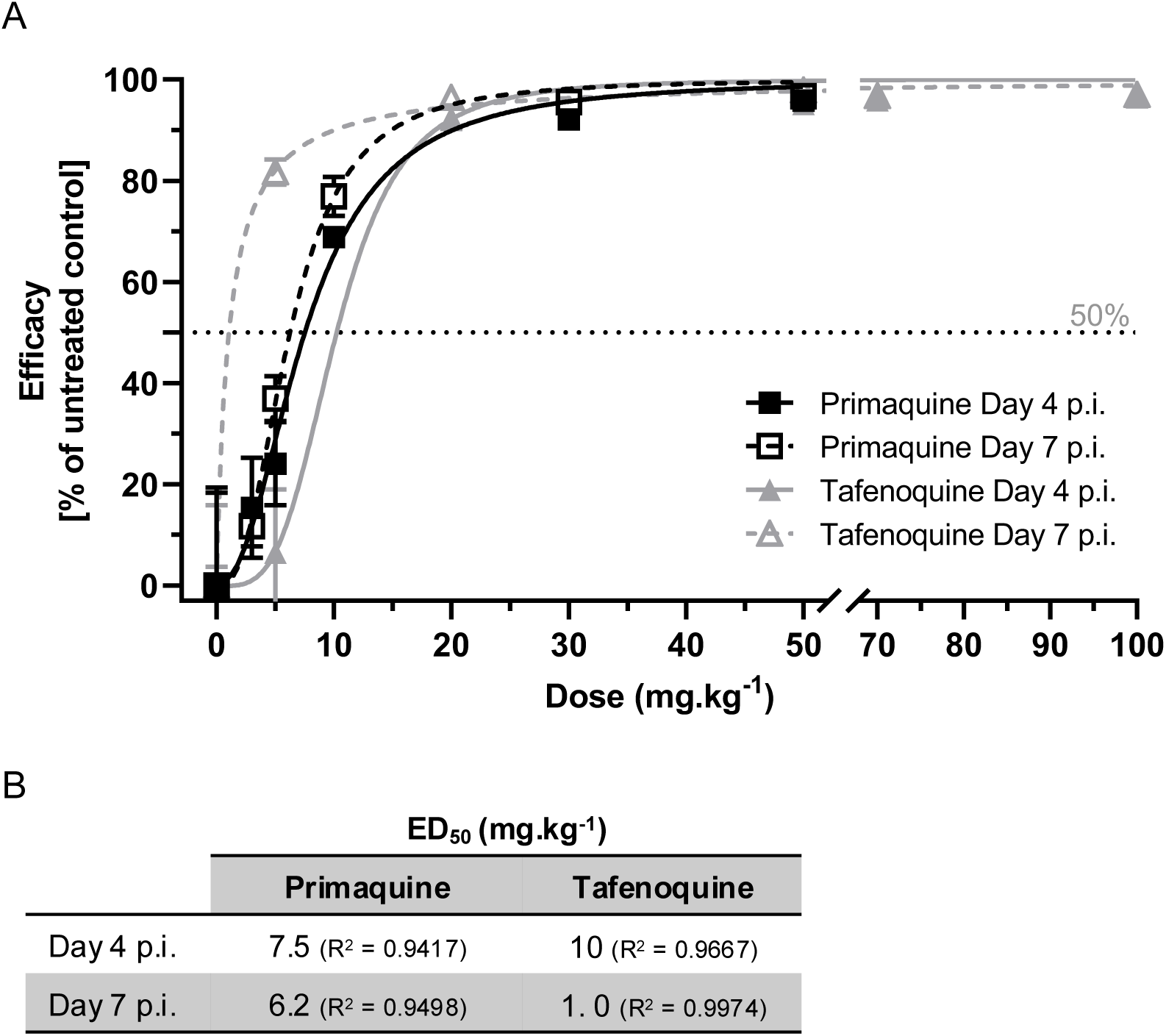
Efficacy of primaquine and tafenoquine compared to untreated control against *P. falciparum* NF54/iGP_RE9H stages IV/V gametocytes in NSG humanized mice. (A) Efficacy was expressed based on the measured luciferase bioluminescence compared to the untreated control mice (Efficacy=100-(RLU treated mice *100/ RLU untreated controls), on Day 4 (full line) and Day 7 (dashed line) post-infection (p.i.) Non-linear regressions were performed in GraphPad Prism (variable slope, four parameters). Mean +/− SD of at least 2 mice. The dotted line represents 50% of efficacy. (B) Summary of the ED_50_ values obtain at each timepoint.

Although both methods of observation led to similar results, the bioluminescence read-out indicated a faster onset of clearance compared to observation of blood smears. Within 24h post treatment a reduction of gametocytemia was measurable for high treatment doses (Fig. S3) with the bioluminescent readout, whereas the conventional readout via blood smears indicated a lag phase of >24h for PQ and TQ.

## Discussion

In this study, we used complementary preclinical transmission models to directly compare the transmission-blocking efficacy of the 8-aminoquinolines primaquine (PQ) and tafenoquine (TQ), administered as single-dose agents both alone, and in combination with a curative schizonticide. Given the requirement for hepatic metabolism to generate active species for both compounds (18,19), *in vivo* assessment is essential for evaluating their transmission-blocking potential and informing therapeutic usage. By integrating dose titration, temporal assessment of efficacy, pharmacokinetic analysis, and multiple parasite assay systems, our data provide a comparative framework for informed assessment of understanding the strengths and limitations of PQ and TQ as transmission-blocking compounds in the clinic.

We initially describe the first (to our knowledge) titration of the transmission blocking efficacy of single doses of PQ (from 0.1 mg/kg to 12 mg/kg) to reduce transmission to mosquitoes using a range of *P. berghei* DFA experiments (Fig. 2). Titration of efficacy was significant, predictable and robust, in terms of reduction in oocyst intensity and infection prevalence, with (maximal) 100% inhibition in both metrics observed by 6mg/kg. ED_50_’s of 3.35 mg/kg and 1.26 mg/kg were calculated for reduction in intensity and prevalence, respectively. While higher PQ doses predictably increased transmission-blocking efficacy, their clinical relevance is constrained by safety concerns in G6PD-deficient individuals, reinforcing the importance of optimizing efficacy at low or single-dose regimens.

Following titration, both PQ and TQ were used at ED_50_ in a transmission-based M2M2M model (*5,6,42,43*) that encompassed the full plasmodial lifecycle. The results observed under the tested conditions (mouse-to-mosquito transmission 24 hrs post drug-treatment) indicated that PQ demonstrated modestly superior efficacy to TQ in terms of reducing transmission to the mosquito, and to subsequent naïve mouse populations, after mosquito derived transmission (Table 1). The effect size of 27.1% observed following PQ treatment at 3.35 mg/kg is logical, given the previous effect sizes observed using PQ at lower doses, where an effect size of 8% was observed within this model at a dose of 0.25 mg/kg (*42*). This information aligns with clinical data presented in (*40*), where use of ACT+PQ demonstrated a significantly higher impact on gametocyte infectivity early (2 days) post-dosage when compared to ACT+TQ. The observed marginal superiority of PQ at this time point drove us to directly compare the efficacy of PQ and TQ at ED_50_ doses over a longer time scale (24-, 48-, 72- and 96-hours post-dosage) in direct feeding assays (Figure 3). Consistent with the M2M2M data, PQ significantly outperformed TQ in terms of reduction in transmission to the mosquito (Figs. 3A & B) at 24 hours post exposure, despite comparable circulating gametocyte densities (Fig. 3D). Two possibilities suggest themselves regarding this point; firstly, that the gametocytes observed within only PQ-treated mice 24 hours post-exposure are dead but clearly visible and morphologically unchanged in the blood (*45*); or secondly, that PQ has a biological activity that is acting on a wider range of parasite biology than simple destruction of mature gametocytes. This supports historical data from 12 patients that suggested escalation of PQ efficacy against *P. falciparum* as it passed through mosquito stage development (*46*). Further examination of this may be prudent in the future, especially as the data presented here indicate that simple measurement of reduction of gametocyte number may not always be predictive of reduction of transmission to mosquitoes in feeding assays.

At later time points, this relationship was reversed. TQ consistently outperformed PQ from 48 h onwards, with sustained and pronounced significant reductions in oocyst intensity and prevalence. Pharmacokinetic analysis provided a clear mechanistic explanation: PQ was undetectable in plasma beyond 24 h, whereas TQ exhibited prolonged exposure over the full sampling period (S1, tables S3). The extended half-life and bioavailability of TQ therefore confers a broader time window for transmission blockade, accounting for the superior longer-term efficacy observed here. Observation of longer-term blood-plasma levels of TQ may be advantageous. These data have obvious and significant implications for clinical use of TQ as a single dose transmission blocking drug but also underscore the importance of aligning efficacy assessments with pharmacokinetic profiles and caution against single timepoint comparisons when evaluating transmission-blocking agents.

The addition of a curative schizonticide (SD 8.4 mg/kg) markedly enhanced transmission blockade for both PQ and TQ (Fig. 4), consistent with clinical evidence supporting combination regimens. For example, recent field data demonstrated that a single low dose of PQ at 0.2 mg/kg, when added to ACTs and administered to patients, reduced *P. falciparum* gametocytemia and infectivity to mosquitoes at day 7 and was well tolerated, regardless of the age of the patients and the type of ACT given (*20*). Similar outcomes have been observed following the addition of a single dose of 0.25mg/kg of PQ to artesunate amodiaquine (*47*) and to pyronaridine-artesunate or dihydroartemisinin-piperaquine (*39,48*). Similarly, this also supports findings where supplementation of TQ with dihydroartemisinin-piperaquine (*39*) or sulfadoxine-pyrimethamine-amodiaquine (*47*) resulted in enhanced or full transmission blockade respectively. This data highlights that the combination of transmission-blocking compounds with a schizonticide results in enhanced and longer-lasting transmission blocking efficacy, supporting MMV’s current Target Product Profile 1 (TPP-1), incorporating the use of two or more molecules with asexual activity in combination with a compound that reduces transmission (*49*). This is particularly relevant in *P. falciparum*, where gametocytes take 14 days to mature (*50*). In *P. berghei*, this process is much shorter, with a 24-hour window for gametocytogenesis (*5*). Importantly, our data also illustrate that depletion of asexual parasitemia over time can substantially influence observed transmission-blocking efficacy, a critical consideration when evaluating dual-activity compounds in transmission blocking assays.

To assess gametocytocidal activity independently of asexual-stage effects and in a human parasite context, we employed a humanized NSG mouse model infected with *P. falciparum* stage V gametocytes. Again, treatment with both compounds resulted in dose-dependent clearance of gametocytes. At early timepoints post infection and treatment, ED_50_s (as determined by reduction in bioluminescence) did not significantly vary between PQ and TQ at day 4, but showed significant difference at day 7 post-infection, where TQ exhibited a significantly more potent ED_50_ than PQ (1 mg/kg compared to 6.2 mg/kg). As in the previous *P. berghei* mosquito-feeding experiments, this data further stresses that when expressing efficacy *in vivo* the timing of both the intervention and the efficacy measure need to be carefully considered. Within the NGS humanized mice model, there are two distinct outputs, bioluminescence of NF54/iGP_RE9H stage IV/V gametocytes, or detection of gametocytes by light microscopy. Both outputs generate similar results and trends, with the bioluminescence read-out giving a faster onset of clearance compared to the blood smears. This is likely due to biological differences between these measurables, with dead gametocytes still potentially indistinguishable from live ones by Giemsa stain. Conversely, active and detectable bioluminescence requires the ATP present in live parasites, allowing for a more dynamic identification of live parasites (*7*).

The experiments described here encompass a range of pre-clinical models to assess the transmission-blocking efficacy of both PQ and TQ. Each assay has strengths and weaknesses that must be considered in terms of translational predictivity. The use of *P. falciparum* or *P. berghei* Standard Membrane Feeding Assays (SMFAs) is inappropriate for the assessment of these compounds, due to their requirement for liver metabolism. The use of rodent malaria (i.e., *P. berghei*) direct feeding assays allow for the oral dosing of infected mice, which results in liver metabolism and delivery of the active species, followed by readouts of asexual parasitemia, gametocytemia, and crucially, reduction in transmission to mosquitoes. This is potentially key if the compound has a sporontocidal mode of action independent of, or in addition to, reduction in gametocytemia (*46*), and can be performed at multiple time points post drug-treatment. The negatives of this approach are that rodent malaria parasites exhibit differences in biology when compared to *P. falciparum*, assessing PK/PD is difficult if activity is conferred through transient metabolites and thus translational predictivity is uncertain, and that the assay window only covers a relatively small part of the parasite lifecycle (mature gametocyte to oocyst). To aid the latter, a M2M2M model can be used to examine the impact of treatment on transmission from vertebrate-to-mosquito-to-vertebrate across the entire parasite lifecycle, across a range of transmission settings/biting rates. Although high biological content is achieved, the assay is time consuming (with an approximately 41-day length), and again, uses rodent malaria parasites. The use of a humanized NSG mouse model, infected with *P. falciparum* mitigates this, with use of mature, stage V NF54/iGP_RE9H gametocytes giving evidence of direct gametocytocidal action using human malaria parasites *in vivo*. The use of two distinct outputs (blood smears and bioluminescence) gives increased biological content and sensitivity (*7*). Conversely, the system only allows for detection of impact on a relatively narrow window, and measurement of efficacy on immature to stage V gametocytes. As yet, mosquito feeding assays have not been accomplished using this model but are in development to determine minimum gametocyte sterilising concentrations *in vivo*.

The application of these models for the comparative assessment of PQ and TQ clearly demonstrates that a multi-faceted and diverse approach is needed to fully assess the efficacy of transmission blocking compounds, with consideration needed in terms of durability, time post-treatment and dose. Nevertheless, use of this collection of assays within this context has led to the first *in vivo* titration of the transmission blocking efficacy of both PQ and TQ, demonstrating potent activity with both compounds. While PQ provides stronger early effects, TQ confers superior and more durable transmission blockade beyond 24 h due to its prolonged exposure. These findings, observed across multiple models and in the presence or absence of a schizonticide, suggest that single-dose TQ may have potential as a transmission-blocking intervention in humans.

## Material and Methods

### *P. berghei* mouse-to-mouse transmission model

#### Drug treatments

Primaquine bisphosphate was obtained from Sigma-Aldrich (160393), tafenoquine succinate from Sigma-Aldrich (SML0396), and sulphadiazine from Sigma-Aldrich (S6387). All compounds were diluted in 1% Methyl Cellulose (Sigma-Aldrich no. A7968) which was used as a negative, no drug control for treatment.

#### Parasite maintenance

General parasite maintenance was carried out as described in (*43*). Briefly, *P. berghei* ANKA 2.34 parasites were maintained in 6–8-week-old female Tuck Ordinary (TO) mice (Harlan) by serial mechanical passage (up to a maximum of eight passages). If required, hyper-reticulosis was induced three days before infection by treating mice intraperitoneally (*i.p*) with 200 μl phenylhydrazinium chloride (PH; 6 mg/ml in PBS; ProLabo UK). Mice were infected *i.p*. and infections were monitored using Giemsa-stained tail blood smears as described previously (*5*).

#### Direct feeding assay (DFA)

To identify the primaquine concentration that would result in a 50% reduction in infection prevalence, eighteen individual direct feeding assays (DFAs) were performed at a selection of primaquine concentrations, ranging from 0.1 mg/kg to 12 mg/kg. Briefly, prior to challenge, mice were PH treated, and 3 days later infected intraperitoneally (*i.p*). with 10^6^ *P. berghei* ANKA 2.34 asexual parasites. Two days post-infection, animals were orally dosed with primaquine in 1% methyl cellulose (MeC) at the required dose, or with 1% MeC alone (untreated group). Twenty-four hours post-treatment, individual animals were anesthetized, and >50 female *Anopheles stephensi* mosquitoes allowed to blood feed on each mouse. Unfed mosquitoes were removed, and remaining mosquitoes maintained on 8% (w/v) fructose, 0.05% (w/v) p-aminobenzoic acid at 19-22 ͦC and 50-80% relative humidity. Day 14 post-feeding, mosquito midguts were dissected, and oocyst intensity and prevalence observed by standard phase microscopy and recorded. The required primaquine concentrations were estimated using the fitted values of the most appropriate model, and 95% confidence intervals were obtained from the profile likelihood. The primaquine concentrations required to achieve a 50% reduction in prevalence and intensity were estimated from the best-fit logistic curves, and 95% confidence intervals were estimated from the profile likelihood.

#### In vivo mouse-to-mouse transmission blocking assay

The mouse-to-mouse transmission assay was carried out as previously described (*6,42,43*) (Fig. 1A). Briefly, groups of 5 female TO mice (6- to 8-weeks old) *i.p*. infected with *P. berghei* ANKA 2.34 blood stage parasites and examined for male gametocyte exflagellation in tail blood smears 8 days post-infection. The mice were subsequently treated with a single oral dose of PQ (3.41 mg/kg) or TQ (3.35 mg/kg). After 24h of treatment (day 9 post-infection), 500 starved female *A. stephensi* (line SD 500) were allowed to feed on these anesthetized mice. Mosquitoes that did not take a blood meal were discarded, and the remaining ones were maintained on 8% (w/v) fructose and 0.05% (w/v) p-aminobenzoic acid at 19°C and 80% relative humidity. Ten days later (day 19 post-infection), the midguts of random 50 fed-mosquitoes per cage were dissected and oocyst intensity (mean number of parasites per gut) and prevalence (percentage of infected mosquitoes) were determined and compared to the untreated group to calculate the “classical” inhibition of transmission. All mosquito dissections to assess oocyst intensity and prevalence were performed under randomized and double-blind conditions. Twenty-one days after the feeding (day 30 post-infection), when salivary glands sporozoites are estimated to be most infectious (*5*), for each compound tested, naïve anesthetized mice (3 groups of n=5) were exposed to 2, 5 or 10 potentially infectious mosquitoes for 20 minutes. Successful feeding was confirmed by the presence of blood in the mosquito abdomen. Where necessary, additional mosquitoes were given the opportunity to feed until the required number of successful bites was achieved per mouse. The bitten mice were monitored for up to 10 days post-bite (up to day 40 post-infection) with daily tail blood smears to measure parasitemia (asexual stages and gametocytes) and time to patency.

#### Blood sample analysis for primaquine and tafenoquine

Blood samples were collected during the study (24, 48, 72 and 96h post treatment) and immediately frozen at –80°C. Duplicate samples (50 μl) from each time point were shipped on dry ice to the Centre for Drug Candidate Optimisation, Monash University, Australia for analysis of primaquine and tafenoquine concentrations. Samples were prepared by precipitating the proteins with addition of 2 volumes of acetonitrile containing the internal standard diazepam. followed by centrifugation and separation of the supernatant. Calibration standards were prepared by spiking blank mouse blood and treating in the same manner. Samples and standards were assayed for primaquine or tafenoquine were quantified by LC/MS/MS using a Waters Micromass Quattro Premier mass spectrometer coupled to a Waters Acquity UPLC. The column was a Supelco Ascentis Express RP Amide column (50×2.1 mm, 2.7 μm) and the mobile phase consisted of a water and acetonitrile, each containing 0.05% formic acid and mixed using a 4 min gradient. The injection volume was 3 µL and the flow rate was 0.4 mL/min. Detection was conducted using positive electrospray ionisation multiple-reaction monitoring mode with the transition (m/z), cone voltage (V) and CID (V) being 260.75 > 86.24, 25 and 15, respectively for primaquine and 464.23 > 85.97, 35 and 20, respectively for tafenoquine. The assay calibration ranges were 5-5000 ng/mL (primaquine) and 1-10,000 ng/mL (tafenoquine) and accuracy and precision were within ± 10% and <10%, respectively, for both analytes.

#### Statistical analysis

For the titration of PQ efficacy, reductions in intensity and prevalence in primaquine-treated mice were calculated with respect to control, non-drug treated mice within individual experiments. Efficacies were estimated as functions of primaquine concentrations using a generalized linear model framework (*51*) in which each experimental replicate was treated as a random effect. Two curvilinear functions – logistic and Gompertz (*52*) – were fitted to the data using maximum likelihood, and their fits compared. The relationships between prevalence efficacy and primaquine concentrations were best described by a logistic function, which fit the data significantly better than a Gompertz function in both cases (likelihood ratio tests, p > 0.5).

For the mouse-to-mouse assay, generalized linear mixed models were used to estimate the overall effectiveness of the different interventions combining data from all repeat replicates (*6,42,43*). For the oocyst data we assumed a binomial error structure while for parasitemia data, a zero-inflated negative binomial distribution was used. Ninety-five percent confidence intervals were estimated by bootstrapping, and the model was selected using a likelihood ratio test. Effect size was calculated by use of a chain binomial model, with models fitted to the data using maximum likelihood methods and the 95% confidence interval estimates obtained from the likelihood profile.

For DFAs, analysis was performed with Graphpad Prism Software (Graphpad Software Inc.). Significant differences in oocyst intensity were assessed using the Mann–Whitney-U test (P < 0.05), whereas Fisher’s exact probability test was used to assess differences in infection prevalence (P < 0.05). Differences in parasitemia and gametocytemia were assessed by standard t-test performed at least in biological triplicate (P < 0.05).

### *P. falciparum* humanized NSG mice gametocytocidal model

#### Drug treatments

Primaquine (MMV000023-02, Primaquine di-phosphate) was tested at seven concentrations (50, 30, 10, 5, 3, 1, 0.3 mg/kg body weight) and Tafenoquine (MMV000043-08, Tafenoquine succinate) at five concentrations (100, 70, 50, 20, 5 mg/kg body weight) on at least 2 mice.

#### In vitro production of enriched P. falciparum stages IV/V gametocytes

Highly synchronized *P. falciparum* stages IV/V gametocytes were obtained as described in (*7*). Briefly, synchronous ring-stage parasites of the transgenic NF54/iGP1_RE9H parasite line (4% parasitemia, 5% hematocrit, 10% human serum) were cultured in absence of glucosamine (GlcNAc) and in presence of 0.625 nM Shield-1 under static conditions for 24h, followed by 24h under shaking, to allow high sexual commitment of the parasites (up to 80%, (*7*)). Parasites were subsequently cultured for 10 days in presence of GlcNAc to block further sexual commitment and keep the gametocytes synchronized. This process resulted in highly enriched (up to 3%) stages IV/V gametocytes (*7*).

#### In vivo gametocytocidal efficacy study

NSG mice were engrafted intravenously with human erythrocytes 11 days prior the beginning of the experiment, and at regular interval until the end of the study (*53*). These “humanized” mice were intravenously infected with highly enriched *in vitro*-induced *P. falciparum* NF54/iGP_RE9H stages IV/V gametocytes in human erythrocytes as described above and treated with a single oral dose of primaquine or tafenoquine in 70% Tween-80 and 30% ethanol diluted 1/10 in water (vehicle) or the vehicle alone (untreated group), on the next day (day 1 post-infection). The gametocytemia were monitored by live imaging (bioluminescence, enabled by the red-shifted luciferase RE9H from the gametocyte-specific ulg8 locus (PF3D7_1234700)) and microscopy (gametocyte count from Giemsa-stained tail blood smears) up to day 16 post-infection ((*7*), Fig. 1B).

#### Pharmacokinetic analysis

Blood samples were collected during the study (2, 4 and 24h post-treatment for Primaquine; 2, 4, 24, 48, 72 and 120h for Tafenoquine). Samples were diluted at a 1:1 ratio in water (v/v) and kept at –80°C until shipment in dry ice to Swiss BioQuant for analysis. After precipitation in 3 volumes of acetonitrile containing the internal standard Reserpine, the supernatants were transferred into autosampler vials. Both primaquine and tafenoquine were quantified by LC-MS/MS using HESI ionization in positive mode. A noncompartmental analysis was performed for the determination of the pharmacokinetic parameters, using the Phoenix 64 WinNonlin program (version 8.0).

#### Statistical analysis

Primaquine and tafenoquine gametocytocidal efficacies were expressed as the percent of the untreated control mice group of both read-outs at days 4 and 7 post-infection. Non-linear regressions (variable slope, four parameters) were performed using GraphPad Prism 9.

## Ethical statement

All procedures were performed in accordance with the UK Animals (Scientific Procedures) Act (PP8697814) and approved by the University of Cambridge AWERB. The Office of Laboratory Animal Welfare Assurance for the University of Cambridge covers all Public Health Service supported activities involving live vertebrates in the US (no. A5634-01). This study was carried out in compliance with the ARRIVE guidelines (https://arriveguidelines.org/). *In vivo* studies conducted at the Swiss TPH, Basel, were approved by the veterinary authorities of the Canton Basel-Stadt (permit no. 2992) based on Swiss cantonal (Verordnung Veterinäramt Basel-Stadt) and national regulations (permit no. N-30762, the Swiss animal protection law, Tierschutzgesetz).

## Acknowledgements

We thank M.A. Westwood (Swiss BioQuant) for the pharmacokinetic analysis of the humanized mice samples. The Centre for Drug Candidate Optimisation (CDCO), Monash University, Australia is acknowledged for analyzing the blood samples in the *P. berghei* experiments.

## Funding

Funding for this work was provided by MMV Medicines for Malaria Venture (MMV). A.M.B thanks MMV for funding, and is supported by the MRC [MR/N00227X/1 and MR/W025701/1], Sir Isaac Newton Trust, Alborada Fund, Wellcome Trust ISSF and University of Cambridge JRG Scheme, GHIT, Rosetrees Trust (G109130) and the Royal Society (RGS/R1/201293) (IEC/R3/19302). The CDCO partially supported by Monash University and Therapeutic Innovation Australia (TIA) through the Australian Government National Collaborative Research Infrastructure Strategy (NCRIS) program.

## Competing interests

M.D. was and D.L. is employee of MMV.

## References

1). World Health Organization, World Malaria Report 2025, WHO Press (2025).

2). Woldegerima, WA, Ouifki R, and Banasiak J. Mathematical analysis of the impact of transmission-blocking drugs on the population dynamics of malaria. Applied Mathematics and Computation 400: 126005. (2021).

3). A. B. Tiono, A. Ouédraogo, B. Ogutu, A. Diarra, S. Coulibaly, A. Gansané, S. B. Sirima, G. O’Neil, A. Mukhopadhyay, K. Hamed, A controlled, parallel, cluster-randomized trial of community-wide screening and treatment of asymptomatic carriers of *Plasmodium falciparum* in Burkina Faso., Malaria Journal 12 (2013), doi:10.1186/1475-2875-12-79.

4). M. van der Kolk, S. J. de Vlas, A. Saul, M. van de Vegte-Bolmer, W. M. Eling, R. W. Sauerwein, W. Sauerwein, Evaluation of the standard membrane feeding assay (SMFA) for the determination of malaria transmission-reducing activity using empirical data., Parasitology 130, 13–22 (2005).

5). A. M. Blagborough, T. S. Churcher, L. M. Upton, A. C. Ghani, P. W. Gething, R. E. Sinden, Transmission-blocking interventions eliminate malaria from laboratory populations, Nature Communications 4 (2013), doi:10.1038/ncomms2840.

6). T. Paquet, C. le Manach, D. G. Cabrera, Y. Younis, P. P. Henrich, T. S. Abraham, M. C. S. Lee, R. Basak, S. Ghidelli-Disse, M. J. Lafuente-Monasterio, M. Bantscheff, A. Ruecker, A. M. Blagborough, S. E. Zakutansky, A. M. Zeeman, K. L. White, D. M. Shackleford, J. Mannila, J. Morizzi, C. Scheurer, I. Angulo-Barturen, M. Santosmartínez, S. Ferrer, L. M. Sanz, F. J. Gamo, J. Reader, M. Botha, K. J. Dechering, R. W. Sauerwein, A. Tungtaeng, P. Vanachayangkul, C. S. Lim, J. Burrows, M. J. Witty, K. C. Marsh, C. Bodenreider, R. Rochford, S. M. Solapure, M. B. Jiménez-Díaz, S. Wittlin, S. A. Charman, C. Donini, B. Campo, L. M. Birkholtz, K. Khanson, G. Drewes, C. M. Kocken, M. J. Delves, D. Leroy, D. A. Fidock, D. Waterson, L. J. Street, K. Chibale, Antimalarial efficacy of MMV390048, an inhibitor of *Plasmodium* phosphatidylinositol 4-kinase, Science Translational Medicine 9 (2017), doi:10.1126/scitranslmed.aad9735.

7). Brancucci NMB, Gumpp C, van Gemert GJ, Yu X, Passecker A, Nardella F, Thommen BT, Chambon M, Turcatti G, Halby L, Blasco B, Duffey M, Arimondo PB, Bousema T, Scherf A, Leroy D, Kooij TWA, Rottmann M, Voss TS. An all-in-one pipeline for the *in vitro* discovery and *in vivo* testing of *Plasmodium falciparum* malaria transmission blocking drugs. Nat Commun. 2025 16(1):6884 (2025).

8). J. C. van Pelt-Koops, H. E. Pett, W. Graumans, M. van der Vegte-Bolmer, G. J. van Gemert, M. Rottmann, B. K. S. Yeung, T. T. Diagana, R. W. Sauerwein, The spiroindolone drug candidate NITD609 potently inhibits gametocytogenesis and blocks *Plasmodium falciparum* transmission to Anopheles mosquito vector, Antimicrobial Agents and Chemotherapy 56, 3544–3548 (2012).

9). K. L. Kuhen, A. K. Chatterjee, M. Rottmann, K. Gagaring, R. Borboa, J. Buenviaje, Z. Chen, C. Francek, T. Wu, A. Nagle, S. W. Barnes, D. Plouffe, M. C. S. Lee, D. A. Fidock, W. Graumans, M. van de Vegte-Bolmer, G. J. van Gemert, G. Wirjanata, B. Sebayang, J. Marfurt, B. Russell, R. Suwanarusk, R. N. Price, F. Nosten, A. Tungtaeng, M. Gettayacamin, J. Sattabongkot, J. Taylor, J. R. Walker, D. Tully, K. P. Patra, E. L. Flannery, J. M. Vinetz, L. Renia, R. W. Sauerwein, E. A. Winzeler, R. J. Glynne, T. T. Diagana, KAF156 is an antimalarial clinical candidate with potential for use in prophylaxis, treatment, and prevention of disease transmission, Antimicrobial Agents and Chemotherapy 58, 5060–5067 (2014).

10). T. Bousema, L. Okell, S. Shekalaghe, J. T. Griffin, S. Omar, P. Sawa, C. Sutherland, R. Sauerwein, A. C. Ghani, C. Drakeley, Revisiting the circulation time of *Plasmodium falciparum* gametocytes: molecular detection methods to estimate the duration of gametocyte carriage and the effect of gametocytocidal drugs., Malaria Journal 9, 136 (2010).

11). N. J. White, Primaquine to prevent transmission of *falciparum* malaria. The Lancet Infectious Diseases 13, 175 (2013).

12). D. J. Hayes, C. G. Banda, A. Chipasula-Teleka, D. J. Terlouw, Modelling the therapeutic dose range of single low dose primaquine to reduce malaria transmission through age-based dosing, BMC Infectious Diseases 17 (2017), doi:10.1186/s12879-017-2378-9.

13). A. Dicko, M. E. Roh, H. Diawara, A. Mahamar, H. M. Soumare, K. Lanke, J. Bradley, K. Sanogo, D. T. Kone, K. Diarra, S. Keita, D. Issiaka, S. F. Traore, C. McCulloch, W. J. R. Stone, J. Hwang, O. Müller, J. M. Brown, V. Srinivasan, C. Drakeley, R. Gosling, I. Chen, T. Bousema, Efficacy and safety of primaquine and methylene blue for prevention of *Plasmodium falciparum* transmission in Mali: a phase 2, single-blind, randomised controlled trial, The Lancet Infectious Diseases 18, 627–639 (2018).

14). A. W. Sweeney, Wartime research on malaria chemotherapy, Parassitologia 42, 33–45 (2000).

15). D. Greenwood, Conflicts of interest: the genesis of synthetic antimalarial agents in peace and war, The Journal of antimicrobial chemotherapy 36, 857–872 (1995).

16.). P. E. Carson, in Antimalarial Drug II, (1984), pp. 83–121.

17). J. K. Baird, K. H. Rieckmann, Can primaquine therapy for vivax malaria be improved? Trends in Parasitology 19, 115–120 (2003).

18). Camarda G, Jirawatcharadech P, Priestley RS, Saif A, March S, Wong MHL, Leung S, Miller AB, Baker DA, Alano P, Paine MJI, Bhatia SN, O’Neill PM, Ward SA, Biagini GA. Antimalarial activity of primaquine operates via a two-step biochemical relay. Nat. Commun. 10, 3226. (2019).

19). Marcsisin, S. R., Reichard G., Pybus B. S., Primaquine pharmacology in the context of CYP 2D6 pharmacogenomics: Current state of the art, Pharmacology & Therapeutics, 161, Pages 1–10, ISSN 0163-7258 (2016).

20). Yilma D, Stepniewska K, Bousema T, Drakeley C, Eachempati P, Guerin PJ, Mårtensson A, Mwaiswelo R, Taylor WR, Barnes KI; WWARN Paediatric Primaquine for *P falciparum* Transmission Blocking Study Group. Safety and efficacy of single-dose primaquine to interrupt *Plasmodium falciparum* malaria transmission in children compared with adults: a systematic review and individual patient data meta-analysis. Lancet Infect Dis. 25(9):965–976. (2025).

21). Taylor WR, Olupot-Olupot P, Onyamboko MA, Peerawaranun P, Weere W, Namayanja C, Onyas P, Titin H, Baseke J, Muhindo R, Kayembe DK, Ndjowo PO, Basara BB, Bongo GS, Okalebo CB, Abongo G, Uyoga S, Williams TN, Taya C, Dhorda M, Tarning J, Dondorp AM, Waithira N, Fanello C, Maitland K, Mukaka M, Day NJP. Safety of age-dosed, single low-dose primaquine in children with glucose-6-phosphate dehydrogenase deficiency who are infected with *Plasmodium falciparum* in Uganda and the Democratic Republic of the Congo: a randomised, double-blind, placebo-controlled, non-inferiority trial. Lancet Infect Dis. 23(4):471–483.

22). World Health Organization, WHO Guidelines for malaria, WHO Press (2021).

23). B. P. Gonçalves, A. B. Tiono, A. Ouédraogo, W. M. Guelbéogo, J. Bradley, I. Nebie, D. Siaka, K. Lanke, A. C. Eziefula, A. Diarra, H. Pett, E. C. Bougouma, S. B. Sirima, C. Drakeley, T. Bousema, Single low dose primaquine to reduce gametocyte carriage and *Plasmodium falciparum* transmission after artemetherlumefantrine in children with asymptomatic infection: A randomised, double-blind, placebo-controlled trial, BMC Medicine 14 (2016), doi:10.1186/s12916-016-0581-y.

24). G. N. L. Galappaththy, P. Tharyan, R. Kirubakaran, Primaquine for preventing relapse in people with *Plasmodium vivax* malaria treated with chloroquine, Cochrane Database of Systematic Reviews 2013 (2013), doi:10.1002/14651858.CD004389.pub3.

25). Y. R. Kim, H. J. Kuh, M. Y. Kim, Y. S. Kim, W. C. Chung, S. I. Kim, M. W. Kang, Pharmacokinetics of primaquine and carboxyprimaquine in Korean patients with vivax malaria, Archives of Pharmacal Research 27, 576–580 (2004).

26). J. Raman, E. Allen, L. Workman, A. Mabuza, H. Swanepoel, G. Malatje, J. Frean, L. Wiesner, K. I. Barnes, Safety and tolerability of single low-dose primaquine in a low-intensity transmission area in South Africa: An open-label, randomized controlled trial, Malaria Journal 18 (2019), doi:10.1186/s12936-019-2841-8.

27). B. Balikagala, N. Fukuda, M. Ikeda, O. T. Katuro, S.-I. Tachibana, M. Yamauchi, W. Opio, S. Emoto, D. A. Anywar, E. Kimura, N. M. Q. Palacpac, E. I. Odongo-Aginya, M. Ogwang, T. Horii, T. Mita, Evidence of Artemisinin-Resistant Malaria in Africa, New England Journal of Medicine 385, 1163–1171 (2021).

28). Mayence, A.; Vanden Eynde, J.J. Tafenoquine: A 2018 Novel FDA-Approved Prodrug for the Radical Cure of Plasmodium vivax Malaria and Prophylaxis of Malaria. Pharmaceuticals, 12, 115. (2009).

29). J. E. Frampton, Tafenoquine: First Global Approval, Drugs 78, 1517–1523 (2018).

30). Chu CS, Phyo AP, Turner C, Win HH, Poe NP, Yotyingaphiram W, Thinraow S, Wilairisak P, Raksapraidee R, Carrara VI, Paw MK, Wiladphaingern J, Proux S, Bancone G, Sriprawat K, Lee SJ, Jeeyapant A, Watson J, Tarning J, Imwong M, Nosten F, White NJ. Chloroquine Versus Dihydroartemisinin-Piperaquine with Standard High-dose Primaquine Given Either for 7 Days or 14 Days in *Plasmodium vivax* Malaria. Clin Infect Dis. 8;68(8):1311–1319. (2019)

31). N. J. Elmes, P. E. Nasveld, S. J. Kitchener, D. A. Kocisko, M. D. Edstein, The efficacy and tolerability of three different regimens of tafenoquine versus primaquine for post-exposure prophylaxis of *Plasmodium vivax* malaria in the Southwest Pacific, Transactions of the Royal Society of Tropical Medicine and Hygiene 102, 1095–1101 (2008).

32). B. R. Hale, S. Owusu-Agyei, D. J. Fryauff, K. A. Koram, M. Adjuik, A. R. Oduro, W. R. Prescott, J. K. Baird, F. Nkrumah, T. L. Ritchie, E. D. Franke, F. N. Binka, J. Horton, S. L. Hoffman, A randomized, double-blind, placebo-controlled, dose-ranging trial of tafenoquine for weekly prophylaxis against *Plasmodium falciparum*, Clinical Infectious Diseases 36, 541–549 (2003).

33). G. D. Shanks, A. J. Oloo, G. M. Aleman, C. Ohrt, F. W. Klotz, D. Braitman, J. Horton, R. Brueckner, A new primaquine analogue, tafenoquine (WR 238605), for prophylaxis against *Plasmodium falciparum* malaria, Clinical Infectious Diseases 33, 1968–1974 (2001).

34). Lacerda MVG, Llanos-Cuentas A, Krudsood S, Lon C, Saunders DL, Mohammed R, Yilma D, Batista Pereira D, Espino FEJ, Mia RZ, Chuquiyauri R, Val F, Casapía M, Monteiro WM, Brito MAM, Costa MRF, Buathong N, Noedl H, Diro E, Getie S, Wubie KM, Abdissa A, Zeynudin A, Abebe C, Tada MS, Brand F, Beck HP, Angus B, Duparc S, Kleim JP, Kellam LM, Rousell VM, Jones SW, Hardaker E, Mohamed K, Clover DD, Fletcher K, Breton JJ, Ugwuegbulam CO, Green JA, Koh GCKW. Single-Dose Tafenoquine to Prevent Relapse of *Plasmodium vivax* Malaria. N. Eng.l J. Med. 17;380(3) (2019).

35). D. S. Walsh, C. Eamsila, T. Sasiprapha, S. Sangkharomya, P. Khaewsathien, P. Supakalin, D. B. Tang, P. Jarasrumgsichol, C. Cherdchu, M. D. Edstein, K. H. Rieckmanm, T. G. Brewer, Efficacy of monthly tafenoquine for prophylaxis of *Plasmodium vivax* and multidrug-resistant *P*. *falciparum* malaria, Journal of Infectious Diseases 190, 1456–1463 (2004).

36). H. Shiraki, M. P. Kozar, V. Melendez, T. H. Hudson, C. Ohrt, A. J. Magill, A. J. Lin, Antimalarial activity of novel 5-aryl-8-aminoquinoline derivatives, Journal of Medicinal Chemistry 54, 131–142 (2011).

37). N. Ponsa, J. Sattabongkot, P. Kittayapong, N. Eikarat, R. E. Coleman, Transmission-blocking activity of tafenoquine (WR-238605) and artelinic acid against naturally circulating strains of *Plasmodium vivax* in Thailand., The American Journal of Tropical Medicine and Hygiene 69, 542–547 (2003).

38). S. Tasai, T. Saiwichai, M. Kaewthamasorn, S. Tiawsirisup, P. Buddhirakkul, S. Chaichalotornkul, S. Pattaradilokrat, Artesunate-tafenoquine combination therapy promotes clearance and abrogates transmission of the avian malaria parasite *Plasmodium gallinaceum*, Veterinary Parasitology 233, 97–106 (2017).

39). Stone W, Mahamar A, Smit MJ, Sanogo K, Sinaba Y, Niambele SM, Sacko A, Keita S, Dicko OM, Diallo M, Maguiraga SO, Samake S, Attaher O, Lanke K, Ter Heine R, Bradley J, McCall MBB, Issiaka D, Traore SF, Bousema T, Drakeley C, Dicko A. Single low-dose tafenoquine combined with dihydroartemisinin-piperaquine to reduce Plasmodium falciparum transmission in Ouelessebougou, Mali: a phase 2, single-blind, randomised clinical trial. Lancet Microbe. 3(5):e336–e347. (2002).

40). Vanheer LN, Ramjith J, Mahamar A, Smit MJ, Lanke K, Roh ME, Sanogo K, Sinaba Y, Niambele SM, Diallo M, Maguiraga SO, Keita S, Samake S, Youssouf A, Diawara H, Traore SF, Gosling R, Brown JM, Drakeley C, Dicko A, Stone W, Bousema T. The transmission blocking activity of artemisinin-combination, non-artemisinin, and 8-aminoquinoline antimalarial therapies: A pooled analysis of individual participant data. PLoS Med.; 22(8):e1004683. (2025).

41). Webster R, Mitchell H, Peters JM, Heunis J, O’Neill B, Gower J, Lynch S, Jennings H, Amante FH, Llewellyn S, Marquart L, Potter AJ, Birrell GW, Edstein MD, Shanks GD, McCarthy JS, Barber BE. Transmission Blocking Activity of Low-dose Tafenoquine in Healthy Volunteers Experimentally Infected With *Plasmodium falciparum*, Clinical Infectious Diseases, 76, 3, (2003)

42). L. M. Upton, P. M. Brock, T. S. Churcher, A. C. Ghani, P. W. Gething, M. J. Delves, K. A. Sala, D. Leroy, R. E. Sinden, A. M. Blagborough, Lead clinical and preclinical antimalarial drugs can significantly reduce sporozoite transmission to vertebrate populations, Antimicrobial Agents and Chemotherapy 59, 490–497 (2015).

43). Baragaña B, Hallyburton I, Lee MC, Norcross NR, Grimaldi R, Otto TD, Proto WR, Blagborough AM, Meister S, Wirjanata G, Ruecker A, Upton LM, Abraham TS, Almeida MJ, Pradhan A, Porzelle A, Luksch T, Martínez MS, Luksch T, Bolscher JM, Woodland A, Norval S, Zuccotto F, Thomas J, Simeons F, Stojanovski L, Osuna-Cabello M, Brock PM, Churcher TS, Sala KA, Zakutansky SE, Jiménez-Díaz MB, Sanz LM, Riley J, Basak R, Campbell M, Avery VM, Sauerwein RW, Dechering KJ, Noviyanti R, Campo B, Frearson JA, Angulo-Barturen I, Ferrer-Bazaga S, Gamo FJ, Wyatt PG, Leroy D, Siegl P, Delves MJ, Kyle DE, Wittlin S, Marfurt J, Price RN, Sinden RE, Winzeler EA, Charman SA, Bebrevska L, Gray DW, Campbell S, Fairlamb AH, Willis PA, Rayner JC, Fidock DA, Read KD, Gilbert IH. A novel multiple-stage antimalarial agent that inhibits protein synthesis. Nature. 18;522(7556):315–20. (2015).

44). Ashley EA, Phyo AP. Drugs in Development for Malaria. Drugs; 78(9):861–879. doi: 10.1007/s40265-018-0911-9 (2018).

45). Ruecker A, Mathias DK, Straschil U, Churcher TS, Dinglasan RR, Leroy D, Sinden RE, Delves MJ. A male and female gametocyte functional viability assay to identify biologically relevant malaria transmission-blocking drugs. Antimicrob Agents Chemother.;58(12):7292–302 (2014).

46). Burgess, R. W., & Bray, R. S. The effect of a single dose of primaquine on the gametocytes, gametogony and sporogony of *Laverania falciparum*. Bulletin of the World Health Organization, 24(4-5), 451. (1961).

47). Mahamar A, Vanheer LN, Smit MJ, Sanogo K, Sinaba Y, Niambele SM, Diallo M, Dicko OM, Diarra RS, Maguiraga SO, Youssouf A, Sacko A, Keita S, Samake S, Dembele A, Teelen K, Dicko Y, Traore SF, Dondorp A, Drakeley C, Stone W, Dicko A. Artemether-lumefantrine-amodiaquine or artesunate-amodiaquine combined with single low-dose primaquine to reduce Plasmodium falciparum malaria transmission in Ouélessébougou, Mali: a five-arm, phase 2, single-blind, randomised controlled trial. Lancet Microbe. 6(2):100966. (2025).

48). Vantaux A, Kim S, Piv E, Chy S, Berne L, Khim N, Lek D, Siv S, Mukaka M, Taylor WR, Ménard D. Significant Efficacy of a Single Low Dose of Primaquine Compared to Stand-Alone Artemisinin Combination Therapy in Reducing Gametocyte Carriage in Cambodian Patients with Uncomplicated Multidrug-Resistant *Plasmodium falciparum* Malaria. Antimicrob. Agents Chemother. 64:10.1128. (2020)

49). Burrows, J.N., Hooft van Huijsduijnen, R., Möhrle, J.J, Oeuvray C, Wells TN. Designing the next generation of medicines for malaria control and eradication. Malar. J. 12, 187 (2013).

50). Talman AM, Domarle O, McKenzie FE, Ariey F, Robert V. Gametocytogenesis: the puberty of *Plasmodium falciparum*. Malar. J. 14;3:24. (2004).

51). P. W. Gething, D. L. Smith, A. P. Patil, A. J. Tatem, R. W. Snow, S. I. Hay, Climate change and the global malaria recession, Nature 465, 342–345 (2010).

52). B. M. Bolker, M. E. Brooks, C. J. Clark, S. W. Geange, J. R. Poulsen, M. H. H. Stevens, J. S. S. White, Generalized linear mixed models: a practical guide for ecology and evolution, Trends in Ecology and Evolution 24, 127–135 (2009).

53). M. B. Jiménez-Díaz, T. Mulet, S. Viera, V. Gómez, H. Garuti, J. Ibáñez, A. Alvarez-Doval, L. D. Shultz, A. Martínez, D. Gargallo-Viola, I. Angulo-Barturen, Improved murine model of malaria using *Plasmodium falciparum* competent strains and non-myelodepleted NOD-scid IL2Rγnull mice engrafted with human erythrocytes, Antimicrobial Agents and Chemotherapy 53, 4533–4536 (2009).

