## Supplemental data for "Comparing the transmission blocking efficacy of Primaquine and Tafenoquine with *in vivo* pre-clinical models"

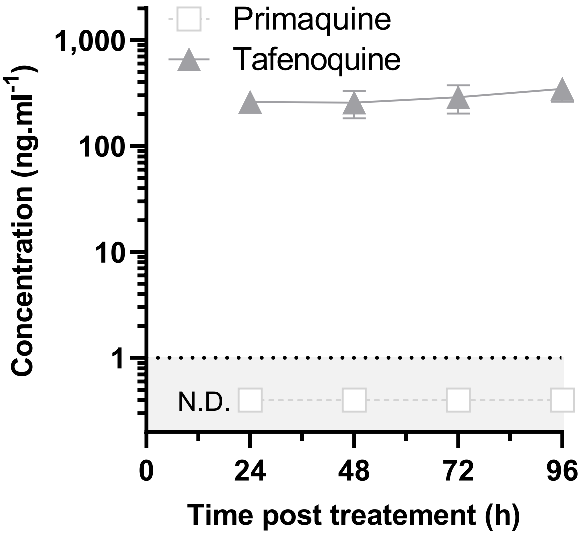

**Figure S1: Individual blood concentration-time profile after a single dose oral administration of Primaquine (3.35 mg.kg$^{-1}$) (A) and Tafenoquine (3.41 mg.kg$^{-1}$) (B) in mouse-to-mouse transmission model infected with *P. berghei* parasites.** The lower limit of quantification is indicated by the dashed line. Mean +/- SD for 4 samples from 2 mice. The levels of primaquine were below the lower limit of quantification and are artificially represented here (dashed line; N.D., non-determined).
Raw data are outlined in Table S1.

**Figure S1**

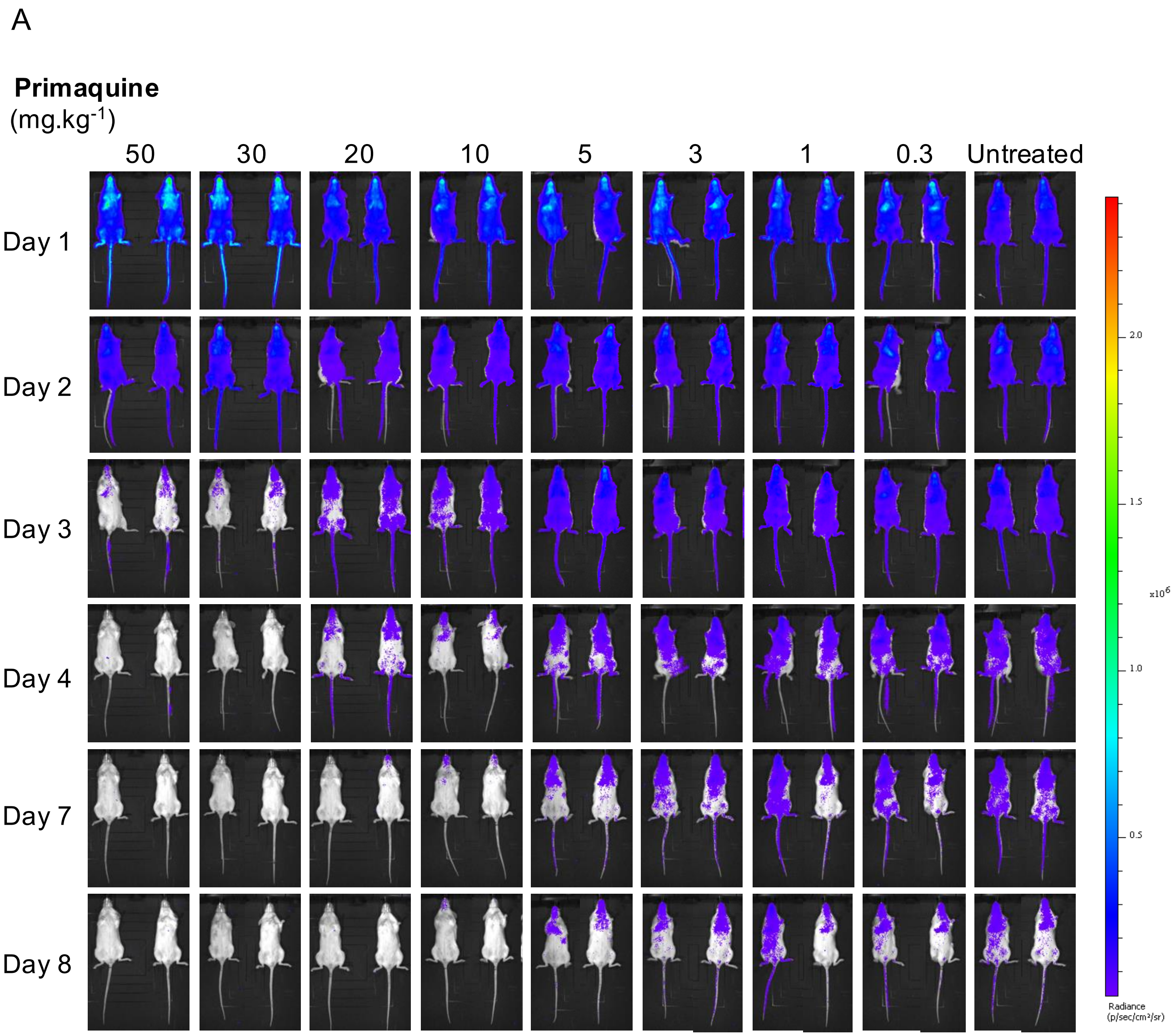
A
Primaquine
(mg.kg-1)
50
30
20
10
5
3
1
0.3
Untreated
Day 1
Day 2
Day 3
Day 4
Day 7
Day 8
2.0
1.5
1.0
0.5
x10^6
Radiance
(p/sec/cm²/sr)

**Figure S2**

B

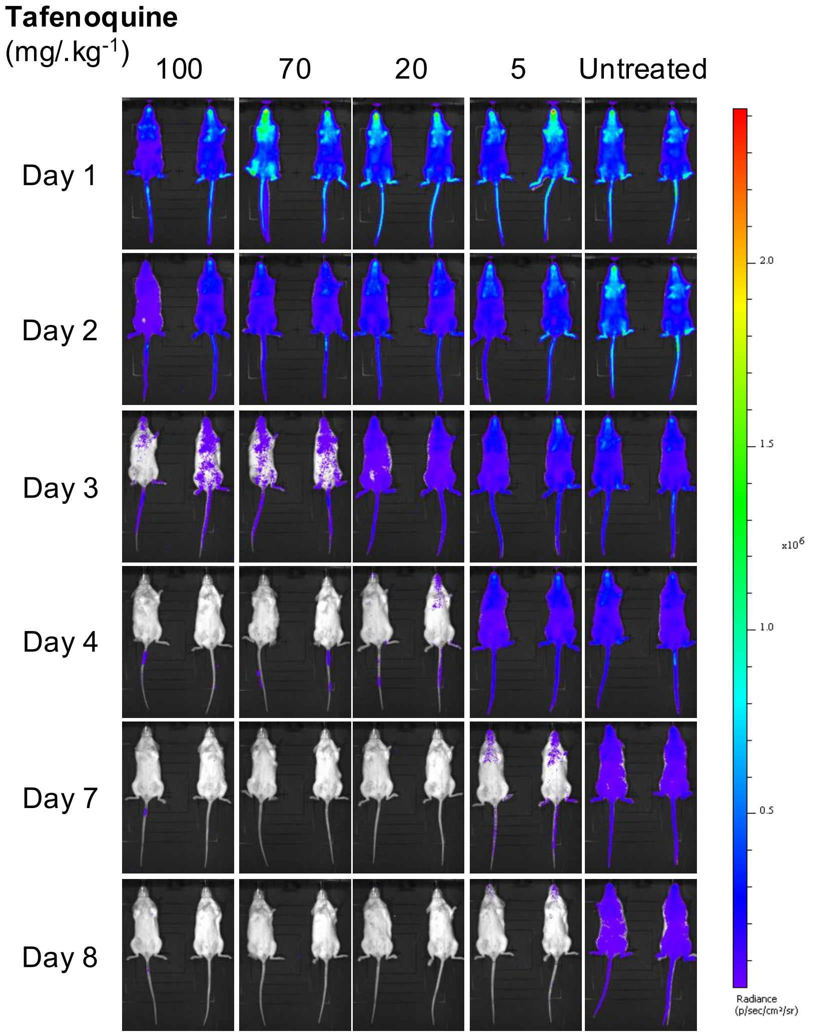

**Figure S2: Qualitative therapeutic efficacy against *P. falciparum* NF54/iGP_RE9H gametocytes determined by bioluminescence in NSG humanized mice.** Representative ventral images of mice infected with $2x10^8$ stages IV/V gametocytes. Mice were infected on day 0, treated on day 1 with various doses of Primaquine (A) or Tafenoquine (B) and compared to untreated control mice. Pseudo color heat-maps indicate intensity of detected bioluminescence from low (blue) to high (red).

**Figure S2**

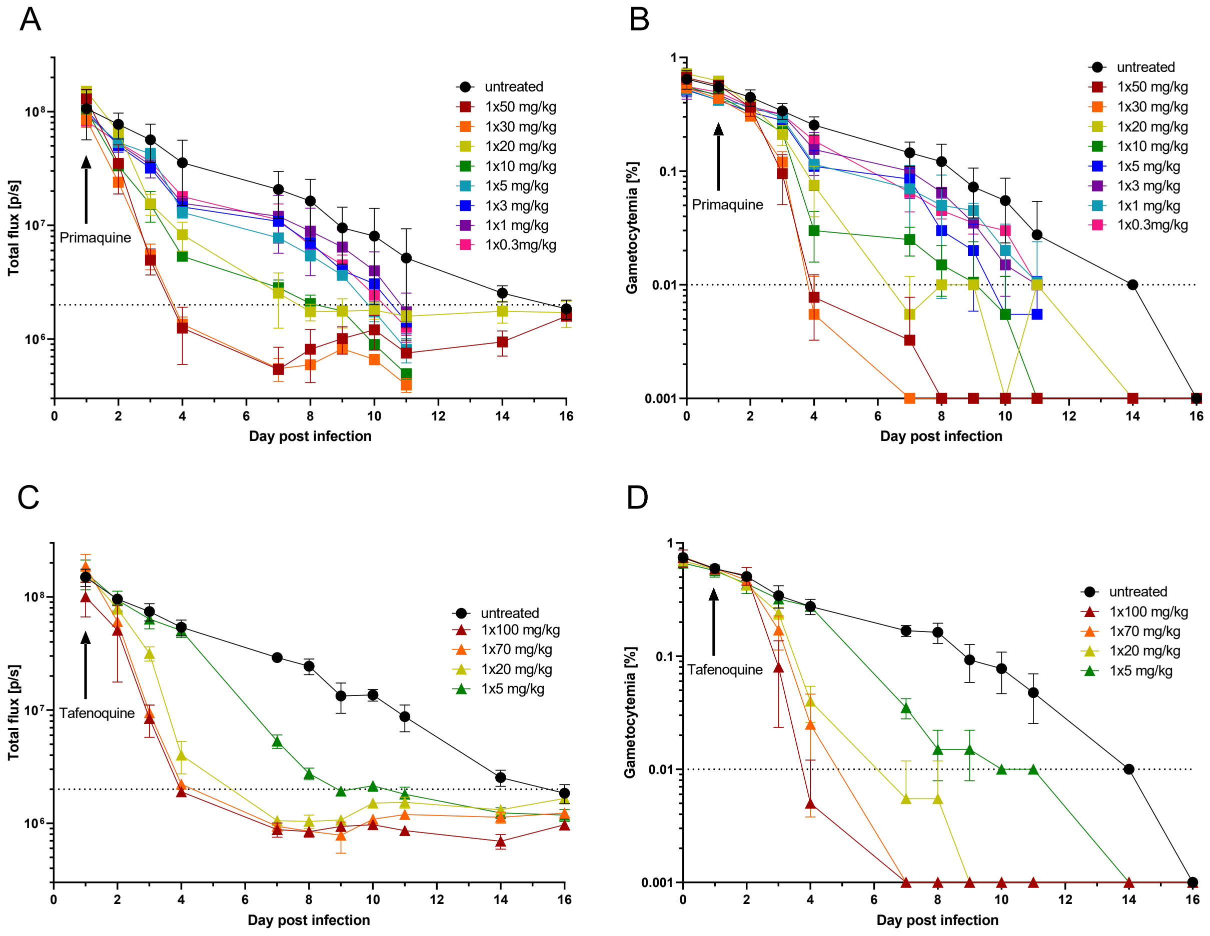

**Figure S3: Quantitative therapeutic efficacy against *P. falciparum* NF54/iGP_RE9H gametocytes evaluated with two distinct read-outs.** Efficacy of Primaquine determined by bioluminescence (A) and microscopic gametocyte count (B); and efficacy of Tafenoquine determined by bioluminescence (C) and microscopic gametocyte count (D). Mice were infected *P. falciparum* NF54/iGP_RE9H gametocytes stages IV/V on day 0 and treated on day 1 with various doses of Primaquine or Tafenoquine. The arrow indicates the day of treatment. The lower limit of quantification is indicated by the dashed line. Mean +/- SD of at least 2 mice.

**Figure S3**

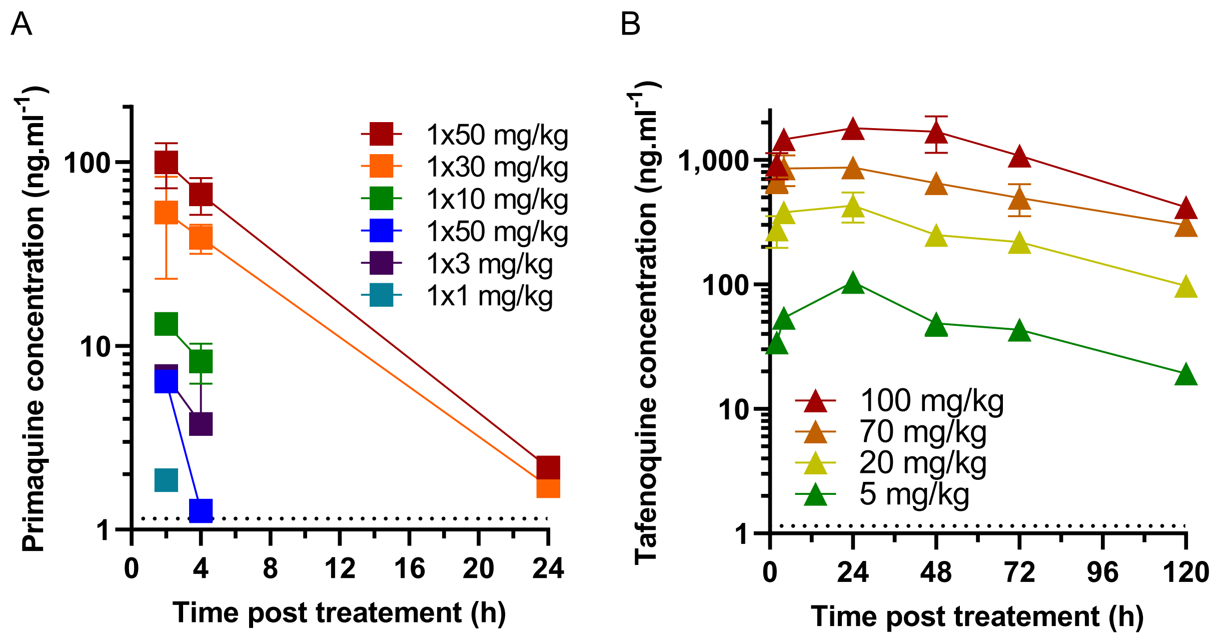

**Figure S4: Individual blood concentration-time profile after oral administration of Primaquine (A) and Tafenoquine (B) in NSG humanized mice infected with *P. falciparum* NF54/iGP_RE9H gametocytes.** The lower limit of quantification is indicated by the dashed line. Mean +/- SD of at least 2 mice.
Raw data and calculated pharmacokinetic parameters are display in Table S4.

**Figure S4**

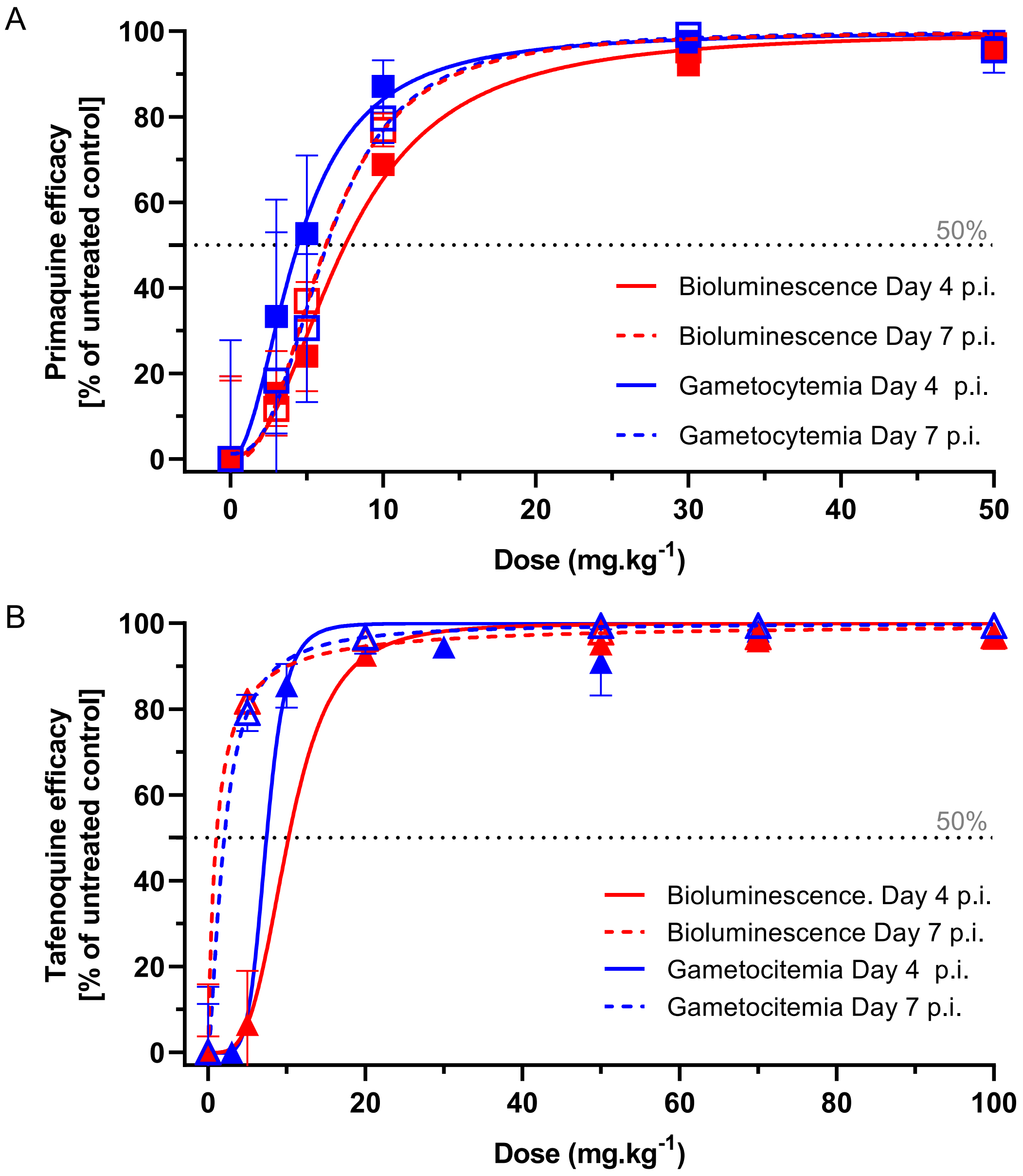

C

**$ED_{50}$ (mg.kg$^{-1}$)**

| | **Primaquine** | | **Tafenoquine** | |
|---|---|---|---|---|
| *Read-out* | *Bioluminescence* | *Gametocytemia* | *Bioluminescence* | *Gametocytemia* |
| Day 4 p.i. | 7.5 ($R^2$ = 0.9417) | 6.4 ($R^2$ = 0.8516) | 10 ($R^2$ = 0.9667) | 7.4 ($R^2$ = 0.9657) |
| Day 7 p.i. | 6.2 ($R^2$ = 0.9498) | 4.4 ($R^2$ = 0.9057) | 1. 0 ($R^2$ = 0.9974) | 2. 0 ($R^2$ = 0.9844) |

**Figure S5: Efficacy of Primaquine (A) and Tafenoquine (B) compared to untreated control against *P. falciparum* NF54/iGP_RE9H stages IV/V gametocytes in NSG humanized mice.** Efficacy was expressed based on the measured bioluminescence (red) or gametocytemia count (blue), compared to the untreated control mice, on Day 4 (full line) and Day 7 (dashed line) post-infection (p.i.) Non-linear regressions were performed in GraphPad Prism (variable slope, four parameters). Mean +/- SD of at least 2 mice. The dotted line represent 50% of efficacy. (C) Summary of the $ED_{50}$ values obtain in each condition and timepoint.

**Figure S5**

### Table S1: Oocyst counts and calculations for Primaquine titration

| *Oocyst scores* | Primaquine concentration (mg/kg) | | | | | | | | | | | | | | | | | | | | | | | | | |
|---|---|---|---|---|---|---|---|---|---|---|---|---|---|---|---|---|---|---|---|---|---|---|---|---|---|---|
| | 0 | 0 | 0 | 0 | 0 | 0 | 0 | 0.25 | 1.25 | 1.5 | 1.75 | 2 | 2.5 | 2.75 | 2.9 | 3 | 3 | 3.1 | 3.2 | 3.3 | 3.4 | 3.5 | 4 | 5 | 6 | 12 |
| Experiment 1 | 81 | 131 | 38 | 270 | 139 | 43 | 0 | 61 | 0 | 62 | 59 | 31 | 39 | 18 | 24 | 0 | 40 | 45 | 0 | 0 | 6 | 0 | 0 | 0 | 0 | 0 |
| Experiment 2 | 5 | 284 | 58 | 239 | 28 | 193 | 67 | 94 | 0 | 85 | 48 | 93 | 48 | 11 | 19 | 57 | 55 | 4 | 0 | 1 | 0 | 0 | 0 | 0 | 0 | 0 |
| Experiment 3 | 45 | 55 | 69 | 142 | 62 | 17 | 16 | 112 | 51 | 100 | 4 | 42 | 79 | 15 | 36 | 1 | 9 | 9 | 6 | 4 | 1 | 23 | 0 | 0 | 0 | 0 |
| Experiment 4 | 45 | 146 | 133 | 1 | 3 | 51 | 0 | 34 | 0 | 0 | 69 | 1 | 23 | 12 | 55 | 0 | 16 | 57 | 0 | 3 | 0 | 0 | 0 | 0 | 0 | 0 |
| Experiment 5 | 58 | 127 | 23 | 42 | 155 | 87 | 13 | 1 | 26 | 46 | 0 | 91 | 34 | 9 | 0 | 0 | 31 | 39 | 1 | 0 | 14 | 0 | 0 | 0 | 0 | 0 |
| Experiment 6 | 154 | 93 | 76 | 48 | 51 | 8 | 3 | 139 | 7 | 182 | 0 | 75 | 22 | 44 | 89 | 0 | 14 | 54 | 21 | 4 | 0 | 10 | 0 | 0 | 0 | 0 |
| Experiment 7 | 41 | 122 | 0 | 3 | 62 | 0 | 41 | 9 | 33 | 58 | 0 | 20 | 21 | 12 | 5 | 0 | 21 | 57 | 2 | 0 | 1 | 10 | 0 | 0 | 0 | 0 |
| | 110 | | 44 | 0 | 36 | 78 | 39 | 159 | 191 | 18 | 21 | 6 | 3 | 36 | 14 | 1 | 23 | 54 | 0 | 0 | 0 | 3 | 0 | 0 | 0 | 0 |
| | 93 | | 36 | 170 | 0 | 12 | 54 | 27 | 0 | 20 | 53 | 46 | 62 | 17 | 10 | 0 | 0 | 46 | 33 | 0 | 0 | 5 | 0 | 0 | 0 | 0 |
| | 92 | | 101 | 0 | 105 | 155 | 0 | 0 | 140 | 61 | 21 | 0 | 5 | 2 | 21 | 0 | 7 | 21 | 0 | 5 | 0 | 0 | 0 | 0 | 0 | 0 |
| | 30 | | 165 | 10 | 13 | 0 | 186 | 33 | 27 | 52 | 4 | 3 | 0 | 17 | 0 | 0 | 64 | 37 | 4 | 0 | 4 | 5 | 0 | | 0 | 0 |
| | 43 | | 52 | 0 | 5 | 6 | 0 | 11 | 2 | 34 | 14 | 0 | 17 | 30 | 0 | 0 | 18 | 63 | 19 | 0 | 0 | 7 | 0 | | 0 | 0 |
| | 49 | | 102 | 126 | 26 | 0 | 9 | 0 | 167 | 10 | 0 | 70 | 136 | 61 | 32 | 5 | 30 | 35 | 0 | 1 | 0 | 0 | 0 | | 0 | 0 |
| | 42 | | 95 | 6 | 15 | 8 | 0 | 12 | 0 | 32 | 49 | 14 | 7 | 0 | 0 | 1 | 15 | 31 | 0 | 0 | 10 | 0 | 0 | | 0 | 0 |
| | 79 | | 72 | 0 | 0 | 0 | 16 | 40 | 64 | 0 | 129 | 52 | 11 | 21 | 22 | 0 | 30 | 28 | 0 | 0 | 0 | 0 | 0 | | 0 | 0 |
| | 54 | | 56 | 2 | 89 | 1 | 16 | 55 | 16 | 17 | 3 | 0 | 0 | 2 | 22 | 0 | 117 | 80 | 2 | 0 | 6 | 0 | 0 | | 0 | 0 |
| | 61 | | 51 | 69 | 103 | 53 | 0 | 26 | 11 | 244 | 29 | 89 | 45 | 0 | 1 | 0 | 9 | 39 | 0 | 0 | 3 | 4 | 0 | | 0 | 0 |
| | 57 | | 115 | 13 | 187 | 21 | 245 | 0 | 88 | 83 | 0 | 10 | 12 | 18 | 0 | 2 | 4 | 47 | 32 | 0 | 0 | 2 | 0 | | 0 | 0 |
| | 59 | | 133 | 41 | 22 | 2 | 0 | 17 | 35 | 148 | 7 | 83 | 0 | 26 | 20 | 0 | 15 | 35 | 10 | 0 | 13 | 0 | 0 | | 0 | 0 |
| | 33 | | 83 | 0 | 3 | 11 | 250 | 3 | 82 | 161 | 10 | 0 | 78 | 21 | 6 | 1 | 35 | 22 | 4 | 0 | 6 | 12 | 0 | | 0 | 0 |
| | 47 | | 69 | 149 | 7 | 73 | 19 | 0 | 65 | | | | | 2 | 3 | 0 | 19 | 39 | 0 | 0 | 3 | 0 | | | 0 | 0 |
| | 64 | | 27 | 4 | 0 | 3 | 8 | 64 | 27 | | | | | 0 | 2 | 0 | 40 | 33 | 1 | 0 | 17 | 0 | | | 0 | 0 |
| | 69 | | 94 | 51 | 98 | 1 | 0 | 56 | 36 | | | | | 8 | 6 | 15 | 0 | 38 | 0 | 0 | 16 | 0 | | | 0 | 0 |
| | 70 | | 131 | 208 | 96 | 25 | 23 | 0 | 36 | | | | | 4 | 0 | 1 | 76 | 11 | 0 | 0 | 8 | 12 | | | 0 | 0 |
| | 45 | | 39 | 92 | 86 | 30 | 6 | 0 | 0 | | | | | 0 | 2 | 0 | 11 | 30 | 0 | 1 | 0 | 0 | | | 0 | 0 |
| | 109 | | 75 | 13 | 49 | 7 | 0 | 15 | 8 | | | | | 37 | 43 | 0 | 31 | 41 | 0 | 1 | 10 | 0 | | | 0 | 0 |
| | 39 | | 15 | 222 | 67 | 283 | 45 | 63 | 2 | | | | | 5 | 64 | 2 | 66 | 22 | 4 | 2 | 2 | 3 | | | 0 | 0 |
| | 36 | | 36 | 0 | 53 | 65 | 1 | 25 | 4 | | | | | 0 | 0 | 0 | 43 | 17 | 0 | 0 | 3 | 2 | | | 0 | 0 |
| | 9 | | 77 | 0 | 0 | 2 | 89 | 22 | 8 | | | | | 7 | 43 | 2 | 4 | 50 | 2 | 0 | 14 | 0 | | | 0 | 0 |
| | 15 | | 31 | 0 | 93 | 2 | 6 | 0 | 11 | | | | | 2 | 9 | 0 | 0 | 25 | 5 | 0 | 0 | 0 | | | 0 | 0 |
| | 79 | | 50 | 100 | 8 | 22 | 9 | 99 | 10 | | | | | 21 | 9 | 1 | 25 | 8 | 32 | 1 | 0 | 0 | | | 0 | 0 |
| | 28 | | 44 | 0 | 68 | 9 | 0 | 0 | 0 | | | | | 41 | 0 | 0 | 27 | 7 | 6 | 0 | 0 | 4 | | | 0 | 0 |
| | 29 | | 2 | 82 | 58 | 28 | 112 | 21 | 144 | | | | | 0 | 10 | 0 | 2 | 9 | 7 | 0 | 10 | 0 | | | 0 | 0 |
| | 64 | | 83 | 0 | 47 | 139 | 88 | 112 | 65 | | | | | 3 | 7 | 0 | 12 | 26 | 0 | 0 | 0 | 29 | | | 0 | 0 |
| | 36 | | 89 | 127 | 53 | 106 | 2 | 48 | 55 | | | | | 0 | 0 | 2 | 25 | 85 | 0 | 3 | 0 | 2 | | | 0 | 0 |
| | 42 | | 105 | 0 | 0 | 27 | 120 | 32 | 1 | | | | | 57 | 14 | 2 | 61 | 44 | 0 | 0 | 0 | 0 | | | 0 | 0 |
| | 7 | | 43 | 0 | 0 | 45 | 19 | 13 | 0 | | | | | 2 | 0 | 0 | 3 | 36 | 0 | 0 | 0 | 0 | | | 0 | 0 |
| | 67 | | 39 | 0 | 154 | 0 | 0 | 0 | 59 | | | | | 0 | 6 | 0 | 29 | 42 | 0 | 9 | 15 | 6 | | | 0 | 0 |
| | 88 | | 0 | 228 | 0 | 227 | 31 | 79 | 0 | | | | | 9 | 1 | 0 | 35 | 55 | 26 | 0 | 0 | 28 | | | 0 | 0 |
| | 27 | | 51 | 161 | 0 | 104 | 1 | 37 | 0 | | | | | 66 | 32 | 0 | 35 | 18 | 0 | 0 | 0 | 0 | | | 0 | 0 |
| | 39 | | 0 | 0 | 3 | 53 | 93 | 35 | 11 | | | | | 23 | 11 | 11 | 15 | 4 | 19 | 0 | 3 | 0 | | | 0 | 0 |
| | 13 | | 34 | 0 | 85 | 137 | 152 | 194 | 69 | | | | | 11 | 0 | 1 | 0 | 6 | 14 | 0 | 3 | 0 | | | 0 | 0 |
| | 84 | | 0 | 6 | 0 | 152 | 49 | 1 | 0 | | | | | 0 | 1 | 0 | 7 | 56 | 0 | 0 | 0 | 0 | | | 0 | 0 |
| | 54 | | 32 | 0 | 27 | 14 | 29 | 51 | 23 | | | | | 6 | 4 | 1 | 42 | 19 | 0 | 2 | 0 | 0 | | | 0 | 0 |
| | 118 | | 109 | 51 | 156 | 0 | 11 | 0 | 51 | | | | | 0 | 7 | 0 | 16 | 8 | 0 | 0 | 0 | 0 | | | 0 | 0 |
| | 91 | | 59 | 112 | 38 | 0 | 0 | 43 | 49 | | | | | 42 | 10 | 0 | 0 | 19 | 1 | 0 | 0 | 4 | | | 0 | 0 |
| | | | 88 | 4 | 24 | 0 | 1 | 158 | 88 | | | | | 2 | 18 | 0 | 29 | 9 | 0 | 5 | 15 | 3 | | | 0 | 0 |
| | | | 38 | 7 | 35 | 100 | 17 | 1 | 11 | | | | | 52 | 3 | 0 | 1 | 22 | 14 | 3 | 0 | 4 | | | 0 | 0 |
| | | | 33 | 0 | 80 | 16 | 42 | 21 | 0 | | | | | 3 | 0 | 2 | 11 | 9 | 0 | 0 | 0 | 0 | | | 0 | 0 |
| | | | 53 | 6 | 91 | 16 | 25 | 65 | 33 | | | | | 28 | 1 | 0 | 4 | 24 | 2 | 0 | 0 | 0 | | | 0 | 0 |
| | | | | 0 | 26 | 13 | 78 | 0 | 20 | | | | | | | 1 | | | | | | | | | 0 | 0 |
| | | | | 54 | 3 | 0 | 63 | 77 | 79 | | | | | | | 0 | | | | | | | | | 0 | 0 |
| | | | | 82 | 13 | 28 | 24 | 0 | 1 | | | | | | | 2 | | | | | | | | | 0 | 0 |
| | | | | 83 | 0 | 0 | 130 | 16 | 52 | | | | | | | | | | | | | | | | 0 | 0 |
| | | | | 160 | 2 | 18 | 0 | 4 | 2 | | | | | | | | | | | | | | | | 0 | 0 |
| | | | | 129 | 12 | 13 | | 177 | 16 | | | | | | | | | | | | | | | | 0 | 0 |
| | | | | 11 | 25 | 97 | | 8 | 3 | | | | | | | | | | | | | | | | | |
| | | | | | 32 | 9 | | 62 | 33 | | | | | | | | | | | | | | | | | |
| | | | | | 4 | 7 | | 0 | 38 | | | | | | | | | | | | | | | | | |
| | | | | | 3 | 68 | | 25 | 1 | | | | | | | | | | | | | | | | | |
| | | | | | 10 | 133 | | 50 | 1 | | | | | | | | | | | | | | | | | |
| | | | | | 10 | | | 0 | 0 | | | | | | | | | | | | | | | | | |
| | | | | | 52 | | | 1 | 31 | | | | | | | | | | | | | | | | | |
| | | | | | | | | 65 | 101 | | | | | | | | | | | | | | | | | |
| | | | | | | | | 211 | 267 | | | | | | | | | | | | | | | | | |
| | | | | | | | | 47 | 102 | | | | | | | | | | | | | | | | | |
| | | | | | | | | 10 | 14 | | | | | | | | | | | | | | | | | |
| | | | | | | | | | 0 | | | | | | | | | | | | | | | | | |
| | | | | | | | | | 86 | | | | | | | | | | | | | | | | | |
| | | | | | | | | | 14 | | | | | | | | | | | | | | | | | |
| | | | | | | | | | 8 | | | | | | | | | | | | | | | | | |
| | | | | | | | | | 144 | | | | | | | | | | | | | | | | | |

*Oocyst scores*

| | | | | | | | | | | | | | | | | | | | | | | | | | | |
|---|---|---|---|---|---|---|---|---|---|---|---|---|---|---|---|---|---|---|---|---|---|---|---|---|---|---|
| Uninfected | 0 | 0 | 4 | 18 | 10 | 10 | 13 | 14 | 14 | 2 | 5 | 4 | 3 | 10 | 12 | 33 | 5 | 0 | 26 | 35 | 27 | 29 | 20 | 10 | 56 | 56 |
| Infected | 46 | 7 | 46 | 39 | 53 | 51 | 42 | 53 | 58 | 18 | 15 | 16 | 17 | 40 | 38 | 20 | 45 | 50 | 24 | 15 | 23 | 21 | 0 | 0 | 0 | 0 |

### Table S1: Oocyst counts and calculations for Primaquine titration (continued)

| | Primaquine concentration (mg/kg) | | | | | | | | | | | | | | | | | | | | | | | | | |
|---|---|---|---|---|---|---|---|---|---|---|---|---|---|---|---|---|---|---|---|---|---|---|---|---|---|---|
| | 0 | 0 | 0 | 0 | 0 | 0 | 0 | 0.25 | 1.25 | 1.5 | 1.75 | 2 | 2.5 | 2.75 | 2.9 | 3 | 3 | 3.1 | 3.2 | 3.3 | 3.4 | 3.5 | 4 | 5 | 6 | 12 |

***Oocyst scores***

| | | | | | | | | | | | | | | | | | | | | | | | | | | |
|---|---|---|---|---|---|---|---|---|---|---|---|---|---|---|---|---|---|---|---|---|---|---|---|---|---|---|
| **Uninfected** | 0 | 0 | 4 | 18 | 10 | 10 | 13 | 14 | 14 | 2 | 5 | 4 | 3 | 10 | 12 | 33 | 5 | 0 | 26 | 35 | 27 | 29 | 20 | 10 | 56 | 56 |
| **Infected** | 46 | 7 | 46 | 39 | 53 | 51 | 42 | 53 | 58 | 18 | 15 | 16 | 17 | 40 | 38 | 20 | 45 | 50 | 24 | 15 | 23 | 21 | 0 | 0 | 0 | 0 |
| **Mean Intensity** | 57 | 137 | 61 | 58 | 44 | 46 | 41 | 42 | 39 | 71 | 26 | 36 | 32 | 16 | 14 | 2 | 24 | 32 | 5 | 1 | 4 | 4 | 0 | 0 | 0 | 0 |
| **Std. Deviation** | 31 | 72 | 39 | 76 | 47 | 62 | 59 | 51 | 52 | 66 | 33 | 36 | 35 | 18 | 19 | 8 | 23 | 20 | 9 | 2 | 5 | 7 | 0 | 0 | 0 | 0 |
| **Std. Error** | 5 | 27 | 5 | 10 | 6 | 8 | 8 | 6 | 6 | 15 | 7 | 8 | 8 | 3 | 3 | 1 | 3 | 3 | 1 | 0 | 1 | 1 | 0 | 0 | 0 | 0 |
| **Prevalence** | 100% | 100% | 92% | 68% | 84% | 84% | 76% | 79% | 81% | 90% | 75% | 80% | 85% | 80% | 76% | 38% | 90% | 100% | 48% | 30% | 46% | 42% | 0% | 0% | 0% | 0% |

***Transmission blocking efficacy***

| | | | | | | | | | | | | | | | | | | | | | | | | | | |
|---|---|---|---|---|---|---|---|---|---|---|---|---|---|---|---|---|---|---|---|---|---|---|---|---|---|---|
| **Intensity (%)** | N/A | N/A | N/A | N/A | N/A | N/A | N/A | 27 | 11 | 48 | 81 | 73 | 77 | 74 | 78 | 96 | 60 | 47 | 91 | 99 | 94 | 94 | 100 | 100 | 100 | 100 |
| **Prevalence (%)** | N/A | N/A | N/A | N/A | N/A | N/A | N/A | -16 | 4 | 10 | 25 | 20 | 15 | 13 | 17 | 62 | 2 | -9 | 48 | 67 | 50 | 54 | 100 | 100 | 100 | 100 |

### Table S2: Transmission blocking efficacy of Primaquine and Tafenoquine in DFA

| *Oocyst scores* | 0h | | 24h | | 48h | | 72h | | 96h | |
|---|---|---|---|---|---|---|---|---|---|---|
| | Untreated | | Primaquine | Tafenoquine | Primaquine | Tafenoquine | Primaquine | Tafenoquine | Primaquine | Tafenoquine |
| Experiment 1 | 60 | 171 | 3 | 62 | 31 | 0 | 0 | 23 | 13 | 0 |
| Experiment 2 | 12 | 152 | 0 | 94 | 0 | 0 | 44 | 9 | 26 | 0 |
| Experiment 3 | 86 | 67 | 13 | 27 | 7 | 0 | 0 | 0 | 27 | 12 |
| | 92 | 132 | 2 | 45 | 0 | 0 | 23 | 0 | 51 | 2 |
| | 5 | 78 | 0 | 81 | 0 | 0 | 10 | 8 | 0 | 0 |
| | 11 | 95 | 0 | 116 | 0 | 0 | 11 | 0 | 10 | 0 |
| | 101 | 0 | 5 | 45 | 1 | 0 | 9 | 27 | 19 | 4 |
| | 15 | 139 | 0 | 81 | 2 | 0 | 26 | 23 | 41 | 0 |
| | 58 | 119 | 0 | 15 | 0 | 0 | 19 | 8 | 0 | 0 |
| | 1 | 98 | 20 | 0 | 8 | 9 | 69 | 0 | 53 | 17 |
| | 89 | 33 | 22 | 19 | 24 | 0 | 8 | 0 | 85 | 3 |
| | 35 | 40 | 6 | 35 | 0 | 0 | 124 | 0 | 26 | 0 |
| | 96 | 52 | 0 | 56 | 6 | 1 | 1 | 2 | 41 | 0 |
| | 77 | 44 | 0 | 12 | 0 | 0 | 0 | 17 | 0 | 0 |
| | 28 | 43 | 0 | 0 | 36 | 1 | 0 | 0 | 111 | 4 |
| | 37 | 110 | 0 | 11 | 15 | 0 | 91 | 4 | 42 | 0 |
| | 104 | 71 | 3 | 49 | 0 | 0 | 31 | 0 | 86 | 1 |
| | 0 | 107 | 0 | 31 | 0 | 0 | 17 | 0 | 0 | 0 |
| | 10 | 119 | 0 | 19 | 0 | 0 | 12 | 30 | 5 | 4 |
| | 55 | 83 | 4 | 0 | 0 | 0 | 39 | 0 | 0 | 0 |
| | 9 | 78 | 0 | 17 | 0 | 3 | 38 | 11 | 7 | 7 |
| | 120 | 93 | 10 | 35 | 1 | 6 | 0 | 9 | 5 | 0 |
| | 152 | | 0 | 31 | 0 | 7 | 0 | 1 | 0 | 0 |
| | 42 | | 3 | 21 | 14 | 0 | 0 | 7 | 15 | 5 |
| | 69 | | 24 | 6 | 36 | 6 | 4 | 0 | 58 | 0 |
| | 34 | | 5 | 61 | 0 | 0 | 23 | 0 | 11 | 1 |
| | 0 | | 0 | 47 | 0 | 0 | 0 | 0 | 17 | 0 |
| | 18 | | 1 | 90 | 0 | 0 | 20 | 26 | 41 | 8 |
| | 86 | | 0 | 44 | 18 | 0 | 36 | 0 | 0 | 0 |
| | 3 | | 3 | 66 | 5 | 0 | 0 | 0 | 12 | 0 |
| | 31 | | 1 | 64 | 85 | 0 | 0 | 0 | 51 | 0 |
| | 14 | | 9 | 42 | 5 | 0 | 0 | 8 | 74 | 0 |
| | 79 | | 0 | 2 | 10 | 0 | 0 | 0 | 0 | 3 |
| | 0 | | 0 | 0 | 24 | 0 | 3 | 18 | 0 | 0 |
| | 0 | | 8 | 14 | 0 | 0 | 87 | 22 | 0 | 0 |
| | 89 | | 19 | 82 | 8 | 0 | 34 | 3 | 0 | 0 |
| | 31 | | 8 | 51 | 12 | 0 | 0 | 0 | 13 | 0 |
| | 0 | | 15 | 74 | 1 | 0 | 16 | 0 | 0 | 0 |
| | 136 | | 2 | 22 | 0 | 0 | 2 | 7 | 0 | 0 |
| | 81 | | 0 | 38 | 31 | 0 | 0 | 8 | 26 | 0 |
| | 111 | | 0 | 66 | 17 | 0 | 3 | 0 | 35 | 0 |
| | 59 | | 0 | 82 | 27 | 0 | 22 | 0 | 5 | 0 |
| | 22 | | 0 | 0 | 8 | 0 | 0 | 0 | 37 | 0 |
| | 56 | | 0 | 39 | 0 | 0 | 64 | 0 | 18 | 5 |
| | 0 | | 19 | | 0 | 0 | 0 | 12 | 167 | 0 |
| | 1 | | 43 | | 0 | 0 | 0 | 15 | 27 | 0 |
| | 0 | | 0 | | 2 | 0 | 10 | 0 | 51 | 0 |
| | 97 | | 0 | | | 0 | 0 | 12 | 23 | 0 |
| | 115 | | 7 | | | 0 | 14 | 3 | 0 | 5 |
| | 12 | | 0 | | | 0 | 0 | 0 | 0 | 1 |
| | 6 | | | | | 0 | 72 | 0 | 15 | 0 |
| | 0 | | | | | 0 | 10 | 52 | 57 | 24 |
| | 3 | | | | | 0 | 0 | 34 | 57 | 8 |
| | | | | | | 0 | | 8 | 72 | 0 |
| | | | | | | 0 | | 0 | 17 | 0 |
| | | | | | | | | 0 | 0 | 0 |
| | | | | | | | | | 1 | 0 |
| | | | | | | | | | 7 | 8 |
| | | | | | | | | | 29 | 11 |
| | | | | | | | | | 57 | |
| | | | | | | | | | 14 | |

| *Oocyst scores* | | | | | | | | | | |
|---|---|---|---|---|---|---|---|---|---|---|
| **Uninfected** | 8 | 1 | 25 | 5 | 21 | 49 | 20 | 28 | 16 | 39 |
| **Infected** | 45 | 21 | 25 | 40 | 26 | 6 | 48 | 28 | 45 | 20 |
| **Mean Intensity** | 46.19 | 87.45 | 5.10 | 40.73 | 9.23 | 0.60 | 18.72 | 7.27 | 27.13 | 2.25 |
| **Std. Deviation** | 43.29 | 42.39 | 8.61 | 29.98 | 15.62 | 1.89 | 27.56 | 10.97 | 31.98 | 4.59 |
| **Std. Error** | 5.95 | 9.04 | 1.22 | 4.52 | 2.28 | 0.26 | 3.79 | 1.47 | 4.09 | 0.60 |
| **Prevalence** | 85% | 95% | 50% | 89% | 55% | 11% | 71% | 50% | 74% | 34% |

| *Transmission blocking efficacy* | | | | | | | | | |
|---|---|---|---|---|---|---|---|---|---|
| **Intensity (%)** | 100% | 7.6% | 60.9% | 13.8% | 0.9% | 28.0% | 10.9% | 40.6% | 3.4% |
| **Prevalence (%)** | 90% | 50% | 89% | 55% | 11% | 71% | 50% | 74% | 34% |

#### Table S2: Transmission blocking efficacy of Primaquine and Tafenoquine in DFA (continued)

| *Oocyst scores* | 0h | | 24h | | 48h | | 72h | | 96h | |
|---|---|---|---|---|---|---|---|---|---|---|
| | Untreated | | Primaquine | Tafenoquine | Primaquine | Tafenoquine | Primaquine | Tafenoquine | Primaquine | Tafenoquine |
| Experiment 1 | 0 | 97 | 0 | 55 | 62 | 0 | 3 | 0 | 2 | 0 |
| Experiment 2 | 52 | 50 | 32 | 26 | 31 | 14 | 6 | 5 | 31 | 58 |
| Experiment 3 | 75 | 11 | 13 | 10 | 44 | 3 | 20 | 6 | 7 | 13 |
| | 16 | 56 | 0 | 31 | 39 | 27 | 14 | 0 | 56 | 19 |
| | 0 | 59 | 6 | 12 | 31 | 0 | 9 | 37 | 1 | 0 |
| | 74 | 105 | 27 | 48 | 29 | 0 | 31 | 0 | 4 | 0 |
| | 0 | 43 | 21 | 9 | 28 | 0 | 24 | 0 | 5 | 10 |
| | 170 | 0 | 0 | 0 | 147 | 0 | 68 | 0 | 4 | 3 |
| | 78 | 14 | 0 | 0 | 42 | 0 | 94 | 6 | 0 | 26 |
| | 118 | 54 | 17 | 4 | 18 | 0 | 9 | 0 | 24 | 0 |
| | 2 | 0 | 3 | 103 | 42 | 0 | 10 | 1 | 5 | 0 |
| | 159 | 98 | 26 | 112 | 39 | 1 | 61 | 6 | 19 | 71 |
| | 69 | 83 | 6 | 52 | 47 | 0 | 0 | 8 | 65 | 0 |
| | 71 | 39 | 5 | 63 | 0 | 0 | 31 | 0 | 35 | 0 |
| | 45 | 0 | 12 | 78 | 6 | 11 | 5 | 0 | 18 | 30 |
| | 0 | 59 | 36 | 83 | 0 | 0 | 0 | 0 | 27 | 0 |
| | 4 | 29 | 5 | 71 | 9 | 39 | 31 | 0 | 55 | 35 |
| | 62 | 151 | 20 | 32 | 26 | 21 | 43 | 0 | 16 | 8 |
| | 49 | 109 | 14 | 43 | 42 | 0 | 57 | 12 | 24 | 0 |
| | 95 | 66 | 22 | 89 | 59 | 0 | 73 | 9 | 18 | 0 |
| | 0 | 11 | 0 | 0 | 72 | 24 | 0 | 0 | 18 | 25 |
| | 67 | 34 | 0 | 38 | 159 | 0 | 22 | 9 | 24 | 12 |
| | 40 | 12 | 12 | 219 | 37 | 0 | 4 | 0 | 0 | 0 |
| | 64 | 58 | 59 | 0 | 32 | 3 | 57 | 22 | 22 | 6 |
| | 55 | 16 | 13 | 27 | 111 | 0 | 18 | 1 | 13 | 32 |
| | 51 | 36 | 0 | 83 | 3 | 2 | 23 | 0 | 0 | 0 |
| | 78 | 113 | 11 | 57 | 71 | 34 | 41 | 35 | 29 | 5 |
| | 11 | 123 | 19 | 30 | 18 | 0 | 0 | 0 | 0 | 68 |
| | 40 | 109 | 28 | 81 | 41 | 6 | 46 | 5 | 12 | 0 |
| | 66 | 35 | 35 | 25 | 30 | 0 | 42 | 0 | 19 | 29 |
| | 0 | 0 | 5 | 44 | 21 | 3 | 8 | 0 | 39 | 21 |
| | 98 | 38 | 0 | 19 | 0 | 0 | 1 | 1 | 0 | 0 |
| | 55 | 41 | 0 | 61 | 0 | 12 | 73 | 21 | 9 | 1 |
| | 118 | 0 | 3 | 9 | 47 | 3 | 69 | 9 | 34 | 6 |
| | 12 | 59 | 20 | 54 | 18 | 0 | 4 | 0 | 0 | 0 |
| | 31 | 69 | 29 | 11 | 0 | 0 | 0 | 46 | 29 | 32 |
| | 47 | 0 | 53 | 0 | 19 | 0 | 19 | 0 | 0 | 7 |
| | 83 | 76 | 16 | 26 | 6 | 0 | 2 | 0 | 5 | 0 |
| | 27 | 121 | 0 | 24 | 63 | 5 | 0 | 0 | 33 | 1 |
| | 36 | 42 | 4 | 39 | 0 | 0 | 12 | 14 | 12 | 0 |
| | 0 | 56 | 0 | 43 | 41 | 0 | 0 | 28 | 12 | 33 |
| | 7 | 0 | 0 | 49 | 24 | 0 | 18 | 0 | 31 | 0 |
| | 79 | 86 | 6 | 133 | 56 | 0 | 40 | 4 | 18 | 25 |
| | 81 | 43 | 15 | 39 | 51 | 0 | 16 | 2 | 0 | 0 |
| | 0 | 0 | 22 | 53 | 28 | 4 | 19 | 3 | 9 | 0 |
| | 57 | 60 | 46 | 86 | 178 | 0 | 41 | 21 | 14 | 0 |
| | 52 | 37 | 62 | 48 | 45 | 0 | 21 | 26 | 12 | 30 |
| | 81 | 28 | 29 | 0 | 47 | 19 | 57 | 0 | 0 | 1 |
| | 0 | 5 | 0 | 18 | 133 | 0 | 5 | 1 | 39 | 0 |
| | 16 | 21 | 2 | 51 | 96 | 3 | 26 | 0 | 0 | 44 |

| *Oocyst scores* | | | | | | | | | | |
|---|---|---|---|---|---|---|---|---|---|---|
| **Uninfected** | 9 | 8 | 13 | 6 | 6 | 31 | 7 | 24 | 10 | 22 |
| **Infected** | 41 | 42 | 37 | 44 | 44 | 19 | 43 | 26 | 40 | 28 |
| **Mean Intensity** | 49.04 | 48.01 | 15.08 | 45.76 | 40.81 | 4.68 | 25.46 | 6.76 | 16.98 | 13.02 |
| **Std. Deviation** | 41.31 | 39.19 | 16.16 | 40.51 | 41.01 | 9.34 | 24.34 | 11.15 | 16.01 | 18.55 |
| **Std. Error** | 5.84 | 5.54 | 2.29 | 5.73 | 5.80 | 1.32 | 3.44 | 1.58 | 2.27 | 2.62 |
| **Prevalence** | 82% | 84% | 74% | 88% | 88% | 38% | 86% | 52% | 80% | 56% |

| *Transmission blocking efficacy* | | | | | | | | | | |
|---|---|---|---|---|---|---|---|---|---|---|
| **Intensity (%)** | 100% | | 31.1% | 94.3% | 84.1% | 9.6% | 52.5% | 13.9% | 35.0% | 26.8% |
| **Prevalence (%)** | 83% | | 74% | 88% | 88% | 38% | 86% | 52% | 80% | 56% |

#### Table S2: Transmission blocking efficacy of Primaquine and Tafenoquine in DFA (continued)

| *Oocyst scores* | 0h | | 24h | | 48h | | 72h | | 96h | |
|---|---|---|---|---|---|---|---|---|---|---|
| | Untreated | | Primaquine | Tafenoquine | Primaquine | Tafenoquine | Primaquine | Tafenoquine | Primaquine | Tafenoquine |
| Experiment 1 | 39 | 61 | 0 | 56 | 33 | 0 | 41 | 2 | 0 | 0 |
| Experiment 2 | 103 | 0 | 0 | 67 | 0 | 0 | 52 | 0 | 0 | 0 |
| Experiment 3 | 87 | 87 | 0 | 95 | 0 | 13 | 0 | 0 | 0 | 0 |
| | 57 | 52 | 0 | 21 | 0 | 23 | 0 | 0 | 2 | 0 |
| | 5 | 78 | 0 | 6 | 0 | 4 | 6 | 0 | 0 | 0 |
| | 78 | 19 | 0 | 0 | 16 | 0 | 44 | 0 | 0 | 0 |
| | 0 | 98 | 0 | 86 | 46 | 0 | 0 | 0 | 0 | 0 |
| | 0 | 92 | 0 | 36 | 1 | 2 | 0 | 7 | 0 | 0 |
| | 48 | 111 | 0 | 14 | 0 | 0 | 3 | 4 | 0 | 0 |
| | 29 | 4 | 0 | 6 | 0 | 0 | 0 | 0 | 0 | 0 |
| | 66 | 60 | 0 | 1 | 0 | 4 | 14 | 0 | 0 | 0 |
| | 0 | 82 | 0 | 82 | 1 | 0 | 23 | 0 | 1 | 0 |
| | 46 | 62 | 0 | 56 | 0 | 0 | 22 | 0 | 4 | 0 |
| | 57 | 51 | 0 | 12 | 0 | 3 | 17 | 0 | 0 | 0 |
| | 106 | 35 | 0 | 90 | 0 | 6 | 0 | 0 | 0 | 0 |
| | 46 | 50 | 0 | 10 | 12 | 6 | 0 | 0 | 0 | 0 |
| | 19 | 106 | 0 | 77 | 0 | 0 | 0 | 0 | 0 | 15 |
| | 0 | 30 | 0 | 31 | 32 | 0 | 0 | 0 | 0 | 0 |
| | 11 | 69 | 0 | 59 | 0 | 0 | 13 | 0 | 0 | 0 |
| | 182 | 4 | 0 | 31 | 2 | 5 | 37 | 0 | 0 | 0 |
| | 42 | 62 | 0 | 83 | 0 | 0 | 0 | 0 | 0 | 0 |
| | 47 | 114 | 0 | 71 | 13 | 0 | 4 | 0 | 25 | 0 |
| | 29 | 94 | 0 | 84 | 0 | 0 | 0 | 0 | 0 | 7 |
| | 16 | 117 | 0 | 54 | 7 | 0 | 0 | 0 | 8 | 0 |
| | 20 | 63 | 0 | 0 | 0 | 0 | 49 | 0 | 0 | 0 |
| | 41 | 64 | 0 | 0 | 0 | 0 | 0 | 0 | 16 | 0 |
| | 42 | 134 | 0 | 26 | 22 | 4 | 7 | 0 | 0 | 0 |
| | 77 | 65 | 0 | 0 | 15 | 0 | 24 | 0 | 0 | 0 |
| | 10 | 47 | 0 | 12 | 0 | 4 | 28 | 0 | 7 | 0 |
| | 35 | 0 | 0 | 4 | 0 | 0 | 0 | 0 | 0 | 0 |
| | 2 | 69 | 0 | 4 | 0 | 0 | 0 | 0 | 0 | 78 |
| | 2 | 51 | 0 | 83 | 6 | 0 | 21 | 4 | 25 | 0 |
| | 14 | 53 | 0 | 9 | 0 | 0 | 11 | 0 | 9 | 0 |
| | 31 | 0 | 0 | 54 | 0 | 0 | 23 | 0 | 0 | 0 |
| | 0 | 37 | 0 | 40 | 0 | 0 | 0 | 0 | 0 | 0 |
| | 61 | 25 | 0 | 48 | 3 | 0 | 26 | 0 | 0 | 0 |
| | 42 | 27 | 0 | 0 | 0 | 0 | 0 | 0 | 0 | 0 |
| | 31 | 23 | 0 | 22 | 0 | 0 | 0 | 0 | 0 | 0 |
| | 22 | 86 | 0 | 8 | 0 | 6 | 0 | 0 | 0 | 0 |
| | 5 | 51 | 0 | 33 | 0 | 0 | 0 | 0 | 0 | 0 |
| | 22 | 65 | 0 | 47 | 0 | 0 | 0 | 0 | 0 | 0 |
| | 63 | 43 | 0 | 9 | 0 | 0 | 0 | 0 | 0 | 3 |
| | 26 | 0 | 0 | 22 | 0 | 0 | 0 | 0 | 0 | 0 |
| | 12 | | 0 | 2 | 0 | 6 | 0 | 7 | 22 | 0 |
| | 39 | | 0 | 5 | 0 | 3 | 0 | 0 | 0 | 0 |
| | 0 | | 0 | 0 | 0 | 0 | 50 | 0 | 0 | 31 |
| | 4 | | 0 | 27 | 0 | 20 | 0 | 0 | 0 | 2 |
| | 26 | | 0 | 47 | 0 | 0 | 5 | 0 | 0 | 0 |
| | 43 | | 0 | 48 | 4 | 0 | 0 | 0 | 0 | 0 |
| | 11 | | 0 | 0 | 0 | 0 | 0 | 0 | 0 | 0 |

*Oocyst scores*

| | | | | | | | | | | |
|---|---|---|---|---|---|---|---|---|---|---|
| **Uninfected** | 6 | 4 | 50 | 7 | 35 | 35 | 28 | 45 | 40 | 44 |
| **Infected** | 44 | 39 | 0 | 43 | 15 | 15 | 22 | 5 | 10 | 6 |
| **Mean Intensity** | 56.77 | 56.77 | 0.00 | 33.56 | 4.35 | 2.18 | 10.40 | 0.48 | 2.72 | 2.72 |
| **Std. Deviation** | 34.48 | 34.85 | 0.00 | 30.47 | 9.81 | 4.77 | 15.72 | 1.58 | 6.26 | 11.92 |
| **Std. Error** | 4.88 | 5.31 | 0.00 | 4.31 | 1.39 | 0.67 | 2.22 | 0.22 | 0.89 | 1.69 |
| **Prevalence** | 88% | 91% | 0% | 86% | 30% | 30% | 44% | 10% | 20% | 12% |

*Transmission blocking efficacy*

| | | | | | | | | | |
|---|---|---|---|---|---|---|---|---|---|
| **Intensity (%)** | 100% | 0.0% | 59.1% | 7.7% | 3.8% | 18.3% | 0.8% | 4.8% | 4.8% |
| **Prevalence (%)** | 89% | 0% | 86% | 30% | 30% | 44% | 10% | 20% | 12% |

**Table S3: Pharmacokinetic profiles of Primaquine and Tafenoquine in the M2M model.** ND: not detected (below the quantification level).

***Exposure levels***

| **Time post-treatement (h)** | **Primaquine 1x3.35 mg/kg** | | | | | | |
|---|---|---|---|---|---|---|---|
| | M1.1 (ng/ml) | M1.2 (ng/ml) | M2.1 (ng/ml) | M2.2 (ng/ml) | **Mean (ng/ml)** | **SD (ng/ml)** | **CV (%)** |
| 24 | ND | ND | ND | ND | | | |
| 48 | ND | ND | ND | ND | | | |
| 72 | ND | ND | ND | ND | | | |
| 96 | ND | ND | ND | ND | | | |

***Exposure levels***

| **Time post-treatement (h)** | **Tafenoquine 3.41 mg/kg** | | | | | | |
|---|---|---|---|---|---|---|---|
| | M1.1 (ng/ml) | M1.2 (ng/ml) | M2.1 (ng/ml) | M2.2 (ng/ml) | **Mean (ng/ml)** | **SD (ng/ml)** | **CV (%)** |
| 24 | 43.9* | 268.5 | 202.5 | 308.4 | **259.8** | **53.5** | **20.6** |
| 48 | 213.9 | 274 | 354.1 | 185.4 | **256.85** | **74.6** | **29.0** |
| 72 | 330.4 | 160.1 | 329.3 | 333.7 | **288.375** | **85.5** | **29.7** |
| 96 | 425.3 | 256.7 | 409.7 | 292 | **345.925** | **84.1** | **24.3** |

* value omitted from mean calculations

**Table S4: Effect Primaquine and Tafenoquine against sexual (A) and asexual (B) blood stages in the M2M model**

(A)

***Asexual Blod stages***

| Time post-treatement (h) | Primaquine M1 | Primaquine M2 | Primaquine M3 | Primaquine M4 | Primaquine M5 | Primaquine **Mean** | Tafenoquine M1 | Tafenoquine M2 | Tafenoquine M3 | Tafenoquine M4 | Tafenoquine M5 | Tafenoquine **Mean** |
|---|---|---|---|---|---|---|---|---|---|---|---|---|
| 0h | 6.86% | 9.37% | 4.70% | 8.89% | 6.67% | 7.36% | 13.58% | 5.98% | 6.61% | 7.29% | 3.67% | 7.36% |
| 24h | 2.40% | 7.50% | 4.27% | 3.33% | | 4.38% | 6.19% | 6.39% | 8.19% | 7.14% | | 6.98% |
| 48h | | 10.98% | 4.24% | 7.83% | | 7.68% | | 3.00% | 6.21% | 3.77% | | 4.33% |
| 72h | | | 12.63% | 10.19% | | 11.41% | | | 4.62% | 0.83% | | 2.72% |
| 96h | | | | 19.18% | | 19.18% | | | | 0.00% | | 0.00% |

| Time post-treatement (h) | Primaquine M1 | Primaquine M2 | Primaquine M3 | Primaquine M4 | Primaquine M5 | Primaquine **Mean** | Tafenoquine M1 | Tafenoquine M2 | Tafenoquine M3 | Tafenoquine M4 | Tafenoquine M5 | Tafenoquine **Mean** |
|---|---|---|---|---|---|---|---|---|---|---|---|---|
| 0h | 8.00% | 6.58% | 6.63% | 6.81% | 12.41% | 9.00% | 9.03% | 9.18% | 10.86% | 9.86% | 10.61% | 9.00% |
| 24h | 2.69% | 1.88% | 3.04% | 3.00% | | 2.65% | 8.26% | 5.78% | 4.81% | 7.44% | | 6.57% |
| 48h | | 10.73% | 5.16% | 5.96% | | 7.28% | | 6.51% | 9.27% | 5.44% | | 7.07% |
| 72h | | | 9.34% | 10.64% | | 9.99% | | | 8.29% | 13.33% | | 10.81% |
| 96h | | | | 6.86% | | 6.86% | | | | 20.00% | | 20.00% |

| Time post-treatement (h) | Primaquine M1 | Primaquine M2 | Primaquine M3 | Primaquine M4 | Primaquine M5 | Primaquine **Mean** | Tafenoquine M1 | Tafenoquine M2 | Tafenoquine M3 | Tafenoquine M4 | Tafenoquine M5 | Tafenoquine **Mean** |
|---|---|---|---|---|---|---|---|---|---|---|---|---|
| 0h | 11.27% | 11.20% | 11.88% | 12.32% | 8.25% | 9.67% | 9.44% | 6.51% | 9.34% | 10.37% | 6.07% | 9.67% |
| 24h | 9.08% | 10.30% | 5.53% | 8.47% | | 8.35% | 8.18% | 4.13% | 17.40% | 7.31% | | 9.25% |
| 48h | | 13.33% | 8.54% | 12.73% | | 11.53% | | 2.59% | 7.80% | 13.73% | | 8.04% |
| 72h | | | 19.47% | 16.04% | | 17.76% | | | 5.00% | 29.78% | | 17.39% |
| 96h | | | | 28.78% | | 28.78% | | | | 27.25% | | 27.25% |

Experiment 1

Experiment 2

Experiment 3

**Table S4: Effect Primaquine and Tafenoquine against sexual (A) and asexual (B) blood stages in the M2M model (continued)**

(B)

***Gametocytes***

| | Primaquine | | | | | | Tafenoquine | | | | | |
|---|---|---|---|---|---|---|---|---|---|---|---|---|
| Time post-treatement (h) | M1 | M2 | M3 | M4 | M5 | **Mean** | M1 | M2 | M3 | M4 | M5 | **Mean** |
| 0h | 0.74% | 0.55% | 2.39% | 0.81% | 0.88% | 1.22% | 1.36% | 0.38% | 1.43% | 1.45% | 2.22% | 1.22% |
| 24h | 0.00% | 0.00% | 0.53% | 0.53% | | 0.27% | 0.00% | 0.69% | 0.64% | 0.44% | | 0.44% |
| 48h | | 0.12% | 0.17% | 0.00% | | 0.10% | | 0.67% | 0.52% | 0.43% | | 0.54% |
| 72h | | | 0.00% | 0.19% | | 0.09% | | | 0.00% | 0.00% | | 0.00% |
| 96h | | | | 0.41% | | 0.41% | | | | 0.00% | | 0.00% |

| | Primaquine | | | | | | Tafenoquine | | | | | |
|---|---|---|---|---|---|---|---|---|---|---|---|---|
| Time post-treatement (h) | M1 | M2 | M3 | M4 | M5 | **Mean** | M1 | M2 | M3 | M4 | M5 | **Mean** |
| 0h | 3.50% | 2.05% | 1.88% | 1.59% | 1.72% | 2.15% | 1.45% | 2.04% | 4.00% | 0.82% | 2.45% | 2.15% |
| 24h | 0.96% | 1.09% | 0.89% | 2.33% | | 1.32% | 1.74% | 1.33% | 0.76% | 1.54% | | 1.34% |
| 48h | | 2.68% | 0.94% | 0.70% | | 1.44% | | 0.63% | 1.45% | 0.88% | | 0.99% |
| 72h | | | 0.49% | 0.85% | | 0.67% | | | 1.14% | 1.04% | | 1.09% |
| 96h | | | | 0.00% | | 0.00% | | | | 0.40% | | 0.40% |

| | Primaquine | | | | | | Tafenoquine | | | | | |
|---|---|---|---|---|---|---|---|---|---|---|---|---|
| Time post-treatement (h) | M1 | M2 | M3 | M4 | M5 | **Mean** | M1 | M2 | M3 | M4 | M5 | **Mean** |
| 0h | 2.58% | 3.20% | 1.88% | 1.79% | 1.23% | 1.92% | 1.67% | 0.95% | 1.18% | 3.33% | 1.43% | 1.92% |
| 24h | 0.92% | 1.34% | 1.28% | 1.02% | | 1.14% | 0.91% | 0.32% | 2.00% | 1.54% | | 1.19% |
| 48h | | 1.28% | 1.95% | 1.59% | | 1.61% | | 0.17% | 0.51% | 1.02% | | 0.57% |
| 72h | | | 1.40% | 0.83% | | 1.12% | | | 0.50% | 2.39% | | 1.45% |
| 96h | | | | 1.63% | | 1.63% | | | | 1.37% | | 1.37% |

Experiment 1
Experiment 2
Experiment 3

**Table S5: Transmission blocking efficacy of Primaquine and Tafenoquine + Sulphadiazine in DFA**

PQ: Primaquine, TF: Tafenoquine, SD: Sulphadiazine

| *Oocytes scores* | 0h | 24h | | | 48h | | | 72h | | | 96h | | |
|---|---|---|---|---|---|---|---|---|---|---|---|---|---|
| | Untreated | PQ + SD | TF + SD | SD | PQ + SD | TF + SD | SD | PQ + SD | TF + SD | SD | PQ + SD | TF + SD | SD |
| Experiment 1 | 54 | 1 | 0 | 72 | 1 | 0 | 86 | 0 | 0 | 0 | 0 | 0 | 0 |
| Experiment 2 | 150 | 1 | 0 | 73 | 5 | 0 | 58 | 0 | 0 | 0 | 0 | 0 | 0 |
| Experiment 3 | 32 | 1 | 0 | 0 | 5 | 0 | 64 | 0 | 0 | 0 | 0 | 0 | 0 |
| | 102 | 1 | 0 | 0 | 1 | 0 | 76 | 0 | 0 | 0 | 0 | 0 | 0 |
| | 80 | 0 | 0 | 80 | 15 | 0 | 15 | 0 | 0 | 0 | 0 | 0 | 0 |
| | 145 | 0 | 0 | 85 | 2 | 0 | 17 | 0 | 0 | 0 | 0 | 0 | 0 |
| | 64 | 0 | 0 | 115 | 5 | 0 | 99 | 0 | 0 | 0 | 0 | 0 | 0 |
| | 39 | 0 | 0 | 140 | 7 | 0 | 165 | 0 | 0 | 0 | 0 | 0 | 0 |
| | 38 | 0 | 0 | 0 | 0 | 0 | 30 | 0 | 0 | 0 | 0 | 0 | 0 |
| | 92 | 0 | 0 | 57 | 0 | 0 | 24 | 0 | 0 | 0 | 0 | 0 | 0 |
| | 80 | 0 | 0 | 60 | 0 | 0 | 75 | 0 | 0 | 0 | 0 | 0 | 0 |
| | 4 | 0 | 0 | 153 | 0 | 0 | 38 | 0 | 0 | 0 | 0 | 0 | 0 |
| | 164 | 0 | 0 | 136 | 0 | 0 | 100 | 0 | 0 | 0 | 0 | 0 | 0 |
| | 89 | 0 | 0 | 121 | 0 | 0 | 19 | 0 | 0 | 0 | 0 | 0 | 0 |
| | 20 | 0 | 0 | 8 | 0 | 0 | 63 | 0 | 0 | 0 | 0 | 0 | 0 |
| | 92 | 0 | 0 | 40 | 0 | 0 | 63 | 0 | 0 | 0 | 0 | 0 | 0 |
| | 131 | 0 | 0 | 0 | 0 | 0 | 52 | 0 | 0 | 0 | 0 | 0 | 0 |
| | 132 | 0 | 0 | 119 | 0 | 0 | 66 | 0 | 0 | 0 | 0 | 0 | 0 |
| | 137 | 0 | 0 | 7 | 0 | 0 | 100 | 0 | 0 | 0 | 0 | 0 | 0 |
| | 0 | 0 | 0 | 39 | 0 | 0 | 39 | 0 | 0 | 0 | 0 | 0 | 0 |
| | 36 | 0 | 0 | 60 | 0 | 0 | 109 | 0 | 0 | 0 | 0 | 0 | 0 |
| | 52 | 0 | 0 | 0 | 0 | 0 | 3 | 0 | 0 | 0 | 0 | 0 | 0 |
| | 0 | 0 | 0 | 23 | 0 | 0 | 85 | 0 | 0 | 0 | 0 | 0 | 0 |
| | 62 | 0 | 0 | 111 | 0 | 0 | 79 | 0 | 0 | 0 | 0 | 0 | 0 |
| | 118 | 0 | 0 | 67 | 0 | 0 | 9 | 0 | 0 | 0 | 0 | 0 | 0 |
| | 31 | 0 | 0 | 95 | 0 | 0 | 0 | 0 | 0 | 0 | 0 | 0 | 0 |
| | 103 | 0 | 0 | 85 | 0 | 0 | 69 | 0 | 0 | 0 | 0 | 0 | 0 |
| | 50 | 0 | 0 | 69 | 0 | 0 | 55 | 0 | 0 | 0 | 0 | 0 | 0 |
| | 43 | 0 | 0 | 103 | 0 | 0 | 38 | 0 | 0 | 0 | 0 | 0 | 0 |
| | 31 | 0 | 0 | 33 | 0 | 0 | 93 | 0 | 0 | 0 | 0 | 0 | 0 |
| | 174 | 0 | 0 | 128 | 0 | 0 | 59 | 0 | 0 | 0 | 0 | 0 | 0 |
| | 71 | 0 | 0 | 140 | 0 | 0 | 104 | 0 | 0 | 0 | 0 | 0 | 0 |
| | 63 | 0 | 0 | 156 | 0 | 0 | 76 | 0 | 0 | 0 | 0 | 0 | 0 |
| | 79 | 0 | 0 | 41 | 0 | 0 | 34 | 0 | 0 | 0 | 0 | 0 | 0 |
| | 76 | 0 | 0 | 3 | 0 | 0 | 77 | 0 | 0 | 0 | 0 | 0 | 0 |
| | 112 | 0 | 0 | 172 | 0 | 0 | 62 | 0 | 0 | 0 | 0 | 0 | 0 |
| | 130 | 0 | 0 | 88 | 0 | 0 | 72 | 0 | 0 | 0 | 0 | 0 | 0 |
| | 9 | 0 | 0 | 78 | 0 | 0 | 44 | 0 | 0 | 0 | 0 | 0 | 0 |
| | 8 | 0 | 0 | 69 | 0 | 0 | 73 | 0 | 0 | 0 | 0 | 0 | 0 |
| | 12 | 0 | 0 | 140 | 0 | 0 | 0 | 0 | 0 | 0 | 0 | 0 | 0 |
| | 129 | 0 | 0 | 81 | 0 | 0 | 95 | 0 | 0 | 0 | 0 | 0 | 0 |
| | 153 | 0 | 0 | 151 | 0 | 0 | 44 | 0 | 0 | 0 | 0 | 0 | 0 |
| | 111 | 0 | 0 | 107 | 0 | 0 | 69 | 0 | 0 | 0 | 0 | 0 | 0 |
| | 32 | 0 | 0 | 93 | 0 | 0 | 0 | 0 | 0 | 0 | 0 | 0 | 0 |
| | 62 | 0 | 0 | 64 | 0 | 0 | 0 | 0 | 0 | 0 | 0 | 0 | 0 |
| | 163 | 0 | 0 | 113 | 0 | 0 | 50 | 0 | 0 | 0 | 0 | 0 | 0 |
| | 89 | 0 | 0 | 103 | 0 | 0 | 42 | 0 | 0 | 0 | 0 | 0 | 0 |
| | 0 | 0 | | 14 | 0 | 0 | 38 | 0 | 0 | 0 | 0 | 0 | 0 |
| | 34 | 0 | | 14 | 0 | 0 | 33 | 0 | 0 | 0 | 0 | 0 | 0 |
| | 62 | 0 | | 6 | 0 | 0 | 104 | 0 | 0 | 0 | 0 | 0 | 0 |

*Oocytes scores*

| | | | | | | | | | | | | | |
|---|---|---|---|---|---|---|---|---|---|---|---|---|---|
| **Uninfected** | 3 | 46 | 47 | 5 | 42 | 50 | 4 | 50 | 50 | 50 | 50 | 50 | 50 |
| **Infected** | 47 | 4 | 0 | 45 | 8 | 0 | 46 | 0 | 0 | 0 | 0 | 0 | 0 |
| **Mean Intensity** | 74.80 | 0.08 | 0.00 | 74.24 | 0.82 | 0.00 | 57.30 | 0.00 | 0.00 | 0.00 | 0.00 | 0.00 | 0.00 |
| **Std. Deviation** | 49.47 | 0.27 | 0.00 | 50.10 | 2.56 | 0.00 | 34.86 | 0.00 | 0.00 | 0.00 | 0.00 | 0.00 | 0.00 |
| **Std. Error** | 7.00 | 0.04 | 0.00 | 7.08 | 0.36 | 0.00 | 4.93 | 0.00 | 0.00 | 0.00 | 0.00 | 0.00 | 0.00 |
| **Prevalence** | 94% | 8% | 0% | 90% | 16% | 0% | 92% | 0% | 0% | 0% | 0% | 0.00 | 0.00 |

*Transmission blocking efficacy*

| | | | | | | | | | | | | | |
|---|---|---|---|---|---|---|---|---|---|---|---|---|---|
| **Intensity (%)** | 100% | 0.1% | 0.0% | 99.3% | 1.1% | 0.0% | 76.6% | 0.0% | 0.0% | 0.0% | 0.0% | 0.0% | 0.0% |
| **Prevalence (%)** | 94% | 8% | 0% | 90% | 16% | 0% | 92% | 0% | 0% | 0% | 0% | 0% | 0% |

#### Table S5: Transmission blocking efficacy of Primaquine and Tafenoquine + Sulphadiazine in DFA (continued)

PQ: Primaquine, TF: Tafenoquine, SD: Sulphadiazine

| *Oocytes scores* | 0h | 24h | | | 48h | | | 72h | | | 96h | | |
|---|---|---|---|---|---|---|---|---|---|---|---|---|---|
| | Untreated | PQ + SD | TF + SD | SD | PQ + SD | TF + SD | SD | PQ + SD | TF + SD | SD | PQ + SD | TF + SD | SD |
| Experiment 1 | 64 | 22 | 0 | 5 | 2 | 0 | 19 | 0 | 0 | 1 | 1 | 0 | 0 |
| Experiment 2 | 74 | 197 | 0 | 91 | 2 | 0 | 32 | 0 | 0 | 2 | 1 | 0 | 0 |
| Experiment 3 | 165 | 305 | 5 | 61 | 1 | 0 | 16 | 0 | 0 | 3 | 0 | 0 | 0 |
| | 0 | 0 | 298 | 115 | 0 | 0 | 33 | 0 | 0 | 1 | 0 | 0 | 0 |
| | 25 | 99 | 49 | 48 | 0 | 0 | 118 | 0 | 0 | 5 | 0 | 0 | 0 |
| | 58 | 176 | 240 | 73 | 1 | 0 | 67 | 0 | 0 | 2 | 0 | 0 | 0 |
| | 8 | 20 | 32 | 236 | 1 | 0 | 4 | 0 | 0 | 0 | 0 | 0 | 0 |
| | 27 | 167 | 64 | 231 | 1 | 0 | 6 | 0 | 0 | 0 | 0 | 0 | 0 |
| | 6 | 95 | 44 | 314 | 0 | 0 | 48 | 0 | 0 | 0 | 0 | 0 | 0 |
| | 11 | 13 | 4 | 226 | 0 | 0 | 1 | 0 | 0 | 0 | 0 | 0 | 0 |
| | 15 | 172 | 86 | 251 | 0 | 0 | 196 | 0 | 0 | 0 | 0 | 0 | 0 |
| | 32 | 32 | 5 | 83 | 0 | 0 | 9 | 0 | 0 | 0 | 0 | 0 | 0 |
| | 142 | 16 | 93 | 212 | 0 | 0 | 24 | 0 | 0 | 0 | 0 | 0 | 0 |
| | 250 | 11 | 4 | 140 | 0 | 0 | 11 | 0 | 0 | 0 | 0 | 0 | 0 |
| | 1 | 0 | 253 | 212 | 0 | 0 | 57 | 0 | 0 | 0 | 0 | 0 | 0 |
| | 129 | 0 | 35 | 84 | 0 | 0 | 4 | 0 | 0 | 0 | 0 | 0 | 0 |
| | 134 | 0 | 28 | 11 | 0 | 0 | 0 | 0 | 0 | 0 | 0 | 0 | 0 |
| | 14 | 9 | 339 | 9 | 0 | 0 | 49 | 0 | 0 | 0 | 0 | 0 | 0 |
| | 108 | 22 | 82 | 59 | 0 | 0 | 7 | 0 | 0 | 0 | 0 | 0 | 0 |
| | 53 | 115 | 0 | 48 | 0 | 0 | 228 | 0 | 0 | 0 | 0 | 0 | 0 |
| | 5 | 119 | 0 | 5 | 0 | 0 | 50 | 0 | 0 | 0 | 0 | 0 | 0 |
| | 8 | 22 | 185 | 0 | 0 | 0 | 11 | 0 | 0 | 0 | 0 | 0 | 0 |
| | 0 | 14 | 20 | 7 | 0 | 0 | 130 | 0 | 0 | 0 | 0 | 0 | 0 |
| | 38 | 84 | 3 | 85 | 0 | 0 | 4 | 0 | 0 | 0 | 0 | 0 | 0 |
| | 154 | 16 | 90 | 21 | 0 | 0 | 5 | 0 | 0 | 0 | 0 | 0 | 0 |
| | 21 | 97 | 29 | 47 | 0 | 0 | 4 | 0 | 0 | 0 | 0 | 0 | 0 |
| | 0 | 20 | 148 | 0 | 0 | 0 | 38 | 0 | 0 | 0 | 0 | 0 | 0 |
| | 159 | 0 | 160 | 12 | 0 | 0 | 231 | 0 | 0 | 0 | 0 | 0 | 0 |
| | 131 | 14 | 47 | 64 | 0 | 0 | 32 | 0 | 0 | 0 | 0 | 0 | 0 |
| | 70 | 69 | 0 | 35 | 0 | 0 | 5 | 0 | 0 | 0 | 0 | 0 | 0 |
| | 157 | 120 | 176 | 71 | 0 | 0 | 0 | 0 | 0 | 0 | 0 | 0 | 0 |
| | 272 | 0 | 14 | 17 | 0 | 0 | 7 | 0 | 0 | 0 | 0 | 0 | 0 |
| | 72 | 16 | 22 | 56 | 0 | 0 | 19 | 0 | 0 | 0 | 0 | 0 | 0 |
| | 9 | 24 | 169 | 0 | 0 | 0 | 0 | 0 | 0 | 0 | 0 | 0 | 0 |
| | 149 | 0 | 168 | 0 | 0 | 0 | 21 | 0 | 0 | 0 | 0 | 0 | 0 |
| | 100 | 19 | 6 | 40 | 0 | 0 | 21 | 0 | 0 | 0 | 0 | 0 | 0 |
| | 4 | 35 | 186 | 56 | 1 | 0 | 6 | 0 | 0 | 0 | 0 | 0 | 0 |
| | 6 | 18 | 88 | 98 | 0 | 0 | 3 | 0 | 0 | 0 | 0 | 0 | 0 |
| | 1 | 21 | 153 | 63 | 0 | 0 | 4 | 0 | 0 | 0 | 0 | 0 | 0 |
| | 233 | 144 | 8 | 49 | 0 | 0 | 3 | 0 | 0 | 0 | 0 | 0 | 0 |
| | 3 | 55 | 40 | 0 | 0 | 0 | 3 | 0 | 0 | 0 | 0 | 0 | 0 |
| | 85 | 8 | 41 | 5 | 1 | 0 | 9 | 0 | 0 | 0 | 0 | 0 | 0 |
| | 411 | 8 | 2 | 91 | 0 | 0 | 8 | 0 | 0 | 0 | 0 | 0 | 0 |
| | 4 | 4 | 27 | 40 | 0 | 0 | 8 | 0 | 0 | 0 | 0 | 0 | 0 |
| | 57 | 7 | 3 | 69 | 0 | 0 | 6 | 0 | 0 | 0 | 0 | 0 | 0 |
| | 154 | 185 | 42 | 48 | 0 | 0 | 23 | 0 | 0 | 0 | 0 | 0 | 0 |
| | 97 | 44 | 0 | 103 | 0 | 0 | 8 | 0 | 0 | 0 | 0 | 0 | 0 |
| | 15 | 28 | 0 | 97 | 0 | 0 | 27 | 0 | 0 | 0 | 0 | 0 | 0 |
| | 33 | 0 | 1 | | 0 | 0 | 92 | 0 | 0 | 0 | 0 | 0 | 0 |
| | 131 | 140 | 48 | | 0 | 0 | 148 | 0 | 0 | 0 | 0 | 0 | 0 |

*Oocytes scores*

| | | | | | | | | | | | | | |
|---|---|---|---|---|---|---|---|---|---|---|---|---|---|
| Uninfected | 3 | 8 | 7 | 5 | 42 | 50 | 3 | 50 | 50 | 44 | 48 | 50 | 50 |
| Infected | 47 | 42 | 43 | 43 | 8 | 0 | 47 | 0 | 0 | 6 | 2 | 0 | 0 |
| Mean Intensity | 77.90 | 56.04 | 70.74 | 76.83 | 0.20 | 0.00 | 37.10 | 0.00 | 0.00 | 0.28 | 0.04 | 0.00 | 0.00 |
| Std. Deviation | 86.42 | 69.01 | 86.32 | 77.49 | 0.49 | 0.00 | 57.20 | 0.00 | 0.00 | 0.90 | 0.20 | 0.00 | 0.00 |
| Std. Error | 12.22 | 9.76 | 12.21 | 11.19 | 0.07 | 0.00 | 8.09 | 0.00 | 0.00 | 0.13 | 0.03 | 0.00 | 0.00 |
| Prevalence | 94% | 84% | 86% | 90% | 16% | 0% | 94% | 0% | 0% | 12% | 4% | 0% | 0% |

*Transmission blocking efficacy*

| | | | | | | | | | | | | | |
|---|---|---|---|---|---|---|---|---|---|---|---|---|---|
| Intensity (%) | 100% | 71.9% | 90.8% | 98.6% | 0.3% | 0.0% | 47.6% | 0.0% | 0.0% | 0.4% | 0.1% | 0.0% | 0.0% |
| Prevalence (%) | 94% | 84% | 86% | 90% | 16% | 0% | 94% | 0% | 0% | 12% | 4% | 0% | 0% |

**Table S5: Transmission blocking efficacy of Primaquine and Tafenoquine + Sulphadiazine in DFA (continued)**

PQ: Primaquine, TF: Tafenoquine, SD:Sulphadiazine

| ***Oocytes scores*** | **0h** | **24h** | | | **48h** | | | **72h** | | | **96h** | | |
|---|---|---|---|---|---|---|---|---|---|---|---|---|---|
| | **Untreated** | **PQ + SD** | **TF + SD** | **SD** | **PQ + SD** | **TF + SD** | **SD** | **PQ + SD** | **TF + SD** | **SD** | **PQ + SD** | **TF + SD** | **SD** |
| Experiment 1 | 18 | 0 | 0 | 49 | 9 | 1 | 36 | 0 | 0 | 1 | 0 | 0 | 0 |
| Experiment 2 | 89 | 0 | 0 | 56 | 7 | 0 | 74 | 0 | 0 | 0 | 0 | 0 | 0 |
| Experiment 3 | 50 | 0 | 0 | 65 | 7 | 0 | 49 | 0 | 0 | 0 | 0 | 0 | 0 |
| | 0 | 0 | 0 | 23 | 15 | 0 | 50 | 0 | 0 | 0 | 0 | 0 | 0 |
| | 60 | 0 | 0 | 63 | 11 | 0 | 53 | 0 | 0 | 0 | 0 | 0 | 0 |
| | 0 | 0 | 0 | 25 | 5 | 0 | 0 | 0 | 0 | 0 | 0 | 0 | 0 |
| | 71 | 0 | 0 | 51 | 6 | 0 | 21 | 0 | 0 | 0 | 0 | 0 | 0 |
| | 5 | 0 | 0 | 50 | 0 | 0 | 27 | 0 | 0 | 0 | 0 | 0 | 0 |
| | 39 | 0 | 0 | 36 | 3 | 0 | 24 | 0 | 0 | 0 | 0 | 0 | 0 |
| | 66 | 0 | 0 | 75 | 0 | 0 | 20 | 0 | 0 | 0 | 0 | 0 | 0 |
| | 0 | 0 | 0 | 7 | 11 | 0 | 12 | 0 | 0 | 0 | 0 | 0 | 0 |
| | 83 | 0 | 0 | 67 | 0 | 0 | 25 | 0 | 0 | 0 | 0 | 0 | 0 |
| | 0 | 0 | 0 | 4 | 0 | 0 | 0 | 0 | 0 | 0 | 0 | 0 | 0 |
| | 57 | 0 | 0 | 9 | 3 | 0 | 28 | 0 | 0 | 0 | 0 | 0 | 0 |
| | 42 | 0 | 0 | 80 | 10 | 0 | 85 | 0 | 0 | 0 | 0 | 0 | 0 |
| | 23 | 0 | 0 | 39 | 0 | 0 | 14 | 0 | 0 | 0 | 0 | 0 | 0 |
| | 76 | 0 | 0 | 56 | 11 | 0 | | 0 | 0 | 0 | 0 | 0 | 0 |
| | 20 | | | 62 | 0 | 0 | | 0 | 0 | 0 | 0 | 0 | 0 |
| | 0 | | | 17 | 13 | 0 | | 0 | 0 | 0 | 0 | 0 | 0 |
| | 0 | | | 0 | 12 | 0 | | 0 | 0 | 0 | 0 | 0 | 0 |
| | 0 | | | 55 | 10 | 0 | | 0 | 0 | 0 | 0 | 0 | 0 |
| | 48 | | | 24 | 14 | 0 | | 0 | 0 | 0 | 0 | 0 | 0 |
| | 0 | | | 26 | 0 | 0 | | 0 | 0 | 0 | 0 | 0 | 0 |
| | 0 | | | 7 | 3 | 0 | | 0 | 0 | 0 | 0 | 0 | 0 |
| | 24 | | | 50 | 0 | 0 | | 0 | 0 | 0 | 0 | 0 | 0 |
| | 40 | | | 50 | 0 | 0 | | 0 | 0 | 0 | 0 | 0 | 0 |
| | 0 | | | 0 | 0 | 0 | | 0 | 0 | 0 | 0 | 0 | 0 |
| | 72 | | | 3 | 2 | 0 | | 0 | 0 | 0 | 0 | 0 | 0 |
| | 27 | | | 9 | | 0 | | 0 | 0 | 0 | 0 | 0 | 0 |
| | 94 | | | 2 | | 0 | | 0 | 0 | 0 | 0 | 0 | 0 |
| | 0 | | | 89 | | 0 | | 0 | 0 | 0 | 0 | 0 | 0 |
| | 29 | | | 3 | | 0 | | 0 | 0 | 0 | 0 | 0 | 0 |
| | 0 | | | 58 | | 0 | | 0 | 0 | 0 | 0 | 0 | 0 |
| | 74 | | | 25 | | 0 | | 0 | 0 | 0 | 0 | 0 | 0 |
| | 92 | | | 63 | | 0 | | 0 | 0 | 0 | 0 | 0 | 0 |
| | 22 | | | 35 | | 0 | | 0 | 0 | 0 | 0 | 0 | 0 |
| | 86 | | | 45 | | 0 | | 0 | 0 | 0 | 0 | 0 | 0 |
| | 69 | | | 0 | | 0 | | 0 | 0 | 0 | 0 | 0 | 0 |
| | 39 | | | 54 | | 0 | | 0 | 0 | 0 | 0 | 0 | 0 |
| | 28 | | | 68 | | 0 | | 0 | 0 | 0 | 0 | 0 | 0 |
| | 0 | | | 71 | | 0 | | 0 | 0 | 0 | 0 | 0 | 0 |
| | 0 | | | 45 | | 0 | | 0 | 0 | 0 | 0 | 0 | 0 |
| | 84 | | | 53 | | 0 | | 0 | 0 | 0 | 0 | 0 | 0 |
| | 101 | | | 36 | | 0 | | 0 | 0 | 0 | 0 | 0 | 0 |
| | 80 | | | 0 | | 0 | | 0 | 0 | 0 | 0 | 0 | 0 |
| | 0 | | | 43 | | 0 | | 0 | 0 | 0 | 0 | 0 | 0 |
| | 87 | | | 62 | | 0 | | 0 | 0 | 0 | 0 | 0 | 0 |
| | 74 | | | 90 | | 0 | | 0 | 0 | 0 | 0 | 0 | 0 |
| | 0 | | | 23 | | 0 | | 0 | 0 | 0 | 0 | 0 | 0 |
| | 47 | | | 42 | | 0 | | 0 | 0 | 0 | 0 | 0 | 0 |

***Oocytes scores***

| | | | | | | | | | | | | | |
|---|---|---|---|---|---|---|---|---|---|---|---|---|---|
| **Uninfected** | 16 | 17 | 17 | 4 | 10 | 49 | 2 | 50 | 50 | 49 | 50 | 50 | 50 |
| **Infected** | 34 | 0 | 0 | 46 | 18 | 1 | 14 | 0 | 0 | 1 | 0 | 0 | 0 |
| **Mean Intensity** | 38.32 | 0.00 | 0.00 | 39.30 | 5.43 | 0.02 | 32.38 | 0.00 | 0.00 | 0.02 | 0.00 | 0.00 | 0.00 |
| **Std. Deviation** | 34.51 | 0.00 | 0.00 | 25.90 | 5.25 | 0.14 | 24.26 | 0.00 | 0.00 | 0.14 | 0.00 | 0.00 | 0.00 |
| **Std. Error** | 4.88 | 0.00 | 0.00 | 3.66 | 0.99 | 0.02 | 6.07 | 0.00 | 0.00 | 0.02 | 0.00 | 0.00 | 0.00 |
| **Prevalence** | 68% | 0% | 0% | 92% | 64% | 2% | 88% | 0% | 0% | 2% | 0% | 0% | 0% |

***Transmission blocking efficacy***

| | | | | | | | | | | | | | |
|---|---|---|---|---|---|---|---|---|---|---|---|---|---|
| **Intensity (%)** | 100% | 0.0% | 0.0% | 102.6% | 14.2% | 0.1% | 84.5% | 0.0% | 0.0% | 0.1% | 0.0% | 0.0% | 0.00 |
| **Prevalence (%)** | 68% | 0% | 0% | 92% | 64% | 2% | 88% | 0% | 0% | 2% | 0% | 0% | 0% |

#### Table S6: Effect Primaquine and Tafenoquine with Sulphadiazine against sexual (A) and asexual (B) blood stages in the M2M model

(A)

***Asexual Blood stages***

| | Primaquine + Sulphadiazine | | | | | | Tafenoquine + Sulphadiazine | | | | | Sulphadiazine | | | | |
|---|---|---|---|---|---|---|---|---|---|---|---|---|---|---|---|---|
| Time post-treatement (h) | M1 | M2 | M3 | M4 | M5 | **Mean** | M1 | M2 | M3 | M4 | **Mean** | M1 | M2 | M3 | M4 | **Mean** |
| 0h | 9.58% | 7.12% | 7.50% | 5.52% | 6.29% | 8.14% | 13.44% | 10.26% | 9.73% | 8.57% | 8.14% | 5.00% | 5.15% | 5.71% | 12.00% | 8.14% |
| 24h | 5.56% | 7.27% | 6.49% | 6.46% | | 6.45% | 11.41% | 12.39% | 7.54% | 7.19% | 9.63% | 7.65% | 5.29% | 5.79% | 15.93% | 8.67% |
| 48h | | 1.63% | 0.80% | 5.30% | | 2.58% | | 7.03% | 3.62% | 2.59% | 4.41% | | 2.27% | 1.03% | 9.36% | 4.22% |
| 72h | | | 0.00% | 0.55% | | 0.27% | | | 1.03% | 1.60% | 1.32% | | | 0.00% | 0.00% | 0.00% |
| 96h | | | | 0.00% | | 0.00% | | | | 0.00% | 0.00% | | | | 0.00% | 0.00% |

| | Primaquine + Sulphadiazine | | | | | | Tafenoquine + Sulphadiazine | | | | | Sulphadiazine | | | | |
|---|---|---|---|---|---|---|---|---|---|---|---|---|---|---|---|---|
| Time post-treatement (h) | M1 | M2 | M3 | M4 | M5 | **Mean** | M1 | M2 | M3 | M4 | **Mean** | M1 | M2 | M3 | M4 | **Mean** |
| 0h | 9.29% | 10.17% | 11.67% | 9.63% | 11.36% | 10.16% | 9.38% | 8.78% | 8.00% | 12.88% | 10.16% | 8.27% | 9.84% | 12.10% | 10.75% | 10.16% |
| 24h | 3.82% | 4.66% | 9.34% | 9.00% | | 6.70% | 5.43% | 2.11% | 9.62% | 3.95% | 5.28% | 3.20% | 2.89% | 4.24% | 2.00% | 3.08% |
| 48h | | 0.34% | 2.96% | 0.56% | | 1.29% | | 0.29% | 1.43% | 1.20% | 0.97% | | 0.55% | 0.00% | 0.45% | 0.33% |
| 72h | | | 0.00% | 0.00% | | 0.00% | | | 0.00% | 0.00% | 0.00% | | | 0.00% | 0.00% | 0.00% |
| 96h | | | | 0.00% | | 0.00% | | | | 0.00% | 0.00% | | | | 0.00% | 0.00% |

| | Primaquine + Sulphadiazine | | | | | | Tafenoquine+ Sulphadiazine | | | | | Sulphadiazine | | | | |
|---|---|---|---|---|---|---|---|---|---|---|---|---|---|---|---|---|
| Time post-treatement (h) | M1 | M2 | M3 | M4 | M5 | **Mean** | M1 | M2 | M3 | M4 | **Mean** | M1 | M2 | M3 | M4 | **Mean** |
| 0h | 9.34% | 5.34% | 6.91% | 6.76% | 9.23% | 6.54% | 5.62% | 5.42% | 5.79% | 8.27% | 6.54% | 8.57% | 3.92% | 4.60% | 5.28% | 6.54% |
| 24h | 2.81% | 1.61% | 1.00% | 3.23% | | 2.16% | 2.69% | 18.17% | 4.42% | 4.20% | 7.37% | 11.25% | 4.39% | 3.28% | 9.38% | 7.07% |
| 48h | | 0.79% | 0.61% | 0.56% | | 0.65% | | 3.44% | 1.32% | 1.59% | 2.12% | | 0.22% | 0.17% | 1.95% | 0.78% |
| 72h | | | 0.00% | 0.00% | | 0.00% | | | 0.35% | 0.00% | 0.18% | | | 0.00% | 0.71% | 0.36% |
| 96h | | | | 0.00% | | 0.00% | | | | 0.00% | 0.00% | | | | 0.00% | 0.00% |

Experiment 1
Experiment 2
Experiment 3

**Table S6: Effect Primaquine and Tafenoquine with Sulphadiazine against sexual (A) and asexual (B) blood stages in the M2M model (continued)**

(B)

***Gametocytes***

| | Primaquine + Sulphadiazine | | | | | | Tafenoquine + Sulphadiazine | | | | | Sulphadiazine | | | | |
|---|---|---|---|---|---|---|---|---|---|---|---|---|---|---|---|---|
| Time post-treatement (h) | M1 | M2 | M3 | M4 | M5 | **Mean** | M1 | M2 | M3 | M4 | **Mean** | M1 | M2 | M3 | M4 | **Mean** |
| 0h | , | 0.96% | 1.50% | 1.64% | 0.86% | 1.09% | 1.41% | 0.26% | 1.23% | 1.27% | 1.09% | 1.30% | 0.88% | 1.11% | 0.71% | 1.09% |
| 24h | 0.32% | 0.00% | 0.53% | 0.62% | | 0.36% | 0.00% | 0.30% | 0.77% | 0.18% | 0.31% | 0.20% | 0.59% | 1.05% | 0.17% | 0.50% |
| 48h | | 0.00% | 0.00% | 0.30% | | 0.10% | | 0.00% | 0.43% | 0.86% | 0.43% | | 0.23% | 0.00% | 0.00% | 0.08% |
| 72h | | | 0.00% | 0.00% | | 0.00% | | | 0.00% | 0.00% | 0.00% | | | 0.00% | 0.00% | 0.00% |
| 96h | | | | 0.00% | | 0.00% | | | | 0.00% | 0.00% | | | | 0.00% | 0.00% |

| | Primaquine + Sulphadiazine | | | | | | Tafenoquine + Sulphadiazine | | | | | Sulphadiazine | | | | |
|---|---|---|---|---|---|---|---|---|---|---|---|---|---|---|---|---|
| Time post-treatement (h) | M1 | M2 | M3 | M4 | M5 | **Mean** | M1 | M2 | M3 | M4 | **Mean** | M1 | M2 | M3 | M4 | **Mean** |
| 0h | 2.50% | 2.17% | 1.11% | 0.93% | 0.91% | 1.25% | 1.41% | 2.04% | 0.40% | 0.19% | 1.25% | 1.35% | 0.95% | 0.97% | 1.34% | 1.25% |
| 24h | 1.45% | 1.90% | 2.95% | 2.25% | | 2.14% | 1.30% | 1.40% | 1.35% | 0.92% | 1.24% | 1.33% | 2.89% | 1.21% | 1.17% | 1.65% |
| 48h | | 0.34% | 2.59% | 1.30% | | 1.41% | | 1.30% | 1.43% | 1.20% | 1.31% | | 1.45% | 1.09% | 0.60% | 1.05% |
| 72h | | | 0.00% | 0.00% | | 0.00% | | | 0.00% | 0.00% | 0.00% | | | 0.00% | 0.00% | 0.00% |
| 96h | | | | 0.00% | | 0.00% | | | | 0.00% | 0.00% | | | | 0.00% | 0.00% |

| | Primaquine + Sulphadiazine | | | | | | Tafenoquine+ Sulphadiazine | | | | | Sulphadiazine | | | | |
|---|---|---|---|---|---|---|---|---|---|---|---|---|---|---|---|---|
| Time post-treatement (h) | M1 | M2 | M3 | M4 | M5 | **Mean** | M1 | M2 | M3 | M4 | **Mean** | M1 | M2 | M3 | M4 | **Mean** |
| 0h | 1.31% | 0.55% | 0.37% | 0.00% | 0.92% | 0.80% | 0.56% | 1.86% | 1.58% | 0.80% | 0.80% | 0.57% | 0.41% | 0.48% | 1.01% | 0.80% |
| 24h | 1.75% | 0.81% | 0.86% | 0.32% | | 0.94% | 0.15% | 1.33% | 0.77% | 0.40% | 0.66% | 1.00% | 0.18% | 0.16% | 0.62% | 0.49% |
| 48h | | 0.32% | 0.00% | 0.00% | | 0.11% | | 0.33% | 0.00% | 0.23% | 0.19% | | 0.00% | 0.00% | 0.24% | 0.08% |
| 72h | | | 0.00% | 0.00% | | 0.00% | | | 0.00% | 0.00% | 0.00% | | | 0.00% | 0.00% | 0.00% |
| 96h | | | | 0.00% | | 0.00% | | | | 0.00% | 0.00% | | | | 0.00% | 0.00% |

Experiment 1

Experiment 2

Experiment 3

#### Table S7: Bioluminescence scores in Primaquine- and Tafenoquine-treated NSG mice

***Untreated control***

| days after infection | Untreated controls | | | | | | | | | | | |
|---|---|---|---|---|---|---|---|---|---|---|---|---|
| | M1 | M2 | M3 | M4 | M1 | M2 | M3 | M4 | M1 | M2 | M3 | M4 |
| 0 | ND | ND | ND | ND | ND | ND | ND | ND | ND | ND | ND | ND |
| 1 | 1.21E+08 | 1.69E+08 | 1.47E+08 | 1.38E+08 | 1.68E+08 | 1.18E+08 | 1.74E+08 | 1.39E+08 | 6.98E+07 | 6.57E+07 | 5.38E+07 | 6.18E+07 |
| 2 | 1.59E+08 | 1.10E+08 | 1.46E+08 | 1.27E+08 | 9.86E+07 | 9.35E+07 | 9.60E+07 | 9.52E+07 | 5.55E+07 | 6.09E+07 | 5.85E+07 | 6.08E+07 |
| 3 | 1.51E+08 | 9.89E+07 | 9.58E+07 | 8.42E+07 | 8.71E+07 | 6.42E+07 | 8.35E+07 | 6.23E+07 | 3.56E+07 | 3.59E+07 | 4.61E+07 | 3.84E+07 |
| 4 | 8.94E+07 | 6.35E+07 | 6.44E+07 | 5.28E+07 | 5.57E+07 | 4.40E+07 | 6.45E+07 | 5.14E+07 | 2.17E+07 | 1.46E+07 | 1.65E+07 | 1.57E+07 |
| 5 | ND | ND | ND | ND | ND | ND | ND | ND | ND | ND | ND | ND |
| 6 | ND | ND | ND | ND | ND | ND | ND | ND | ND | ND | ND | ND |
| 7 | 4.40E+07 | 2.72E+07 | 4.83E+07 | 2.34E+07 | 2.79E+07 | 2.98E+07 | 2.85E+07 | 3.02E+07 | 1.17E+07 | 9.18E+06 | 1.40E+07 | 1.43E+07 |
| 8 | 3.49E+07 | 1.69E+07 | 3.08E+07 | 1.90E+07 | 2.19E+07 | 2.79E+07 | 2.04E+07 | 2.76E+07 | 8.27E+06 | 6.83E+06 | 1.02E+07 | 7.83E+06 |
| 9 | 2.85E+07 | 9.07E+06 | 2.20E+07 | 1.48E+07 | 1.31E+07 | 8.20E+06 | 1.42E+07 | 1.79E+07 | 6.03E+06 | 4.48E+06 | 6.81E+06 | 5.26E+06 |
| 10 | 1.85E+07 | 5.97E+06 | 1.78E+07 | 1.07E+07 | 1.12E+07 | 1.44E+07 | 1.43E+07 | 1.45E+07 | 2.87E+06 | 2.09E+06 | 2.94E+06 | 2.05E+06 |
| 11 | 8.98E+06 | 2.66E+06 | 1.43E+07 | 9.99E+06 | 7.27E+06 | 1.22E+07 | 7.45E+06 | 8.12E+06 | 1.15E+06 | 8.46E+05 | 2.56E+06 | 1.62E+06 |
| 12 | ND | ND | ND | ND | ND | ND | ND | ND | ND | ND | ND | ND |
| 13 | ND | ND | ND | ND | ND | ND | ND | ND | ND | ND | ND | ND |
| 14 | 2.48E+06 | 8.04E+05 | 3.22E+06 | 1.46E+06 | 2.16E+06 | 3.12E+06 | 2.40E+06 | 2.45E+06 | ND | ND | ND | ND |
| 15 | ND | ND | ND | ND | ND | ND | ND | ND | ND | ND | ND | ND |
| 16 | 1.34E+06 | 2.72E+05 | 1.62E+06 | 8.88E+05 | 2.10E+06 | 2.00E+06 | 1.94E+06 | 1.32E+06 | ND | ND | ND | ND |

#### Table S7: Bioluminescence scores in Primaquine- and Tafenoquine-treated NSG mice (continued)

***Primaquine***

| Days after infection | 1x50mg/kg | | | | 1x30 mg/kg | | 1x20 mg/kg | | 1x10 mg/kg | |
|---|---|---|---|---|---|---|---|---|---|---|
| | M1 | M2 | M1 | M2 | M1 | M2 | M1 | M2 | M1 | M2 |
| 0 | ND | ND | ND | ND | ND | ND | ND | ND | ND | ND |
| 1 | 1.15E+08 | 9.84E+07 | 1.53E+08 | 1.52E+08 | 7.61E+07 | 9.37E+07 | 1.54E+08 | 1.48E+08 | 8.60E+07 | 1.18E+08 |
| 2 | 1.85E+07 | 2.53E+07 | 5.41E+07 | 4.22E+07 | 2.23E+07 | 2.57E+07 | 7.13E+07 | 6.13E+07 | 3.12E+07 | 3.56E+07 |
| 3 | 3.82E+06 | 4.25E+06 | 4.95E+06 | 6.69E+06 | 4.48E+06 | 6.44E+06 | 1.32E+07 | 1.78E+07 | 1.20E+07 | 1.85E+07 |
| 4 | 7.05E+05 | 7.19E+05 | 1.54E+06 | 2.03E+06 | 1.20E+06 | 1.50E+06 | 6.61E+06 | 9.96E+06 | 5.39E+06 | 5.28E+06 |
| 5 | ND | ND | ND | ND | ND | ND | ND | ND | ND | ND |
| 6 | ND | ND | ND | ND | ND | ND | ND | ND | ND | ND |
| 7 | 4.24E+05 | 3.96E+05 | 3.42E+05 | 9.96E+05 | 4.59E+05 | 6.37E+05 | 1.62E+06 | 3.44E+06 | 3.17E+06 | 2.50E+06 |
| 8 | 3.85E+05 | 5.68E+05 | 1.08E+06 | 1.22E+06 | 5.59E+05 | 6.29E+05 | 1.53E+06 | 1.94E+06 | 2.32E+06 | 1.78E+06 |
| 9 | 8.38E+05 | 7.14E+05 | 1.23E+06 | 1.25E+06 | 7.78E+05 | 8.61E+05 | 1.40E+06 | 2.11E+06 | 1.82E+06 | 1.72E+06 |
| 10 | 7.95E+05 | 9.16E+05 | 1.54E+06 | 1.57E+06 | 7.09E+05 | 6.14E+05 | 1.58E+06 | 2.01E+06 | 8.68E+05 | 9.21E+05 |
| 11 | 3.09E+05 | 3.35E+05 | 1.24E+06 | 1.13E+06 | 4.34E+05 | 3.55E+05 | 1.45E+06 | 1.72E+06 | 5.32E+05 | 4.50E+05 |
| 12 | ND | ND | ND | ND | ND | ND | ND | ND | ND | ND |
| 13 | ND | ND | ND | ND | ND | ND | ND | ND | ND | ND |
| 14 | ND | ND | 7.77E+05 | 1.11E+06 | ND | ND | 1.49E+06 | 2.03E+06 | ND | ND |
| 15 | ND | ND | ND | ND | ND | ND | ND | ND | ND | ND |
| 16 | ND | ND | 1.56E+06 | 1.61E+06 | ND | ND | 1.39E+06 | 2.02E+06 | ND | ND |

| Days after infection | 1x5 mg/kg | | 1x3 mg/kg | | 1x1 mg/kg | | 1x0.3 mg/kg | |
|---|---|---|---|---|---|---|---|---|
| | M1 | M2 | M1 | M2 | M1 | M2 | M1 | M2 |
| 0 | ND | ND | ND | ND | ND | ND | ND | ND |
| 1 | 1.05E+08 | 8.20E+07 | 1.19E+08 | 8.75E+07 | 9.77E+07 | 9.52E+07 | 8.08E+07 | 8.41E+07 |
| 2 | 5.17E+07 | 5.52E+07 | 4.58E+07 | 5.45E+07 | 5.12E+07 | 5.34E+07 | 4.93E+07 | 5.78E+07 |
| 3 | 3.91E+07 | 4.61E+07 | 3.35E+07 | 3.02E+07 | 3.95E+07 | 2.84E+07 | 3.70E+07 | 3.58E+07 |
| 4 | 1.20E+07 | 1.40E+07 | 1.33E+07 | 1.57E+07 | 1.72E+07 | 1.40E+07 | 1.77E+07 | 1.81E+07 |
| 5 | ND | ND | ND | ND | ND | ND | ND | ND |
| 6 | ND | ND | ND | ND | ND | ND | ND | ND |
| 7 | 7.37E+06 | 8.14E+06 | 1.05E+07 | 1.12E+07 | 1.65E+07 | 7.53E+06 | 1.42E+07 | 8.19E+06 |
| 8 | 5.04E+06 | 5.90E+06 | 6.45E+06 | 7.56E+06 | 1.26E+07 | 5.15E+06 | 7.56E+06 | 5.79E+06 |
| 9 | 3.38E+06 | 3.89E+06 | 4.38E+06 | 3.61E+06 | 8.31E+06 | 4.55E+06 | 5.20E+06 | 3.80E+06 |
| 10 | 1.51E+06 | 1.95E+06 | 2.85E+06 | 3.28E+06 | 5.30E+06 | 2.67E+06 | 2.55E+06 | 2.31E+06 |
| 11 | 6.74E+05 | 9.51E+05 | 1.20E+06 | 1.63E+06 | 2.30E+06 | 1.17E+06 | 1.41E+06 | 1.13E+06 |
| 12 | ND | ND | ND | ND | ND | ND | ND | ND |
| 13 | ND | ND | ND | ND | ND | ND | ND | ND |
| 14 | ND | ND | ND | ND | ND | ND | ND | ND |
| 15 | ND | ND | ND | ND | ND | ND | ND | ND |
| 16 | ND | ND | ND | ND | ND | ND | ND | ND |

#### Table S7: Bioluminescence scores in Primaquine- and Tafenoquine-treated NSG mice (continued)

***Tafenoquine***

| Days after infection | 1x100 mg/kg | | 1x70 mg/kg | | 1x50 mg/kg | | 1x20 mg/kg | | 1x5 mg/kg | |
|---|---|---|---|---|---|---|---|---|---|---|
| | M1 | M2 | M1 | M2 | M1 | M2 | M1 | M2 | M1 | M2 |
| 0 | ND | ND | ND | ND | ND | ND | ND | ND | ND | ND |
| 1 | 7.64E+07 | 1.24E+08 | 2.23E+08 | 1.51E+08 | 1.99E+08 | 1.79E+08 | 1.70E+08 | 1.55E+08 | 1.30E+08 | 1.98E+08 |
| 2 | 2.74E+07 | 7.41E+07 | 5.83E+07 | 6.30E+07 | 1.26E+08 | 1.07E+08 | 8.54E+07 | 7.30E+07 | 8.06E+07 | 1.07E+08 |
| 3 | 6.53E+06 | 1.03E+07 | 8.33E+06 | 1.06E+07 | 1.82E+07 | 1.42E+07 | 2.85E+07 | 3.49E+07 | 5.57E+07 | 7.20E+07 |
| 4 | 1.93E+06 | 1.86E+06 | 2.23E+06 | 2.20E+06 | 3.28E+06 | 3.24E+06 | 3.11E+06 | 4.91E+06 | 4.57E+07 | 5.52E+07 |
| 5 | ND | ND | ND | ND | ND | ND | ND | ND | ND | ND |
| 6 | ND | ND | ND | ND | ND | ND | ND | ND | ND | ND |
| 7 | 9.69E+05 | 7.90E+05 | 9.06E+05 | 9.72E+05 | 8.04E+05 | 7.37E+05 | 1.04E+06 | 1.07E+06 | 4.80E+06 | 5.83E+06 |
| 8 | 9.09E+05 | 7.80E+05 | 8.07E+05 | 8.97E+05 | 3.69E+06 | 3.37E+06 | 9.38E+05 | 1.14E+06 | 2.52E+06 | 2.98E+06 |
| 9 | 9.66E+05 | 9.10E+05 | 6.15E+05 | 9.55E+05 | 6.53E+05 | 5.97E+05 | 1.13E+06 | 1.01E+06 | 1.89E+06 | 1.96E+06 |
| 10 | 9.22E+05 | 1.02E+06 | 1.08E+06 | 1.09E+06 | 1.15E+06 | 1.01E+06 | 1.59E+06 | 1.42E+06 | 1.98E+06 | 2.31E+06 |
| 11 | 9.04E+05 | 8.16E+05 | 1.13E+06 | 1.26E+06 | 8.28E+05 | 8.27E+05 | 1.41E+06 | 1.65E+06 | 1.61E+06 | 2.00E+06 |
| 12 | ND | ND | ND | ND | ND | ND | ND | ND | ND | ND |
| 13 | ND | ND | ND | ND | ND | ND | ND | ND | ND | ND |
| 14 | 6.21E+05 | 7.66E+05 | 1.04E+06 | 1.22E+06 | ND | ND | 1.29E+06 | 1.34E+06 | 1.14E+06 | 1.33E+06 |
| 15 | ND | ND | ND | ND | ND | ND | ND | ND | ND | ND |
| 16 | 9.00E+05 | 1.04E+06 | 1.27E+06 | 1.19E+06 | ND | ND | 1.71E+06 | 1.61E+06 | 1.28E+06 | 1.08E+06 |

Values are Total Flux [p/s] per mouse (4cmX15cm ROI: Em Filter=Open , Ex Filter=Block, Bin:(M)4, FOV:24, f1.2, 180s)

m: mouse

ND: not determined

Experiment 1

Experiment 2

Experiment 3

Background calculation: Mean+ 3*SD of days11-16 primaquine 1x 50 mg/kg

days11-16 primaquine 1x 50 mg/kg:

| | |
|---|---|
| mean: | 1.24E+06 |
| SD: | 282904.6 |
| **BG** | **1.80E+06** |

#### Table S8: Gametocyte count in Primaquine- and Tafenoquine-treated NSG mice

***Untreated control***

| Days after infection | Untreated controls | | | | | | | | | | | |
|---|---|---|---|---|---|---|---|---|---|---|---|---|
| | M1 | M2 | M3 | M4 | M1 | M2 | M3 | M4 | M1 | M2 | M3 | M4 |
| 0 | 0.97 | 1.18 | 1.03 | 1.09 | 0.75 | 0.65 | 0.79 | 0.78 | 0.58 | 0.61 | 0.54 | 0.46 |
| 1 | 0.78 | 0.8 | 0.71 | 0.69 | 0.61 | 0.61 | 0.59 | 0.57 | 0.56 | 0.54 | 0.49 | 0.41 |
| 2 | 0.63 | 0.66 | 0.59 | 0.61 | 0.54 | 0.46 | 0.55 | 0.47 | 0.42 | 0.37 | 0.41 | 0.35 |
| 3 | 0.53 | 0.51 | 0.49 | 0.5 | 0.23 | 0.36 | 0.38 | 0.4 | 0.36 | 0.35 | 0.29 | 0.34 |
| 4 | 0.49 | 0.45 | 0.47 | 0.4 | 0.23 | 0.33 | 0.28 | 0.26 | 0.27 | 0.18 | 0.27 | 0.21 |
| 5 | ND | ND | ND | ND | ND | ND | ND | ND | ND | ND | ND | ND |
| 6 | ND | ND | ND | ND | ND | ND | ND | ND | ND | ND | ND | ND |
| 7 | 0.18 | 0.18 | 0.17 | 0.14 | 0.18 | 0.18 | 0.17 | 0.14 | 0.15 | 0.11 | 0.15 | 0.08 |
| 8 | 0.21 | 0.07 | 0.12 | 0.17 | 0.17 | 0.2 | 0.12 | 0.16 | 0.08 | 0.07 | 0.11 | 0.06 |
| 9 | 0.2 | 0.06 | 0.16 | 0.15 | 0.09 | 0.14 | 0.08 | 0.06 | 0.08 | 0.03 | 0.05 | 0.05 |
| 10 | 0.12 | 0.03 | 0.08 | 0.06 | 0.05 | 0.12 | 0.08 | 0.06 | ND | ND | ND | ND |
| 11 | 0.07 | 0.02 | 0.06 | 0.04 | 0.04 | 0.02 | 0.07 | 0.06 | ND | ND | ND | ND |
| 12 | 0.06 | 0.01 | 0.05 | 0.04 | ND | ND | ND | ND | ND | ND | ND | ND |
| 13 | ND | ND | ND | ND | ND | ND | ND | ND | ND | ND | ND | ND |
| 14 | 0.01 | 0.001 | 0.01 | 0.01 | 0.01 | 0.01 | 0.01 | 0.01 | ND | ND | ND | ND |
| 15 | ND | ND | ND | ND | ND | ND | ND | ND | ND | ND | ND | ND |
| 16 | 0.001 | 0.001 | 0.001 | 0.001 | 0.001 | 0.001 | 0.001 | 0.001 | ND | ND | ND | ND |

#### Table S8: Gametocyte count in Primaquine- and Tafenoquine-treated NSG mice (continued)

***Primaquine***

| Days after infection | 1x50mg/kg | | | | 1x30 mg/kg | | 1x20 mg/kg | | 1x10 mg/kg | |
|---|---|---|---|---|---|---|---|---|---|---|
| | M1 | M2 | M1 | M2 | M1 | M2 | M1 | M2 | M1 | M2 |
| 0 | 0.63 | 0.57 | 0.76 | 0.69 | 0.57 | 0.51 | 0.76 | 0.68 | 0.52 | 0.59 |
| 1 | 0.58 | 0.49 | 0.63 | 0.58 | 0.41 | 0.46 | 0.61 | 0.63 | 0.47 | 0.45 |
| 2 | 0.34 | 0.29 | 0.42 | 0.41 | 0.29 | 0.32 | 0.35 | 0.42 | 0.36 | 0.29 |
| 3 | 0.05 | 0.11 | 0.07 | 0.15 | 0.1 | 0.14 | 0.18 | 0.24 | 0.24 | 0.2 |
| 4 | 0.001 | 0.01 | 0.01 | 0.01 | 0.001 | 0.01 | 0.05 | 0.1 | 0.02 | 0.04 |
| 5 | ND | ND | ND | ND | ND | ND | ND | ND | ND | ND |
| 6 | ND | ND | ND | ND | ND | ND | ND | ND | ND | ND |
| 7 | 0.001 | 0.01 | 0.001 | 0.001 | 0.001 | 0.001 | 0.001 | 0.01 | 0.02 | 0.03 |
| 8 | 0.001 | 0.001 | 0.001 | 0.001 | 0.001 | 0.001 | 0.01 | 0.01 | 0.02 | 0.01 |
| 9 | 0.001 | 0.001 | 0.001 | 0.001 | 0.001 | 0.001 | 0.01 | 0.01 | 0.02 | 0.001 |
| 10 | ND | ND | 0.001 | 0.001 | ND | ND | 0.001 | 0.001 | ND | ND |
| 11 | ND | ND | 0.001 | 0.001 | ND | ND | 0.01 | 0.01 | ND | ND |
| 12 | ND | ND | ND | ND | ND | ND | ND | ND | ND | ND |
| 13 | ND | ND | ND | ND | ND | ND | ND | ND | ND | ND |
| 14 | ND | ND | 0.001 | 0.001 | ND | ND | 0.001 | 0.001 | ND | ND |
| 15 | ND | ND | ND | ND | ND | ND | ND | ND | ND | ND |
| 16 | ND | ND | 0.001 | 0.001 | ND | ND | 0.001 | 0.001 | ND | ND |

| Days after infection | 1x5 mg/kg | | 1x3 mg/kg | | 1x1 mg/kg | | 1x0.3 mg/kg | |
|---|---|---|---|---|---|---|---|---|
| | M1 | M2 | M1 | M2 | M1 | M2 | M1 | M2 |
| 0 | 0.56 | 0.47 | 0.46 | 0.61 | 0.56 | 0.49 | 0.57 | 0.54 |
| 1 | 0.43 | 0.41 | 0.44 | 0.5 | 0.43 | 0.41 | 0.49 | 0.5 |
| 2 | 0.32 | 0.34 | 0.37 | 0.34 | 0.36 | 0.39 | 0.37 | 0.37 |
| 3 | 0.31 | 0.25 | 0.3 | 0.33 | 0.29 | 0.3 | 0.34 | 0.29 |
| 4 | 0.14 | 0.08 | 0.11 | 0.2 | 0.11 | 0.12 | 0.18 | 0.19 |
| 5 | ND | ND | ND | ND | ND | ND | ND | ND |
| 6 | ND | ND | ND | ND | ND | ND | ND | ND |
| 7 | 0.1 | 0.07 | 0.07 | 0.13 | 0.1 | 0.04 | 0.08 | 0.05 |
| 8 | 0.03 | 0.03 | 0.06 | 0.07 | 0.08 | 0.02 | 0.05 | 0.04 |
| 9 | 0.01 | 0.03 | 0.01 | 0.06 | 0.04 | 0.05 | 0.04 | 0.03 |
| 10 | ND | ND | ND | ND | ND | ND | ND | ND |
| 11 | ND | ND | ND | ND | ND | ND | ND | ND |
| 12 | ND | ND | ND | ND | ND | ND | ND | ND |
| 13 | ND | ND | ND | ND | ND | ND | ND | ND |
| 14 | ND | ND | ND | ND | ND | ND | ND | ND |
| 15 | ND | ND | ND | ND | ND | ND | ND | ND |
| 16 | ND | ND | ND | ND | ND | ND | ND | ND |

#### Table S8: Gametocyte count in Primaquine- and Tafenoquine-treated NSG mice (continued)

***Tafenoquine***

| Days after infection | 1x100 mg/kg | | 1x70 mg/kg | | 1x50 mg/kg | | 1x20 mg/kg | | 1x5 mg/kg | |
|---|---|---|---|---|---|---|---|---|---|---|
| | M1 | M2 | M1 | M2 | M1 | M2 | M1 | M2 | M1 | M2 |
| 0 | 0.65 | 0.83 | 0.75 | 0.76 | 1.17 | 1.18 | 0.73 | 0.66 | 0.64 | 0.69 |
| 1 | 0.56 | 0.62 | 0.61 | 0.58 | 0.84 | 0.68 | 0.58 | 0.59 | 0.62 | 0.52 |
| 2 | 0.45 | 0.58 | 0.48 | 0.46 | 0.57 | 0.68 | 0.42 | 0.43 | 0.38 | 0.49 |
| 3 | 0.04 | 0.12 | 0.21 | 0.13 | 0.33 | 0.19 | 0.22 | 0.26 | 0.3 | 0.34 |
| 4 | 0.01 | 0 | 0.01 | 0.04 | 0.03 | 0.02 | 0.03 | 0.05 | 0.28 | 0.27 |
| 5 | ND | ND | ND | ND | ND | ND | ND | ND | ND | ND |
| 6 | ND | ND | ND | ND | ND | ND | ND | ND | ND | ND |
| 7 | 0.001 | 0.001 | 0.001 | 0.001 | 0.001 | 0.001 | 0.01 | 0.001 | 0.04 | 0.03 |
| 8 | 0.001 | 0.001 | 0.001 | 0.001 | 0.001 | 0.001 | 0.01 | 0.001 | 0.01 | 0.02 |
| 9 | 0.001 | 0.001 | 0.001 | 0.001 | 0.001 | 0.001 | 0.001 | 0.001 | 0.01 | 0.02 |
| 10 | 0.001 | 0.001 | 0.001 | 0.001 | 0.001 | 0.001 | 0.001 | 0.001 | 0.01 | 0.01 |
| 11 | 0.001 | 0.001 | 0.001 | 0.001 | 0.001 | 0.001 | 0.001 | 0.001 | 0.01 | 0.01 |
| 12 | ND | ND | ND | ND | ND | ND | ND | ND | ND | ND |
| 13 | ND | ND | ND | ND | ND | ND | ND | ND | ND | ND |
| 14 | 0.001 | 0.001 | 0.001 | 0.001 | 0.001 | 0.001 | 0.001 | 0.001 | 0.001 | 0.001 |
| 15 | ND | ND | ND | ND | ND | ND | ND | ND | ND | ND |
| 16 | 0.001 | 0.001 | 0.001 | 0.001 | ND | ND | 0.001 | 0.001 | 0.001 | 0.001 |

All values are % gametocytemia
Values of 0.001 represent no detectable gametocytes (negative slides reading) and are only used for plotting with a logarithmic scale

ND: not determined

Experiment 1
Experiment 2
Experiment 3

**Table S9: Pharmacokinetic profiles of Primaquine and Tafenoquine in the NSG model.** ND: not detected (below the quantification level).

#### *Primaquine*

***Exposure levels***

| Time post-treatement (h) | 50 mg/kg | | | | | 30 mg/kg | | | | |
|---|---|---|---|---|---|---|---|---|---|---|
| | M1 (ng/ml) | M2 (ng/ml) | **Mean (ng/ml)** | **SD (ng/ml)** | **CV (%)** | M1 (ng/ml) | M2 (ng/ml) | **Mean (ng/ml)** | **SD (ng/ml)** | **CV (%)** |
| 2 | 80.2 | 119 | **99.6** | **27.4** | **27.5** | 74.4 | 32 | **53.2** | **30.0** | **56.4** |
| 4 | 77.6 | 56.2 | **66.9** | **15.1** | **22.6** | 43.6 | 33.8 | **38.7** | **6.9** | **17.9** |
| 24 | 2.14 | 2.22 | **2.2** | **0.1** | **2.6** | 1.72 | < 1.15 | **1.7** | **ND** | **ND** |

***Pharmacokinetic parameters***

| | | | | | | | | | | |
|---|---|---|---|---|---|---|---|---|---|---|
| **t_max (h)** | 2.0 | 2.0 | **2.0** | **0.0** | **0.0** | 2.0 | 2.0 | **2.0** | **0.0** | **0.0** |
| **C_max (ng/ml)** | 80 | 119 | **99.6** | **27.4** | **27.5** | 74 | 34 | **54.1** | **28.7** | **53.1** |
| **C_max/dose ((ng/ml)/(mg/kg))** | 1.6 | 2.4 | **2.0** | **0.6** | **27.7** | 2.5 | 1.1 | **1.8** | **1.0** | **52.9** |
| **AUC_all (h*ng/ml)** | 658 | 621 | **639.5** | **26.2** | **4.1** | 449 | 436 | **442.5** | **9.2** | **2.1** |
| **AUC_all/dose ((h*ng/ml)/(mg/kg))** | 13 | 12 | **12.8** | **0.6** | **4.4** | 15 | 15 | **14.8** | **0.4** | **2.4** |

***Exposure levels***

| Time post-treatement (h) | 10 mg/kg | | | | | 5 mg/kg | | | | |
|---|---|---|---|---|---|---|---|---|---|---|
| | M1 (ng/ml) | M2 (ng/ml) | **Mean (ng/ml)** | **SD (ng/ml)** | **CV (%)** | M1 (ng/ml) | M2 (ng/ml) | **Mean (ng/ml)** | **SD (ng/ml)** | **CV (%)** |
| 2 | 12.5 | 13.9 | **13.2** | **1.0** | **7.5** | 6.66 | 6.18 | **6.4** | **0.3** | **5.3** |
| 4 | 6.82 | 9.7 | **8.3** | **2.0** | **24.7** | 1.15 | 1.39 | **1.3** | **0.2** | **13.4** |
| 24 | <1.15 | <1.15 | **ND** | **ND** | **ND** | <1.15 | <1.15 | **ND** | **ND** | **ND** |

***Pharmacokinetic parameters***

| | | | | | | | | | | |
|---|---|---|---|---|---|---|---|---|---|---|
| **t_max (h)** | 2.0 | 2.0 | **2.0** | **0.0** | **0.0** | 2.0 | 2.0 | **2.0** | **0.0** | **0.0** |
| **C_max (ng/ml)** | 13 | 14 | **13.2** | **1.0** | **7.5** | 6.7 | 6.2 | **6.4** | **0.3** | **5.3** |
| **C_max/dose ((ng/ml)/(mg/kg))** | 1.3 | 1.4 | **1.3** | **0.1** | **7.5** | 1.3 | 1.2 | **1.3** | **0.1** | **5.0** |
| **AUC_all (h*ng/ml)** | 99 | 134 | **116.7** | **24.5** | **21.0** | 24 | 27 | **25.5** | **1.5** | **5.8** |
| **AUC_all/dose ((h*ng/ml)/(mg/kg))** | 10 | 13 | **11.7** | **2.4** | **21.0** | 4.9 | 5.3 | **5.1** | **0.3** | **5.7** |

**Table S9: Pharmacokinetic profiles of Primaquine and Tafenoquine in the NSG model (continued).** ND: not detected (below the quantification level).

***Exposure levels***

| Time post-treatement (h) | 3 mg/kg | | | | | 1 mg/kg | | | | |
|---|---|---|---|---|---|---|---|---|---|---|
| | M1 (ng/ml) | M2 (ng/ml) | **Mean (ng/ml)** | **SD (ng/ml)** | **CV (%)** | M1 (ng/ml) | M2 (ng/ml) | **Mean (ng/ml)** | **SD (ng/ml)** | **CV (%)** |
| 2 | 6.54 | 7.16 | **6.9** | **0.4** | **6.4** | 1.86 | <1.15 | **1.9** | **ND** | **ND** |
| 4 | 6.36 | 1.16 | **3.8** | **3.7** | **97.8** | <1.15 | <1.15 | **ND** | **ND** | **ND** |
| 24 | <1.15 | <1.15 | **ND** | **ND** | **ND** | <1.15 | <1.15 | **ND** | **ND** | **ND** |

***Pharmacokinetic parameters***

| | | | | | | | | | | |
|---|---|---|---|---|---|---|---|---|---|---|
| **t_max (h)** | 2.0 | 2.0 | **2.0** | **0.0** | **0.0** | ND | ND | **ND** | **ND** | **ND** |
| **C_max (ng/ml)** | 6.5 | 7.2 | **6.9** | **0.4** | **6.4** | ND | ND | **ND** | **ND** | **ND** |
| **C_max/dose ((ng/ml)/(mg/kg))** | 2.2 | 2.4 | **2.3** | **0.1** | **6.5** | ND | ND | **ND** | **ND** | **ND** |
| **AUC_all (h*ng/ml)** | 83 | 25 | **54.2** | **40.7** | **75.1** | ND | ND | **ND** | **ND** | **ND** |
| **AUC_all/dose ((h*ng/ml)/(mg/kg))** | 38 | 8.5 | **23.1** | **20.7** | **89.6** | ND | ND | **ND** | **ND** | **ND** |

***Exposure levels***

| Time post-treatement (h) | 0.3 mg/kg | | | | |
|---|---|---|---|---|---|
| | M1 (ng/ml) | M2 (ng/ml) | **Mean (ng/ml)** | **SD (ng/ml)** | **CV (%)** |
| 2 | <1.15 | <1.15 | **ND** | **ND** | **ND** |
| 4 | <1.15 | <1.15 | **ND** | **ND** | **ND** |
| 24 | <1.15 | <1.15 | **ND** | **ND** | **ND** |

***Pharmacokinetic parameters***

| | | | | | |
|---|---|---|---|---|---|
| **t_max (h)** | ND | ND | **ND** | **ND** | **ND** |
| **C_max (ng/ml)** | ND | ND | **ND** | **ND** | **ND** |
| **C_max/dose ((ng/ml)/(mg/kg))** | ND | ND | **ND** | **ND** | **ND** |
| **AUC_all (h*ng/ml)** | ND | ND | **ND** | **ND** | **ND** |
| **AUC_all/dose ((h*ng/ml)/(mg/kg))** | ND | ND | **ND** | **ND** | **ND** |

**Table S9: Pharmacokinetic profiles of Primaquine and Tafenoquine in the NSG model (continued).** ND: not detected (below the quantification level).

#### *Tafenoquine*

***Exposure levels***

| Time post-treatement (h) | 100 mg/kg | | | | | 70 mg/kg | | | | |
|---|---|---|---|---|---|---|---|---|---|---|
| | M1 (ng/ml) | M2 (ng/ml) | **Mean (ng/ml)** | **SD (ng/ml)** | **CV (%)** | M1 (ng/ml) | M2 (ng/ml) | **Mean (ng/ml)** | **SD (ng/ml)** | **CV (%)** |
| 2 | 762 | 1070 | **916.0** | **217.8** | **23.8** | 778 | 582 | **680.0** | **138.6** | **20.4** |
| 4 | 1590 | 1340 | **1465.0** | **176.8** | **12.1** | 1020 | 686 | **853.0** | **236.2** | **27.7** |
| 24 | 1720 | 1890 | **1805.0** | **120.2** | **6.7** | 892 | 850 | **871.0** | **29.7** | **3.4** |
| 48 | 2080 | 1300 | **1690.0** | **551.5** | **32.6** | 622 | 680 | **651.0** | **41.0** | **6.3** |
| 72 | 1070 | 1090 | **1080.0** | **14.1** | **1.3** | 596 | 396 | **496.0** | **141.4** | **28.5** |
| 120 | 412 | 422 | **417.0** | **7.1** | **1.7** | 282 | 318 | **300.0** | **25.5** | **8.5** |

***Pharmacokinetic parameters***

| | | | | | | | | | | |
|---|---|---|---|---|---|---|---|---|---|---|
| **t_max (h)** | 48.0 | 24.0 | **36.0** | **17.0** | **47.1** | 4.0 | 24.0 | **14.0** | **14.1** | **101.0** |
| **C_max (ng/ml)** | 2080 | 1890 | **1985.0** | **134.4** | **6.8** | 1020 | 850 | **935.0** | **120.2** | **12.9** |
| **C_max/dose ((ng/ml)/(mg/kg))** | 20.8 | 18.9 | **19.9** | **1.3** | **6.8** | 14.6 | 12.1 | **13.4** | **1.8** | **13.2** |
| **AUC_all (h*ng/ml)** | 151000 | 136000 | **143500.0** | **10606.6** | **7.4** | 74400 | 65200 | **69800.0** | **6505.4** | **9.3** |
| **AUC_all/dose ((h*ng/ml)/(mg/kg))** | 1510 | 1360 | **1435.0** | **106.1** | **7.4** | 1060 | 931 | **995.5** | **91.2** | **9.2** |

***Exposure levels***

| Time post-treatement (h) | 20 mg/kg | | | | | 5 mg/kg | | | | |
|---|---|---|---|---|---|---|---|---|---|---|
| | M1 (ng/ml) | M2 (ng/ml) | **Mean (ng/ml)** | **SD (ng/ml)** | **CV (%)** | M1 (ng/ml) | M2 (ng/ml) | **Mean (ng/ml)** | **SD (ng/ml)** | **CV (%)** |
| 2 | 220 | 330 | **275.0** | **77.8** | **28.3** | 37.2 | 31 | **34.1** | **4.4** | **12.9** |
| 4 | 400 | 358 | **379.0** | **29.7** | **7.8** | 52.4 | 55.2 | **53.8** | **2.0** | **3.7** |
| 24 | 348 | 514 | **431.0** | **117.4** | **27.2** | 94.66 | 114 | **104.3** | **13.7** | **13.1** |
| 48 | 228 | 270 | **249.0** | **29.7** | **11.9** | 41.6 | 55.8 | **48.7** | **10.0** | **20.6** |
| 72 | 197 | 242 | **219.5** | **31.8** | **14.5** | 42.6 | 43.8 | **43.2** | **0.8** | **2.0** |
| 120 | 101 | 94 | **97.5** | **4.9** | **5.1** | 19.4 | 19 | **19.2** | **0.3** | **1.5** |

***Pharmacokinetic parameters***

| | | | | | | | | | | |
|---|---|---|---|---|---|---|---|---|---|---|
| **t_max (h)** | 4.0 | 24.0 | **14.0** | **14.1** | **101.0** | 24.0 | 24.0 | **24.0** | **0.0** | **0.0** |
| **C_max (ng/ml)** | 400 | 514 | **457.0** | **80.6** | **17.6** | 94.6 | 114.0 | **104.3** | **13.7** | **13.2** |
| **C_max/dose ((ng/ml)/(mg/kg))** | 20.0 | 25.7 | **22.9** | **4.0** | **17.6** | 18.9 | 22.8 | **20.9** | **2.8** | **13.2** |
| **AUC_all (h*ng/ml)** | 27100 | 32500 | **29800.0** | **3818.4** | **12.8** | 5570 | 6380 | **5975.0** | **572.8** | **9.6** |
| **AUC_all/dose ((h*ng/ml)/(mg/kg))** | 1360 | 1620 | **1490.0** | **183.8** | **12.3** | 1110.0 | 1280.0 | **1195.0** | **120.2** | **10.1** |
